# A factor-based analysis of individual human microglia uncovers regulators of an Alzheimer-related transcriptional signature

**DOI:** 10.1101/2025.03.27.641500

**Authors:** Victoria S. Marshe, John F. Tuddenham, Kevin Chen, Rebecca Chiu, Verena C. Haage, Yiyi Ma, Annie J. Lee, Neil A. Shneider, Julian P. Agin-Liebes, Roy N. Alcalay, Andrew F. Teich, Peter Canoll, Claire S. Riley, Dirk Keene, Julie A. Schneider, David A. Bennett, Vilas Menon, Mariko Taga, Hans-Ulrich Klein, Marta Olah, Masashi Fujita, Ya Zhang, Peter A. Sims, Philip L. De Jager

## Abstract

Human microglial heterogeneity is only beginning to be appreciated at the molecular level. Here, we present a large, single-cell atlas of expression signatures from 441,088 live microglia broadly sampled across a diverse set of brain regions and neurodegenerative and neuroinflammatory diseases obtained from 161 donors sampled at autopsy or during a neurosurgical procedure. Using single-cell hierarchical Poisson factorization (scHPF), we derived a 23-factor model for continuous gene expression signatures across microglia which capture specific biological processes (e.g., metabolism, phagocytosis, antigen presentation, inflammatory signaling, disease-associated states). Using external datasets, we evaluated the aspects of microglial phenotypes that are encapsulated in various *in vitro* and *in vivo* microglia models and identified and replicated the role of two factors in human postmortem tissue of Alzheimer’s disease (AD). Further, we derived a complex network of transcriptional regulators for all factors, including regulators of an AD-related factor enriched for the mouse disease-associated microglia 2 (DAM2) signature: *ARID5B*, *CEBPA*, *MITF*, and *PPARG*. We replicated the role of these four regulators in the AD-related factor and then designed a multiplexed MERFISH panel to assess our microglial factors using spatial transcriptomics. We find that, unlike cells with high expression of the interferon-response factor, cells with high expression of the AD DAM2-like factor are widely distributed in neocortical tissue. We thus propose a novel analytic framework that provides a taxonomic approach for microglia that is more biologically interpretable and use it to uncover new therapeutic targets for AD.

## Introduction

As one class of tissue-resident macrophages in the central nervous system (CNS), microglia play a key role in development, homeostasis, immunity, and neuropathological processes.^1^ Microglia have a unique ontogeny, deriving from a developmentally distinct lineage compared to other CNS myeloid cells.^2^ Over the past decade, microglia, which constitute 0.5-10% of brain cells depending on the brain region,^3^ have been extensively characterized phenotypically and functionally, particularly in murine models.^1^ However, microglial heterogeneity in humans is only beginning to be appreciated at the molecular level, with evidence highlighting the context-dependent and tissue-specific transcriptional signatures and regulatory networks of among human-derived microglia compared to those derived from mice.^4,5^ Recent advances in single-cell RNA sequencing (scRNA-seq) of live human postmortem microglia have provided insights on aging and disease-associated signatures as compared to single-nucleus and bulk RNA sequencing which may not sufficiently capture activated microglial states at proper resolution and complexity.^6,7^

Despite the challenges of defining microglial subtypes due to the complexity of their likely continuous transcriptomic states,^8^ studies have shown that a disease-associated microglia (DAM) subtype may play a role in murine amyloid proteinopathy models and other neurodegenerative disease models.^9^ The murine DAM signature is enriched in a minority of human microglia, some of which are implicated in disease; the equivalent signature remains to be better defined in humans but has been described in several large single cell or single-nucleus studies^10–16^. Recent efforts to broadly characterize microglial heterogeneity across neurodegenerative diseases point to key processes including metabolism, antigen presentation, motility, and proliferation as axes of polarization across different subtypes.^16^ Functional interrogation of specific microglial subtypes which show enrichment of susceptibility genes for neurodegenerative diseases on one hand, or the DAM signature on the other, may provide therapeutic insights.^17,18^ However, these studies may be limited because of their clustering-based approach to model a complex distribution of individual microglia across a multiplicity of transcriptional gradients: hard boundaries assigning each microglia to a given subtype are not satisfactory for modeling as cells may lie on a continuum of transcriptional and functional states.

Given the need to further enhance and refine the modeling of human microglia across disease categories, we present here a large single-cell transcriptomic data resource derived from 441,088 live microglia broadly sampled from 199 tissue samples across brain regions and neurodegenerative and neuroinflammatory diseases. Without partitioning microglia into ‘subtypes’, as is common with traditional clustering approaches, we used an analytic framework called single-cell hierarchical Poisson factorization (scHPF)^19,20^ to build a 23-factor model of co-expression patterns present across brain regions and neurodegenerative diseases, which will facilitate cross-disease and cross-study comparisons. We also show that the factors are broadly applicable across various human datasets (both using single-cell and single-nucleus platforms), as well as *in vitro* and *in vivo* model systems: these include the HMC3 cell line, induced pluripotent stem cell (iPSC)-derived microglia (iMG) – with and without drug treatment, biological and genetic perturbations – and a murine xenograft iMG model. Furthermore, we assess our factors for association with Alzheimer’s disease (AD)-related traits and determine a complex regulatory network which identifies key transcriptional regulators for each of the 23 factors, and we replicate the roles of these regulators in independent datasets, including regulators for an AD-associated factor. Finally, we report the design and successful deployment of a multiplex MERFISH panel for the 23 human microglial factors that determines the topographical distribution of the AD-associated microglial factor in the human neocortex and contrast it to the aggregated pattern seen with the interferon-response factor.

### A 23-factor model of microglial expression signatures

Given the lack of clearly discrete microglial subpopulations and our prior model that described a collection of different vectors of differentiation that lead to a multipolar projection of microglial states,^16^ we applied scHPF to identify *k* latent factors (see Methods) that may be used to represent continuous gene expression factors. We applied this method to an expanded live microglial **Resource** that doubles the size of the previous version of the resource^16^ and is designed to sample a broad diversity of human microglial states across diseases and across brain regions. Following pre-processing that included the removal of macrophages and monocytes, we retained scRNAseq data from 441,088 live human microglia from 161 donors and 20 unique brain tissue types in downstream analyses (**Figure 1A** and **Figure S1**). The characteristics of these donors are presented in **Table S1,** and we refer to this collection of donors to as the ‘discovery sample’. It contains our earlier data^16^ as well as 225,382 additional transcriptomes from 87 newly-sequenced donors (**Table S2**). Our final model included 23 factors (**Table S3**) (three other factors capturing technical variation were not considered in downstream analyses, see **Methods** for details). Each factor can be captured by a gene expression signature (**Figure 1B**) that can be annotated with a reference of biological pathways to facilitate biological interpretation (**Table S5**). These factors have variable levels of expression across cells (**Figure S1D**). We characterized the factors into seven broad categories based on their overlap of enriched biological processes, including T-cell activation, metabolism, antigen presentation, disease-associated spectrum, inflammation, cellular stress, and miscellaneous (**Figure S1E**).

**Figure 1.**
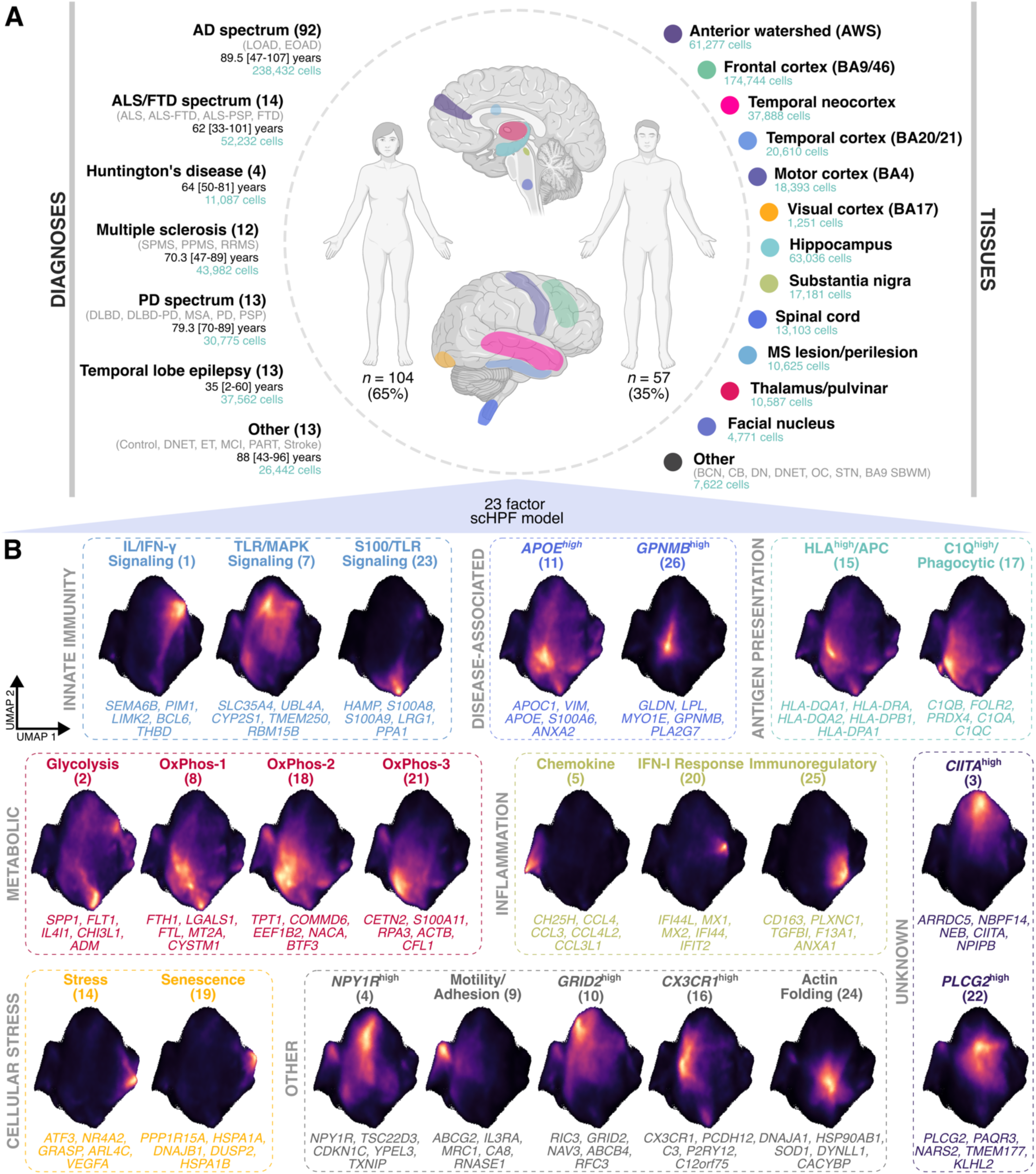
A 23-factor model characterizing microglia function in a broad single-cell dataset. **(A)** Summary of collected tissues. *Abbreviations*: BCN, body of caudate nucleus; CB, cerebellum; DN, dentate nucleus of the cerebellum; DNET, dysembryoplastic neuroepithelial tumor; OC, occipital cortex; SBWM, subcortical white matter; STN, subthalamic nucleus. **(B)** Density maps showing the UMAP space and the distribution of the 23 microglial factors at the single-cell level. Lighter color indicates a relatively greater density of cells with higher factor scores. Note, the densities are not directly comparable across factors. Showing the top five genes per factor and overarching categories.

In our model, factors captured gene signatures of homeostatic microglial functions: the *CX3CR1*^High^ factor (scHPF_16) captured the classic ‘homeostatic’ signature that includes high expression of *CX3CR1*, *P2RY12*, and *TMEM119* (**Figure S2**). Four factors are enriched with genes from the murine DAM signature.^9^ For example, the *APOE*^High^ factor (scHPF_11) included top loading for the *CTSD*, *APOE*, *TREM2* and *CD9* genes. On the other hand, the *GPNMB*^High^ factor (scHPF_26) is enriched for the murine DAM2 signature^9^ and includes high loadings for *TREM2*, *CD9* and *LPL*. Notably, the *GPNMB*^High^ factor also showed GO enrichment for pathways associated with foam cell differentiation, consistent with other reports^21^ (**Table S4**). Among the four factors capturing metabolic processes, three factors showed GO enrichment for oxidative phosphorylation (**Figure S3**). To better understand the differences between these three factors, we conducted mitochondrial functional pathway enrichment using MitoCarta3.0^22^, an inventory of genes and pathways with strong support for mitochondrial localization. Of our 23 microglial factors, the three oxidative phosphorylation factors showed the largest proportion of genes annotated to the mitochondrion (**Figure S3B**). The OxPhos-3 factor (scHPF_18) captured the largest distribution of pathways beyond oxidative phosphorylation, including protein translation, protein homeostasis, as well as lipid and nucleotide metabolism.

### Evaluating sample characteristics in relation to the factor model

Overall, this study was designed to maximize the diversity of microglia that were sampled; it was not designed to compare microglial profiles among diseases; most disease categories have too few donors at this time to capture disease-specific factors and are therefore underpowered to support comparative analyses. However, for completeness, we present the characteristics of our data across conditions of interest, including sex, age, and brain region. We used two complementary approaches to generate preliminary assessments of these variables: *Milo*^23^, a single-cell level test of differential abundance of between conditions, and donor-level linear regressions. *Milo* defines local cellular neighborhoods using a *k*-nearest neighbor (KNN) graph and allows the testing of their differential abundance while accounting for confounding variables^23^. To explore the robustness of signals, we also used donor-aggregated factor scores to explore associations adjusted for confounding variables.

We first explored whether any factors showed sex-or age-specific effects. While most donors in the discovery cohort were female, there was some variation in the proportion of females across diagnostic categories (**Figure S4**). Therefore, we restricted our analyses to donors with a diagnosis of pathological AD (our single largest donor category, *n*=92) to mitigate potential disease-specific effects. Compared to female tissue, male tissue showed a significant depletion of microglia with a high co-expression of the motility (scHPF_9) and chemokine signaling (scHPF_5) factors, as well as enrichment of microglia with a high co-expression of the *GRID2*^High^ (scHPF_24) and motility (scHPF_9) factors (**Figure S4C-G**). In the same subset of AD donors, after covarying for sex, we found that among donors aged >85 years old, there was a strong depletion of microglia highly co-expressing the chemokine and motility factors (scHPF_5, scHPF_9) (**Figure S5**); this effect became more prominent with increasing age with the co-expressed signature being almost entirely depleted in the oldest donors (>95 years, *n*=19). However, at the lower-resolution of donor-level factor scores, this association was not observed. On the other hand, donors >85 years showed an enrichment of microglia defined by high expression of the C1Q^High^/phagocytic factor (scHPF_17), and to a lesser extent oxidative phosphorylation factors (scHPF_18, scHPF_21).

We also explored potential regional differences in factor expression. In a similar targeted analysis in donors diagnosed with pathological AD, we compared deep frontal white matter (anterior watershed; AWS), temporal cortex (BA20/21), and the hippocampus to the frontal cortex (BA9/46) (**Figure S6**). Overall, there were few significant regional differences. Notably, we observed an enrichment of microglia expressing the senescence (scHPF_19) and stress (scHPF_14) factors in the AWS and hippocampus, as well as a depletion of microglia highly expressing the motility/adhesion (scHPF_9) and chemokine signaling (scHPF_5) factors in the AWS and the temporal cortex.

Given that postmortem processing may affect transcriptional profiles, we note that there were few consistent differences in factors associated with longer postmortem interval or time to tissue processing (**Figure S7**). Both samples from the New York Brain Bank (NYBB) and the Religious Order Study/Memory & Aging project (ROS-MAP)^24^ cohorts showed downregulation of the S100/TLR signaling (scHPF_23) factor in association with greater time-to-dissection and postmortem interval. However, while samples from the NYBB cohort showed association between longer time to tissue processing and lower C1Q^High^/phagocytic (scHPF_17) factor scores, in the ROS-MAP cohort the opposite effect was observed. Thus, these three factors may be more vulnerable to changes related to sample handling and should be interpreted cautiously. Overall, these analyses highlight some early evidence of association between certain traits and factors that need to be explored further and accounted for with proper covariates in downstream analyses.

### Projection of external data into our reference model

To demonstrate the broad utility of our defined factor model, we projected external datasets onto our model, including human single-nucleus data, as well as *in vivo* and *in vitro* model systems, with and without biological, drug, and genetic perturbations (**Figure 2**). Most notably, none of the projected datasets captured the full distribution of observable factor combinations derived from live human microglia. Nonetheless, the model is useful for describing within sample variation in repurposed datasets; for example, in glioma myeloid cells, we identified a strong downregulation of the stress factor (scHPF_14) (**Figure S8**) and upregulation of the HLA^High^/antigen presenting cell (APC) factor (scHPF_15) after topotecan treatment in slice culture (**Figure 2B,C** and **Figure S9**). The model was also broadly applicable to an *in vivo* model system of iPSC-derived, murine-xenografted microglia (xMGs), recapitulating a previously reported upregulation of the *GPNMB*^High^ factor (scHPF_26) in xMGs from 5xFAD-MITRG mice compared to WT (**Figure 2E-H** and **Figure S10**).

**Figure 2.**
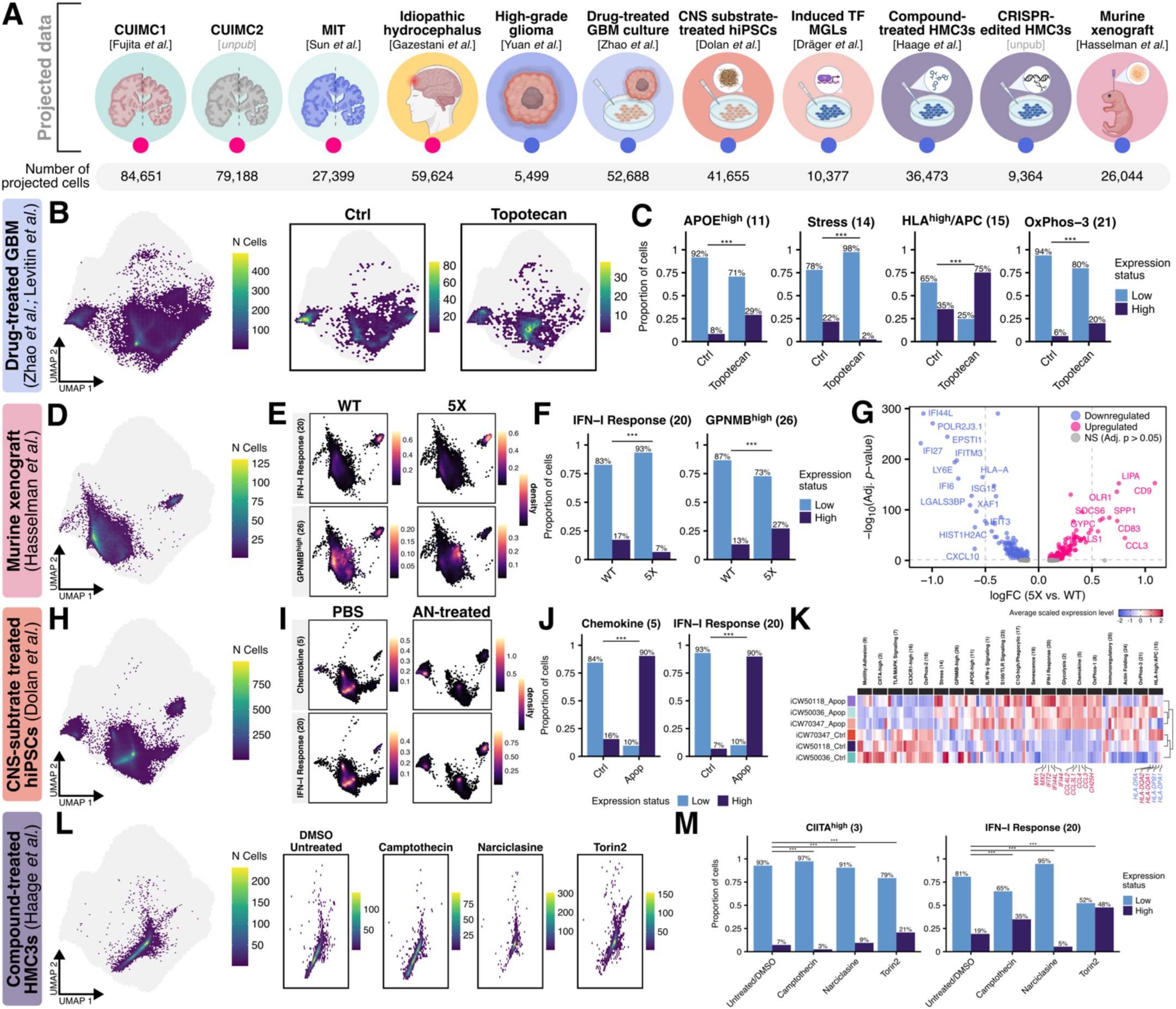
The 23-factor model offers a generalizable reference atlas of microglial functions and cellular processes, allowing for the projection of external data. **(A)** Overview of the external datasets projected onto the model for validation (see **Table S5**). **(B, C)** Projection of 65,179 myeloid transcriptomes isolated from glioblastoma slice cultures within the reference UMAP space (light gray background). Cells represented as binned, hexagonal ‘meta-cells’, colored by the number of cells within each bin. Showing examples of two treatment conditions (out of 14), dark gray meta-cells show control myeloid cells from the same donors. **(C)** Barplot comparing proportion of myeloid cells classified as low vs. high-expressing for topotecan across selected factors with significant perturbation compared to untreated cells. **(D-G)** Projection of 26,044 WT and 5xFAD-MITRG-xenografted human microglia shows upregulation of the *GPNMB*^High^ (scHPF_26) factor and downregulation of IFN-I response (scHPF_20) in 5X-xMGs **(F, G)**, capturing signatures observed at the individual gene level. **(H-K)** Projection of 41,655 iPSC-derived iMGs transcriptomes (Dolan et al., 2023) shows upregulation of two microglial subpopulations highly expressing IFN-I response (scHPF_20) and chemokine (scHPF_5) factors in AN-exposed iPSC-derived iMGs compared to untreated iMGs. **(L, M)** Projection of 36,473 HMC3 cells treated with various compounds. **(M)** Expression factor differences between treated and untreated/DMSO-treated HMC3 cells.

The factor model also showed the ability to capture perturbations in *in vitro* model systems. In iPSC-derived human microglia (iMGs), the model recapitulated cluster-derived signatures (**Figure S11A, B)**. We also recapitulated the observation of upregulated IFN-I response (scHPF_20) in iMGs treated with apoptotic neuron material; further, we also noted, in this context, a subset of microglia which co-express the IFN-I response and the chemokine signaling factors (scHPF_20 and scHPF_5) (**Figure 2I-L** and **Figure S11**). We also observed differential expression of factors in HMC3 cells treated with exogenous compounds. For example, HMC3 cells treated with Torin2, a second generation mTOR inhibitor, showed a strong upregulation of IFN-I response (scHPF_20), while cells treated with Camptothecin, a topoisomerase inhibitor, showed an upregulation of the *CIITA*^High^ factor (scHPF_3) (**Figure 2L, M** and **Figure S12**). Notably, however, signatures identified in *in vitro* model systems before or after treatments aimed towards polarization towards specific subtypes do not fully capture the distribution of microglial factors observed in scRNA-seq or snRNA-seq data (**Figure S13**).

More broadly, we noted that model systems exhibit common perturbations across factors which may indicate context-related transcriptional changes. For example, strong polarization towards IFN-I response (scHPF_20) was observed across multiple model systems, such as in iMGs induced by expression of transcription factors regardless of gene targets for CRISPR perturbations (**Figure S14**). In the case of CRISPR-edited HMC3 cells, different gene knock-outs resulted in similar dysregulation of oxidative phosphorylation factors (**Figure S15**).

### Discovery and replication of AD-associated factors

Projection of DLPFC snRNA-seq data (*n*=424 ROS-MAP participants^13^, CUIMC1 sample set) into our model shows that the nuclear transcriptome captures a significant proportion of the observable variation in microglial gene expression, but it fails to fully capture the distribution observed in the cell soma (**Figure 3A**). Nonetheless, the 23 factors showed substantial alignment with previous annotations based on traditional cluster-based analyses with marker genes, with several factors showing cluster specificity (e.g., *GRID2*^High^, *APOE*^High^, *GPNMB*^High^, IFN-I response) (**Figure S16A**). For example, the *GRID2*^High^ (scHPF_10) factor was uniquely enriched in the microglial homeostatic subtype, while IFN-I response (scHPF_20) and DAM-like (scHPF_11 and scHPF_26) factors were enriched in the Interferon-Response and Disease-Elevated microglial subtypes, respectively. Given the large sample size of this dataset, we were able to perform statistically well-powered analyses of our factors relative to AD outcomes. A number of microglial factors are significantly upregulated with worse AD outcomes at the donor level (FDR *p*-value < 0.05), including the glycolytic (scHPF_2), NPY1R^High^ (scHPF_4), *APOE*^High^ (scHPF_11) and *GPNMB*^High^ (scHPF_26) factors (**Figure 3B**), with the *GPNMB*^High^ (scHPF_26) factor being robustly associated with amyloid levels across all tested brain regions (**Figure S16C**). For completeness, we deployed an alternative strategy examining the data at the cell level (using *Milo*); in this secondary analysis, there was significant enrichment of the *GPNMB*^High^ factor (scHPF_26) (**Figure S17**, and **Figure S18**), which was supported by the upregulation of individual DEGs in tissues with intermediate to high NIA Reagan scores, including *GPNMB*, *LPL*, *MYO1E*, and *CD9* (**Figure S17I-J**).

**Figure 3.**
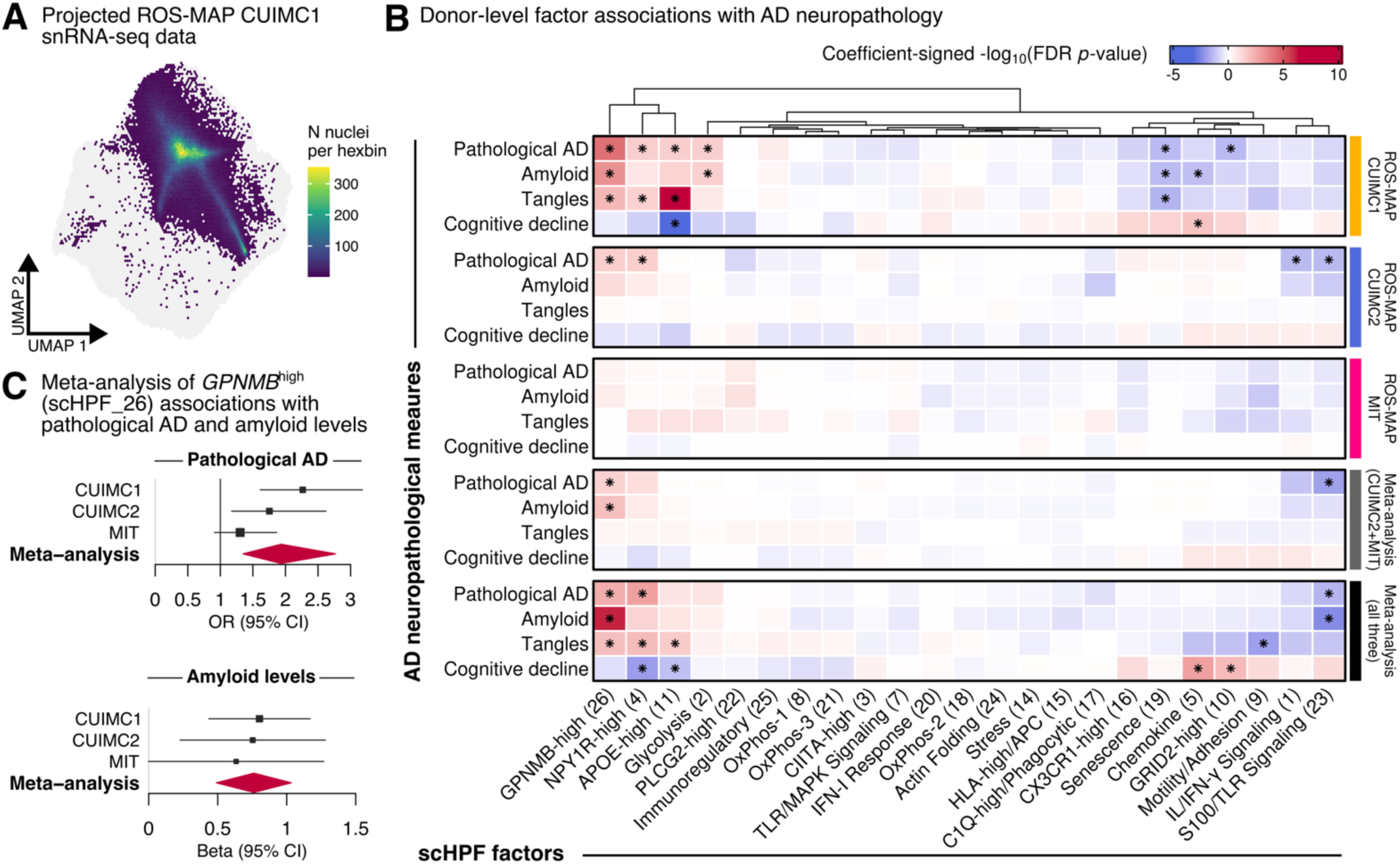
AD-derived microglia upregulate the disease-associated *GPNMB*^high^ factor. **(A)** UMAP embedding of microglial nuclei (*n*=84,651) from DLPFC snRNA-seq (ROS-MAP, *n*=424 donors) compared to the reference model (gray background). Cells represented as binned, hexagonal ‘meta-cells’ colored by the number of nuclei per bin (*nbins*=100). **(B)** Linear regression results for donor-level factor scores (mean-aggregated across cells) adjusted for study, age at death, sex, and postmortem interval. Includes a random-effects meta-analysis across the two datasets (see **Table S7** and **Table S8**). Significance levels: *FDR *p* < 0.05. **(C)** Forest plots for the meta-analysis of *GPNMB*^high^ (scHPF_26) across all three ROS-MAP cohorts in panel B.

We sought to replicate the significant association of factors with AD-related traits using two independent sets of ROS-MAP participants with snRNAseq data: (1) a new sample set produced at Columbia University (CUIMC2) (*n*=212) and (2) a repurposed dataset produced at MIT from which we removed overlapping samples (*n*=130 independent samples) (**Figure S19**). After rigorous FDR-correction for multiple testing (FDR < 0.05), we replicated the association of *GPNMB*^High^ (scHPF_26) with pathological AD and amyloid levels in the ROS-MAP snRNA-seq data (**Figure 3C**) and bulk RNA-seq data (**Figure S20**). We therefore prioritize this factor for further evaluation in downstream analyses involving regulatory networks. Ultimately, we assembled the CUIMC1, CUIMC2, and MIT datasets (*n*=766 unique participants) in a meta-analysis that summarizes all available data and prioritizes a few additional factors that are significantly associated with AD-related traits and deserve further investigation.

### Building a microglial transcriptional network to identify key regulators for each factor

To identify regulators of each microglial factor, we used ARACNe^25^ to reconstruct a predicted transcriptional regulatory network in our scRNAseq microglial resource by evaluating the covariation in expression of transcription factor (TF) genes and their putative targets (**Figure 4**). The resulting network included 676 regulons (a regulon is defined as a TF and its targets), including 8,478 TF target genes and 100,991 interactions (**Table S9**). The largest regulon was *NME2* with 4,204 targets, and the smallest was *EGR3* with 14 targets. The network also showed a high level of complexity with each regulon including a median of 6 other TFs. Several TFs were markers of factor identity themselves (present in the top 100 genes of one of our 23 factors); the chemokine (scHPF_5) factor had the largest number of TF genes in its signature (*n* = 20 TFs; see **Figure S21**).

**Figure 4.**
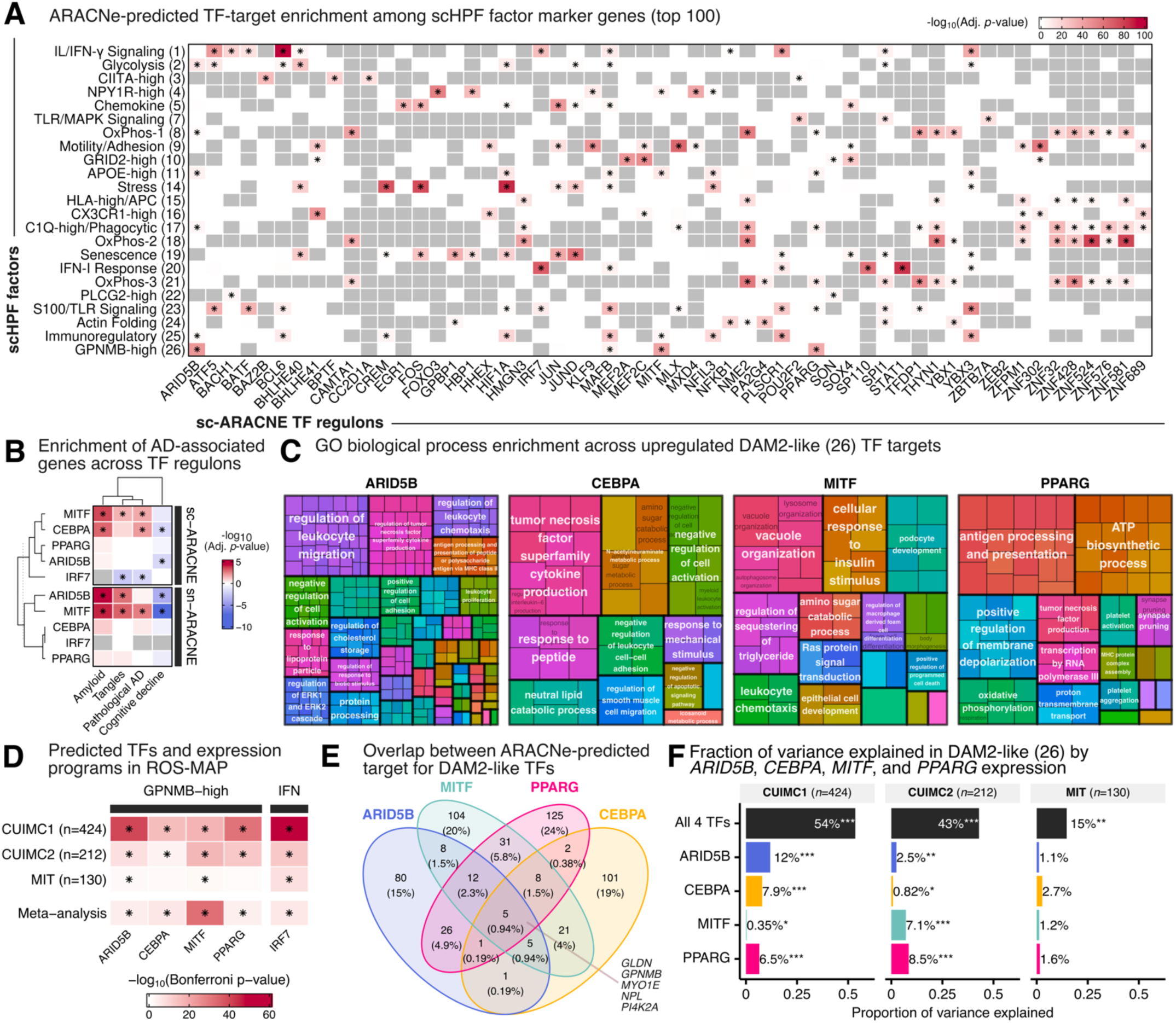
Microglial regulatory network identifies putative *GPNMB*^High^ (scHPF_26) factor regulators. **(A)** Heatmap showing hypergeometric enrichment of the top-100 loaded genes for each factor across ARACNe regulons (see **Table S9**). Grey indicates too few overlapping genes for testing. **(B)** Heatmap showing FGSEA enrichment of DEGs associated with AD-related traits in bulk RNAseq and their top enriched ARACNe regulons, in both the single-cell (sc) and single-nucleus (sn; see **Table S11**) ARACNE networks (see **Table S12**). Showing Bonferroni-corrected p-values across target regulons and traits. **(C)** Reduced GO enrichment results for upregulated regulon targets. Biological processes adjusted FDR *p*-value < 0.05 and *q*-value < 0.05. **(D)** Donor-level regulator associations with factors, adjusted for donor age, donor sex, PMI and sequencing depth. **(E)** Venn diagram showing predicted target overlap across TFs in the sc-ARACNE network. **(F)** Variance partitioning showing the unique total contribution of the four TF adjusted (X1|X2) for covariates (age, sex, PMI, study), as well as individual unique contribution of each TF adjusted (X1|X2) for covariates and the other three TFs. Asterisks denotes FDR-corrected *p*-value for the number of regulators from permutation test for redundancy analysis. *Significance levels*: *** *p* < 0.001, ** *p* < 0.01, * *p* < 0.05.

Concurrently, to identify regulons with robust signatures across the cytoplasmic and nuclear compartments, we built a second, independent ARACNe regulatory network in the ROSMAP CUIMC1 dataset (**Figure S22**). Overall, there was a marked overlap of predicted TFs between the single-cell and single-nucleus ARACNe networks: 663 of 676 TFs are shared (98%) between the two networks. However, there was considerable variability in the predicted target genes between the two networks (mean proportion of overlapping targets across factors = 18.4 ± 15.0%). Notably, the snRNA-seq-derived ARACNe network (83,908 predicted interactions) was smaller than the scRNA-seq-derived network.

In the single-cell data, regulons showed both unique and shared enrichment of targets across factors (**Figure 4A**). For the *GPNMB*^High^ factor (scHPF_26), the top five enriched regulons were *ARID5B*, *CEBPA*, *MITF*, *PPARG* and *MAFB*. These five TFs were also among the top 100 genes comprising the *GPNMB*^High^ factor (scHPF_26). Notably, *ARID5B*, *MITF*, *PPARG*, and *CEBPA* showed significant overlap in predicted target genes between the cytoplasmic and nuclear compartment (**Figure S22B**). Only *MITF* targets were robustly enriched for DEGs associated with worse AD-related clinicopathological traits in the single-cell and single-nucleus networks (**Figure 4B**). It has also been suggested that MITF may be a regulator of the DAM signature^10^. The MITF regulon was enriched across the *GPNMB*^High^ (scHPF_26), *APOE*^High^ (scHPF_11), NPY1R^High^ (scHPF_4), and immunoregulatory (scHPF_25) factors (**Figure 4C**). GO enrichment of upregulated and downregulated genes in the MITF regulon showed upregulation of genes involved in motility, lipid processing and foam cell differentiation, and downregulation of genes involved in inflammatory response, macrophage activation and cytokine production (**Figure 4C**). These results are consistent with published results.^26^

We also aimed to replicate the proposed regulatory network predictions. Therefore, we re-projected the three ROS-MAP snRNAseq data onto our model while excluding *ARID5B*, *CEBPA*, *MITF*, and *PPARG* from the expression matrix. Notably, donor-level expressions of *ARID5B*, *CEBPA*, *MITF*, and *PPARG* showed an independent, positive association with the *GPNMB*^High^ factor (scHPF_26) across all three datasets (**Figure 4D**). While there was an overlap across the targets of the four TFs (**Figure 4E**), variance partition showed that the TFs accounted for 0.03-12% of the variance in *GPNMB*^High^ factor (scHPF_26) expression individually; jointly, they explain an average 33% of variance (**Figure 4F** and **Figure S23**).

The *ARID5B* regulon showed the strongest potential for regulating the *GPNMB*^High^ factor (scHPF_26) in the single-cell and single-nucleus networks, with targets including DAM2 genes *SPP1*, *TREM2*, *CD63*, *CD68*, *CD9*, and *LPL*. While there was no significant pathway enrichment for targets downregulated by ARID5B, upregulated targets showed enrichment for antigen presentation, response to lipids, foam cell differentiation, and astrocyte activation (**Figure 4C**). Upregulated ARID5B scRNA-seq-derived targets showed enrichment of DEGs associated with cognitive decline, but not pathological traits (**Figure 4B)**. Targets from the snRNA-seq-derived network were also enriched for DEGs associated with amyloid, tangles, and cognitive decline (**Figure 4B**).

### A microglial MERFISH panel uncovers distinct topological patterns for the interferon and DAM2-like factors

We developed a new resource (**Figure 5A**): a microglial MERFISH panel of 412 genes that is designed to be deployed on the MERSCOPE platform to interrogate our 23 microglial factors at the single cell level in spatially-registered data (see Methods for details). The targeted gene list, presented in **Table S13**, includes 43 genes to identify previously defined microglial subtypes^27^ as well as 216 genes with which to measure each of the 23 microglial factors. We elected to focus our assessment of these spatially-registered transcriptomic data on the *GPNMB*^High^ (scHPF_26) and IFN-I response (scHPF_20) factors, so we added 164 probes to assess our proposed regulators of these factors and their targets. We then generated data from 12 frozen DLPFC tissue sections obtained from four donors (2 AD and 2 non-AD; *n*=2 female, ages 77-99 years), with three tissue sections per donor. In total, we extracted individual transcriptomes for 50,391 microglia (**Figure 5B**) whose location is registered within a given tissue section, of which 63% (*n*=31,731) are assigned to the cortex (gray matter) and 37% (*n* = 18,660) to the adjacent subcortical white matter.

**Figure 5.**
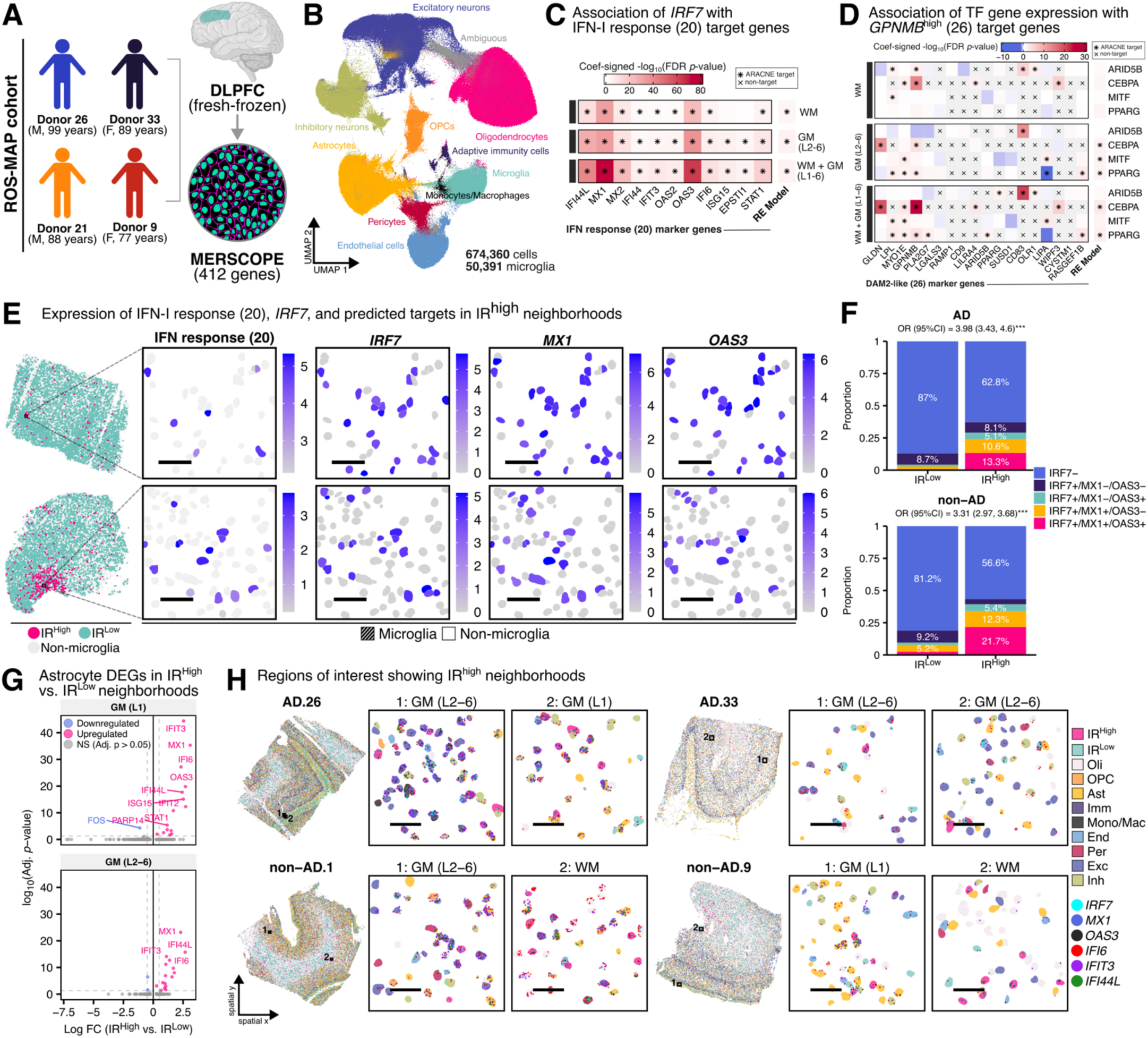
*In situ,* IR^High^ microglia show spatially-distinct neighborhoods characterized by high *IRF7* expression, and co-upregulation of IFN-I response (scHPF_20) genes in astrocytes. (**A**) Overview of the experimental design for MERSCOPE (see **Table S13** for gene list). **(B)** UMAP showing the major cell types identified. (C) *In situ* distribution of IFN-I response (scHPF20) high (i.e., IR^High^) vs. low (i.e., IR^Low^) cellular niches (see **Table S14** and **Table S15**). (D) *In situ* association results of *GPNMB*^High^ (scHPF_26) enriched TFs (*ARID5B*, *CEBPA*, *MITF*, *PPARG*) with their ARACNe-predicted targets across tissue layer niches (see **Table S14** and **Table S15**). **(E)** Regions of interest showing IR^High^ neighborhoods and the expression of *IRF7* ARACNe-predicted targets, *MX1* and *OAS3*. **(F)** Barplot showing the proportion of *IRF7*+/*MX1*+/*OAS3*+ microglia across IR^High^ vs. IR^Low^ microglia. **(G)** Volcano plot showing DEGs in astrocytes in IR^High^ versus IR^Low^ neighborhoods (see **Table S16**). **(H)** Regions of interest showing IR^High^ niches and the expression of *IRF7*, *MX1*, *OAS3*, *IFI6* and *IFIT3*.

The MERFISH-profiled microglia were projected onto the 23-factor scHPF model (**Figure S24**). Notably, the IFN-I response (scHPF_20) and *GPNMB*^High^ (scHPF_26) factors showed distinct distributions in transcriptomic space (**Figure 5C**). First, we explored whether ARACNe-predicted regulators of the IFN-I response (scHPF_20) and *GPNMB*^High^ (scHPF_26) factors were associated with their predicted targets *in situ*. The IFN-I response (scHPF_20) regulator *IRF7* was positively associated with the expression of its ARACNe-predicted targets, in both white and gray matter (**Figure 5D**). When considering microglia across grey and white matter, putative *GPNMB*^High^ (scHPF_26) regulators *CEBPA* and *PPARG* showed a significant association with ARACNe-predicted targets, while *MITF* only showed an association in grey matter (**Figure 5E** and **Figure S25**). However, none of the four *GPNMB*^High^ (scHPF_26) regulators (*ARID5B*, *CEBPA*, *MITF*, and *PPARG*) showed a significant association with ARACNe-predicted targets when considering microglia in white matter alone. Thus, despite very small sample sizes, we recovered some of the snRNA-seq-replicated associations of regulators and their targets using spatially registered data that is quantitative.

To further characterize our scHPF-derived factors, we explored whether microglia that display high-expression of the IFN-I response factor (scHPF_20) tend to cluster, as they might be responding to a local source of type I interferon. IFN-I response factor (scHPF_20) showed statistically significant spatial autocorrelation (i.e., non-random patterning of expression) (**Figure S27**), as well as local ‘hotspots’ (i.e., aggregation of high scores) which were prominently observed in two donors and across adjacent sections (**Figure S28**). We then aimed to explore the distribution of microglia highly expressing IFN response factor (scHPF_20). We defined microglia that display high-expression of the IFN response factor (scHPF_20) as those with expression greater than the median value + 2 times the median absolute deviation (hereafter referred to as IR^High^). IR^High^ microglia representing 3.63% of all microglia in our AD tissue and 5.77% of all microglia in our non-AD tissue. IR^High^ neighborhoods showed greater expression of the IFN-I response (scHPF_20) marker genes, as well as our IFN regulator, *IRF7* (**Figure 5F** and **Figure S24**).

Our hypothesis was that cells with high expression of the IFN-I response factor (scHPF_20) were likely to respond to a focal IFN signal and would therefore be more likely to form topologically restricted communities. *In situ*, IR^High^ microglia tended to occur more frequently in neighborhoods containing other IR^High^ microglia across white and gray matter (**Figure S29**). We found a significant positive correlation between the IFN-I Response (scHPF_20) factor expression and the Getis-Ord Gi* z-score^28,29^, suggesting a higher expression is associated with the presence of spatial hotspots, or clustering of high expression microglia. Notably, IR^High^ microglia were more likely to co-occur in spatial neighborhoods in gray matter (GM) than white matter (WM) (**Figure S29C**). In IR^High^ neighborhoods, most cell types, particularly astrocytes, showed significant upregulation of IFN-I response genes (**Figure S30** and **Table S16**), consistent with the concept that a source of IFN exists in the area and alters gene expression in its vicinity.

We also examined the *in situ* variability in spatial distribution of the *GPNMB*^High^ (scHPF_26) factor. We defined microglia high expression of this factor. *GPNMB*^High^ microglia were defined as those with an expression greater than the median + 2 times the median absolute deviation, representing 3.75% of all microglia in our AD tissue and 1.25% of all microglia in our non-AD tissue. Notably, the proportion of *GPNMB*^High^ microglia was lower in GM layer I (0.42%) compared to deep GM (2.66%) and WM (2.7%). The *GPNMB*^High^ (scHPF_26) factor showed significant patterns of spatial expression in WM (6 of 12 tissues), GM layer I (8 of 8 tissues), and GM layers II-VI (9 of 12 tissues) (**Figure S31**). We found a significant positive correlation between the *GPNMB*^High^ (scHPF_26) factor expression and the Getis-Ord Gi* z-score, suggesting a higher expression is associated with the presence of spatial hotspots, or clustering of high expression microglia. ‘Hotspots’ were present in both AD and non-AD tissue, in WM and deep GM (layers II-VI) (**Figure S32**). Across WM and GM layers, *GPNMB*^High^ microglia were more likely to co-occur in a neighborhood with another *GPNMB*^High^ microglia than *GPNMB*^Low^ microglia (**Figure S33**). However, we did not observe any differentially expressed genes in other cell types in the vicinity of *GPNMB*^High^ microglia (**Table S45**).

Notably, the frequency *GPNMB*^High^ microglia sharing a spatial neighborhood was higher in deep GM than WM, despite the overall proportion of *GPNMB*^High^ microglia being similar, suggesting layer-specific effects.

## Discussion

Here, we present a comprehensive, cross-disease model of microglial factors based on the largest live microglia dataset assembled to date; this resource includes 441,088 transcriptomes broadly sampled across brain tissues and neurodegenerative disorders and is available to be repurposed (see **Data Availability**). The 23-factor model demonstrates broad utility across sequencing platforms, across *in vitro* and *in vivo* human model systems, as well as model systems with biological, chemical, and genetic perturbations. None of the model systems fully captured the spectrum of observable microglial signatures from living human cells, highlighting the importance of a human-derived reference atlas. However, our factor-based model facilitates comparisons among datasets that is readily biologically interpretable; further, the projection of external datasets into the proposed model is straightforward, without the need for label transfer. Our model (**Figure 1**) provides an alternative to traditional subtyping approaches, such as clustering, one that offers finer resolution for capturing different dimensions of microglial function and cellular processes.

Using scHPF, we defined 23 factors which are non-orthogonal (statistically independent) latent factors in a lower cell and gene dimensional space capturing continuous gene expression signatures rather than discrete cell subpopulations. Challenges associated with unsupervised cell clustering have been well noted, particularly the interpretation of cluster annotations and whether they represent biologically meaningful distinctions between cell subtypes.^30^ The factor-based approach to defining continuous expression signatures has already been successfully applied, specifically to T-cells, and has shown good alignment with cluster-based annotation and utility for identifying subtle disease-associated or treatment-associated cellular changes.^19,20,31^ Indeed, we observed substantial alignment between our factors and independently-conducted cluster-based annotation for microglia in single-nucleus data from the ROS-MAP cohorts. The strongest observed alignment was observed between the IFN-I response factor (scHPF_20) and the single-nucleus IFN responsive cluster, a subset which has been inconsistently observed in previous studies.^5,7,13,16,26,32^ In the case of rare microglial signatures, such as interferon responsiveness, increasing sample size may improve detection rates but cluster-based misclassification may also mask this signature. Notably, our factors are not necessarily exclusive to any ‘subtype’ as all cells are scored for every factor, thereby allowing for characterization at finer resolution.

Our 23-factor model provides a comparative context for understanding microglial heterogeneity in external datasets and can serve as a reference nomenclature for characterizing microglia broadly. Within the context of human-derived microglial factors, we observed that most model systems showed limited heterogeneity which is unsurprising given that model systems likely do not fully recapitulate the complexity of human microglia. Notably, none of the model systems sufficiently captured the homeostatic signatures (e.g., *GRID2*^High^) and several of the models showed unique polarization towards extreme signatures, including the IFN-I response factor (scHPF_20). In response to perturbation, while we observed some changes in model systems, the effect sizes were small, potentially due to the limited starting heterogeneity and the limited capacity of the model for differentiation. In addition, while we observed little consistent effect associated with longer time to tissue processing and postmortem interval in scRNA-seq and snRNA-seq data. Such effects doubtlessly occur, and we highlighted some factors that should be interpreted cautiously given possible effects related to sample processing in the discovery cohort: C1Q^High^/phagocytic factor (scHPF_17) and the S100/TLR signaling factor (scHPF_23). Overall, these observations highlight the importance of a reference atlas of microglial functions based on live, high-quality human microglia.

Generally, the derived factors were highly interpretable. However, several factors require further characterization, including two factors, *CIITA*^High^ (scHPF_3) and *PLCG2*^High^ (scHPF_22), which did not have clear biological pathway enrichment. We observed four factors capturing pathways related to heterocyclic metabolism and oxidative phosphorylation that potentially point towards the complexity of metabolic function in microglia. Previous studies have noted the importance of microglial metabolism in AD, as well as in the regulation of key functions, such as phagocytosis.^33,34^ While the oxidative phosphorylation factors are highly correlated, it remains to be clarified which aspects of oxidative phosphorylation each one is capturing.

We also observed the immunoregulatory factor (scHPF_25), characterized by *CD163*, *PLXNC1*, *TGFBI*, *F13A1*, and *ANXA1*, which requires further exploration. Discrimination between microglia and macrophages (e.g., perivascular, infiltrating) is difficult. Despite our stringent quality control to exclude non-microglial transcriptomes in our data pre-processing pipeline, it remains unclear whether the immunoregulatory factor (scHPF_25) is capturing microglial biology or macrophage-associated signatures. While CD163 has been shown to be a distinguishing marker of perivascular macrophages, it has also been shown to be expressed by ‘activated’ anti-inflammatory microglia.^35^ Similarly, *F13A1* has been identified as a brain macrophage marker^36^, while *ANXA1* and *TGFB1*^37^ have been shown to be anti-inflammatory microglia markers. In an scRNA-seq dataset of GBM-derived myeloid cells, although there were strong polarization cells toward the niche of the immunoregulatory factor (scHPF_25), the expression of other factors was also observed, particularly the chemokine factor (scHPF_5), suggesting that multiple factors are broadly applicable to other, non-microglial myeloid cells. While it is possible that the immunoregulatory factor (scHPF_25) may be capturing a specific signature from potential doublets or macrophages that were not removed during quality control, we note that the distribution of the immunoregulatory factor (scHPF_25) in the discovery dataset is not restricted to a small subset of cells, with 99.9% of cells expressing the factor. Future iterations of our modeling will examine a broader set of CNS macrophages to better characterize the extent of sharing of transcriptional programs between microglia and bone-marrow derived microglia.

Microglia have a large and complex network of transcriptional regulators which are stable across the datasets that we have examined. Across two independently-derived regulatory networks (single-cell and single-nucleus), we observed a high overlap between putative TF regulators, but less overlap between downstream targets for key TFs. Nonetheless, we recovered well-validated regulator relationships, such as regulation of interferon response by IRF7, IRF9, STAT1, and STAT2^38^. Our ARACNe network pointed toward key regulators targeting specific factors, including a previously reported putative regulator of the DAM signature, *MITF.*^26^ We also identified and replicated *ARID5B*, *CEBPA*, and *PPARG* as additional regulators of the *GPNMB*^High^ factor (scHPF_26). In snRNA-seq data, we show that collectively, *ARID5B*, *CEBPA*, *MITF*, and *PPARG* may account for up to 54% of the variation observed in the *GPNMB*^High^ factor (scHPF_26) factor with multiplicative effects. *In situ*, there was evidence that *CEBPA* and *PPARG* may be regulating the *GPNMB*^High^ factor (scHPF_26), particularly in gray matter, which requires further investigation. Overall, this suggests that these TFs may play a major role in the regulation of the *GPNMB*^High^ factor (scHPF_26) and are interesting targets to pursue as part of therapeutic development efforts targeting this factor that we identify and replicate as being associated with AD. Previous studies provided evidence that *ARID5B* chromatin peaks were enriched in activated microglia states^12^ and that *ARID5B* expression is increased in an aging-associated homeostatic microglia subtype.^39^ Other studies report that CEBPA has a suppressive effect on homeostatic and anti-inflammatory genes and that there are microglia which highly co-express *PPARG* and *APOE*, a key DAM2 gene.^12,40^ Interestingly, like the pattern of clustering seen among IR^High^ microglia and adjacent cell types (**Figure 5** and **Figure S26**), microglia that express high levels of the *GPNMB*^High^ factor (scHPF_26) appear to be spatially more proximal to each other than *GPNMB*^Low^ microglia in tissue, but *GPNMB*^High^ do not appear to be in cellular neighborhoods that are eliciting a similar gene signature, suggesting that they may not be responding to a spatially limited focus of a given molecular signal in the tissue. Rather, they may reflect an altered state of the parenchymal tissue or a response to a widespread dysfunction of neuronal circuits. We did not observe any upregulated genes in astrocytes or other cell types in proximity to *GPNMB*^High^ microglia; however, it is not clear how proximal the *GPNMB*^High^ microglia are to amyloid β plaques, as our MERFISH analyses did not include amyloid β plaque staining. Studies have shown that microglia upregulate DAM markers in proximity to plaques.^41–43^ Astrocytes, which play a key role in the cellular response to AD,^44^ have also been show to upregulated disease-associated markers in proximity to plaques in the cortex and hippocampus, including *Itm2b*, *Cpe*, *C1qa*, *Atp1a1*, *Ckb*, *Kcnip4* in mice and *SERPINA3* in humans.^41,45^ However, our MERFISH panel had a limited set of genes and it is possible that we could not detect the panoply of disease-associated genes in other cell types, which may have cell-type specific responses in proximity to AD pathology.

Our study has certain other limitations. There are brain states – such as neoplastic tissue – that we have not yet sampled, and, aside from the AD-spectrum samples, each of the diseases sampled to date only has a small number of donors given that our goal was to derive a cross-disease resource. Cross-disease comparisons are of great interest, but they will require much larger sample to be properly powered statistically.

In conclusion, our cross-disease model of microglial factors offers a resource with which to advance our understanding of microglial states in a more nuanced fashion, and it already has yielded insights into biological processes associated with AD. We have identified and replicated the role of key transcriptional regulators – *ARID5B*, *CEBPA*, *MITF*, and *PPARG –* for the DAM2-like factor (scHPF_26) that we have shown to be associated with AD; these results are consistent with reports implicating microglia with a transcriptional signature involving lipid processing and enriched for the DAM signature in AD^21,46^. Our structured approach now provides robust candidates with which to (a) mimic and study factor (scHPF_26) in vitro and (b) guide the development of microglial immunomodulation strategies. Using a large discovery cohort of live human microglia, we show that our model is sufficient to support the analysis of a variety of external datasets, illustrating the power of a shared framework to accelerate the exchange of information and insights in the human microglial community. It represents a step towards a reference nomenclature for microglial transcriptional programs and states that needs to be developed by the community of interested investigators. Future companion models for other cell types in the brain may further capture the complexity of microglial and their cellular interactions that influence the onset and course of neurodegenerative disease.

## Methods

### Source of CNS specimens

The tissue samples were obtained across several sites, including Columbia University Medical Center/New York Brain Bank (New York, NY), Rush University Medical Center/Rush Alzheimer’s Disease Center (Chicago, IL), University of Washington Medical Center (Seattle, WA), Brigham and Women’s Hospital (Boston, MA), Rocky Mountain MS Center (Denver, CO), Massachusetts General Hospital (Boston, MA), and Banner Sun Health Research Institute (Sun City, AZ). Tissue specimens were obtained with informed consent or through donation programs at each institution with approval from their ethical review boards. All procedures and protocols were approved by the Institutional Review Board of Columbia University Irving Medical Center (protocol AAAR4962). For a detailed description of tissue samples see **Table S2**.

### Tissue dissection, microglia isolation and sorting

Tissue samples (from autopsy or epilepsy surgical resection of the temporal lobe) were shipped overnight with priority at 4°C in ice-cold medium (Hibernate-A medium (Gibco, A1247501) containing 1% B27 serum-free supplement (Gibco, 17504044) and 1% GlutaMax (Gibco, 35050061).

Microglial isolation was performed on ice as per our published protocol with minor modifications.4 Briefly, after dissection, the tissue was placed in HBSS (Lonza, 10-508F), weighed, and then homogenized in a 15 mL glass tissue grinder in increments of 0.5 g. The homogenate was filtered through a 70 μm filter and then spun down at 300 RCF for 10 mins. The pellet was resuspended in 2 mL staining buffer (RPMI (Fisher, 72400120) containing 1% B27) per 0.5 g of initial tissue and incubated with anti-myelin magnetic beads (Miltenyi, 130-096-733) for 15 mins, as per the manufacturer’s specification. The homogenate was then washed once with a staining buffer, followed by depletion of the myelin using large separation columns (Miltenyi, 130-042-202).

After spinning down, the cell suspension was incubated for 20 mins on ice with anti-CD11b AlexaFluor488 (BioLegend, 301318), anti-CD45 AlexaFluor647 (BioLegend, 304018), and cell hashing antibodies (see Methods for batch processing), as well as 7AAD (BD Pharmingen, 559925). The cell suspension was then washed twice with a staining buffer. Cells were filtered through a 70 μm filter to capture CD11b+/CD45+/7AAD-or CD45+/7AAD-cells for sorting on a BD FACS Aria II or BD Influx cell sorter. Cells deriving from different brain regions were sorted separately in an A1 well of a 96-well PCR plate (Eppendorf, 951020401) containing 100 μL of PBS buffer with 0.3% BSA. The cells were then sorted using a 100 μm nozzle with variable sorting times and speeds (averaging 10-20 mins per sample with a speed of 8,000-10,000 events per second) depending on sample quality.

Library construction and sequencing was performed according to our published protocol,8 with minor modifications. FACS-sorted CD11b+/CD45+/7AAD-or CD45+/7AAD-cells from the same donor were pooled and single-cell libraries were prepared according to the manufacturer’s protocol using 10x Chromium Next GEM Single Cell 3’ Reagent Kits v3.1 (Dual Index) with Feature Barcode technology for Cell Surface Protein (10x Genomics, Pleasanton, CA). Cells were loaded on the Chromium Controller (10x Genomics). To generate single-cell gel beads in emulsions (GEMs), reverse transcription reagents, barcoded gel beads, and partitioning oil were mixed with the cells. Libraries were generated and sequenced from the cDNA, and the 10x barcodes are used to associate individual reads back to the individual partitions. Incubation with GEMs produced barcoded cDNA from poly-adenylated mRNA and DNA from cell surface protein. After incubation, the GEMs are broken, and the pooled fractions are recovered. For library construction, cell-barcoded cDNA was amplified via PCR and then separated by SPRI size selection into cDNA fractions containing mRNA derived cDNA (>400bp) and HTO-derived cDNAs (<180bp), which were further purified by additional rounds of SPRI selection. Independent mRNA and HTO libraries were pooled and sequenced together on an Illumina HiSeq4000 or a NovaSeq 6000.

FASTQ files were processed using Cell Ranger v6.0.0 (10x Genomics). Reads were aligned to the human GRCh38 reference genome and a UMI count matrix was generated based on the transcriptome model 2020-A (10x Genomics). For multiplexed samples, reads of HTO libraries were processed together by applying a pattern “5PNNNNNNNNNN(BC)NNNNNNNNN” to the R2 reads.

### Quality control and demultiplexing

We filtered potentially low-quality or apoptotic cells based on fraction of transcripts from 24 mitochondrial genes prior to gene filtering based using a threshold of 2 SD deviations above the median absolute deviation or greater than 1.5 × the interquartile range. Cells with low or high UMI count (500 < UMI < 10,000) and four batches, each with less than 150 transcriptomes, were excluded.

Tissue received on the same day was multiplexed using hashtag oligonucleotides (HTOs; see **Table S2**) and loaded on a single 10x Chromium 3’ Chip. In total, there were 191 unique batches including 161 unique donors. For hashed batches, we retained transcriptomes with both RNA and HTO matrices for demultiplexing. For most batches, we used *demuxmix*^47^ to fit a two-component mixture model to assign transcriptomes to a donor of origin. For batches showing poor demultiplexing with demuxmix, we evaluated their demultiplexing results from demuxmix versus *HTOdemux*^48^ and *MULTI-seq*^49^ to preserve as many transcriptomes as possible. For *HTOdemux* and *MULTI-seq*, HTO counts were normalized using centered log-ratio transformation.

Conservatively, we used *DoubletFinder*^50^ and *scrublet*^51^ to identify additional potential multiplets, excluding any transcriptomes flagged as a multiplet by either method. Across all batches, DoubletFinder identified 23,786 potential doublets (4.37%). Across all batches, Scrublet identified 2,000 potential doublets (0.037%).

### Batch Correction

Library preparation batches and diagnoses were confounded by changes in sequencing technology (i.e., 10X chemistry version 2/3) as samples were collected across various sites during different time periods. After log-normalization, we extracted 40 PCs and retained the first 34 PCs which each explained at least 5% of variance and achieved a total cumulative variance of 91%. Cells sequenced across different batches were projected into a shared embedding using Harmony.^52^ After integration, median Local Inverse Simpson’s Index (LISI) scores were comparable between version 2/3 chemistries. For differential expression analysis, we conducted downsampling of batches with the deepest sequencing coverage (version 3 chemistry) using downsampleMatrix from DropletUtils^53^ on a per-batch basis to match the coverage of batches with shallower coverage (version 2).

### Cell type identification

After quality control, we retained 464,680 transcriptomes across 161 donors. Based on log-normalized downsampled transcript counts and 34 retained PCs, we built a KNN graph (*k.param* = 20) in transcriptomes passing quality control using *FindNeighbors* from *Seurat* v4.4.0 and identified major cell type clusters using the Louvain algorithm via *FindClusters*. For annotation, differentially expressed genes (DEGs) were identified using MAST^48^ with downsampled counts. We excluded transcriptomes expressing markers of red blood cells, non-immune cells (oligodendrocytes, astrocytes, neurons), adaptive immune cells, monocytes, and CNS macrophages. In total, we retained 441,088 transcriptomes that were likely microglia which constituted 94.9% of the cells passing quality control.

### scHPF model building and evaluation

scHPF is a recently reported Bayesian factorization method that enables discovery of both continuous and discrete expression patterns from scRNA-seq count data.^19,20^ This algorithm is particularly well suited for extraction of signatures that vary continuously among transitioning cells of the same type.^31^ The consensus scHPF approach involves four primary steps: (1) subsample and downsample the training data to balance batches and sequencing depth, (2) generate multiple scHPF models using different values of k, which represents the number of factors in a given model, (3) cluster the factors from different scHPF models to determine stable factors across models, and (4) initialize and train a final model from the cluster medians calculated in (3).

We conducted the process of generating the consensus scHPF model twice, once to produce a model containing only samples generated with the 10x v3 chemistry, and once with all samples from both 10x v2 and v3 chemistry. By virtue of the availability of samples and the advent of newer 10x chemistry versions, our data has confounding between diagnosis and 10x chemistry version that is difficult to resolve. We chose this approach to enable the comparison of the two different models to explore technical artifact-associated factors. As such, we prepared loom objects using the SeuratDisk extension to Seurat separately for these two versions. To mitigate the effects of lower-quality samples, we retained only samples with greater than 1,500 cells in the v3-only model and greater than 1000 cells in the combined model. This is because in the consensus scHPF workflow, it is necessary to subsample to equalize cell number and downsampled to mitigate depth-based artifact.

Notably, we found that factors between these two versions were quite similar and chose to proceed with the 10x v3-chemistry model to mitigate technical differences, as we could project 10x v2 data into the latent space of the model built only from 10x v3 cells.

### Generating consensus scHPF factors

We subsampled based on donor ID to equalize numbers of cells between batches, and downsampled each sample so that the median UMI number per cell in different batches was equivalent to the lowest median UMI count among our samples to reduce depth-based artifact. In the case of the v3.0 dataset, we downsampled to a median UMI count of 854, based on the deidentified ID of each sample.

To choose factors that represent “real” biological signals specific to myeloid cells, instead of artifacts or signals from other cell types present in the original data that were incompletely scrubbed from our reference data, we took a multi-step process. First, we took our entire dataset, which at baseline contains microglia, adaptive immune cells, macrophages, monocytes, red blood cells, and non-immune cells concordant with neuronal or glial cells. We computed pairwise differential expression between the cells we identified as microglia and each other cell type. To do so, MAST was applied to normalized count data from the “SCT” assay of the Seurat object to find DEGs between every combination of pairs of cell identities. Within each cell type, all the DEGs that were identified with this approach were filtered to only include those that were only found to be differentially expressed in one direction (either up or down). Any genes that were found to be upregulated in comparison to some clusters but downregulated in comparisons to other clusters or vice versa were removed from our downstream analysis. A minimum log fold-change threshold of 0.25 and a minimum percent of cells expressing the marker in question of 0.05 was used.

To evaluate enrichment of genes from non-microglial sources, we performed Fisher’s exact test with Benjamini-Hochberg correction using the top 100 “contaminant” genes per source in the top 100 genes for each factor. While we found some factors that showed enrichment for genes that were more prevalent in other cell sources, manual examination of each of these factors and GO pathway analysis suggested functional roles that could still be present in microglia, so no factors were chosen for removal at this step. For example, one factor that was enriched in macrophage-associated genes was retained, as we felt that the genes and pathways contained therein were likely to be relevant in microglia. Next, we examined genes for signals suggestive of dissociation artifacts. GO pathway analysis was performed with clusterProfiler.^54^ For GO analysis, we conducted analysis with biological process annotation, and the Benjamini-Hochberg correction16 was used to correct p-values for multiple testing. Corrected p-values below a threshold of 0.01 were chosen as significant for annotation results. We also manually examined factors for evident trends. For example, factor 12 in our original model was dominated by histone genes, which we considered to be concerning for technical artifacts. Notably, factors 6 and 13 in the original model were exclusively annotated in association with heat stress and unfolded protein response, respectively, and were thus removed from the model. Other factors had some contribution of these pathways, but also had other functional annotation that emerged, such as interleukin signaling in factor 19 and actin folding in 24, that led us to keep these in the interest of retaining the maximal possible biological signal. Factors were also examined in association with sample ID to ensure that no factors were predominantly found in one sample or individual. Factors were examined for association with gender, but no specific factors associated with sex were found.

### MitoCarta

Given that four of the 23 factors were potentially capturing metabolic signatures, we wanted to explore whether these signatures could point towards mitochondrial pathways. We conducted hypergeometric enrichment analysis for the top-100 loaded genes for each factor across *MitoCarta3.0*^22^ pathways. Hypergeometric tests were conducted using *enricher* from the R package *clusterProfiler* across 77 MitoCarta pathways with at least 10 genes from a universe of 19,247 possible genes.

### Differential abundance testing

Given our 23-factor model, we aimed to identify differentially expressed microglial signatures across tissue– and disease-dependent contexts. To do so, we used *Milo* v.1.9.1 which defines cellular neighborhoods using a *k*-nearest neighbor (KNN) graph within a lower dimensional space tests for differential abundance between conditions while adjusting for covariates and accounting for the non-independence of overlapping neighborhoods.^23^ Using the *Milo* framework, we explored the differential abundance of microglial subpopulations to better characterize the scHPF factors.

### Sex

To mitigate confounding of sex across diagnoses, we restricted the *Milo* analysis to AD tissue only, focusing on four brain regions (AWS, BA9/46, BA20/21, hippocampus) (**Figure S4A, B**). The dataset was downsampled to a maximum of 500 cells per donor per region (85,678 cells). Next, we built a graph using the 23-factor embedding across *k* = 100 nearest neighbors. Vertices were randomly sampled to a proportion of 5% to define neighborhoods and refined the “graph” scheme (**Figure S4C**). Euclidean distance was calculated between cells in a neighborhood using the 23-factor embedding. To explore differential abundance across sexes, we fit a *Milo* model for sex (female = 0, male = 1), adjusted for age, region, and recruitment site. Per neighborhood, differential abundance was calculated as the log-fold change between conditions. Differential abundance testing was adjusted to account for overlapping within the graph network using a spatial FDR weighting scheme (spatial FDR-corrected *p*-value < 0.05). Subsequently, overlapping and concordantly differentially abundant neighborhoods were grouped into larger pseudo-cluster, ‘meta-neighborhoods’ using the Louvain clustering algorithm with a log fold-change greater than 1 required to retain edges between adjacent neighborhoods (**Figure S4D**). To identify overall enriched or depleted meta-neighborhoods, we compared the distribution of log-fold change across individual neighborhoods comparing a single meta-neighborhood versus all other neighborhoods (Student’s *t*-test, FDR-corrected *p*-value < 0.05) (**Figure S4E**). Since the neighborhoods were defined within the scHPF space, we could also aggregate factor scores at the neighborhood level. Therefore, to characterize factor expression, we compared the distribution of median factor scores within a single meta-neighborhood versus all others (Wilcoxon test, FDR-corrected p-value < 0.001) (**Figure S4F**).

### Age

Given the confounding of age across diagnostic categories (**Figure S5A**), we restricted the analysis to AD tissue only, focusing on four brain regions (AWS, BA9/46, BA20/21, hippocampus). The dataset was downsampled to a maximum of 500 cells per donor per region (85,678 cells). To explore differential abundance with increasing age, we fit a *Milo* model with age discretized to compare older versus younger age (age > 85). The model was adjusted for sex, region, and recruitment site (**Figure S5B-F**). At the donor level, we calculated mean-aggregated factor scores to explore the association with older age (> 85 years) after adjusting for the effect of sex across the four brain regions. Regional effects were then meta-analyzed across each factor (**Figure S5G**).

### Brain regions

Similarly to age and sex, we aimed to mitigate the confounding of different brain regions sampled across diagnoses, therefore, we again restricted the *Milo* analysis to AD tissue only and the four commonly sampled brain regions sampled for AD (AWS, BA9/46, BA20/21, hippocampus). The dataset was downsampled to a maximum of 500 cells per donor per region (85,678 cells). Models were adjusted for age, sex, and recruitment site.

### Postmortem interval (PMI)/time-to-dissection

For differential abundance analysis, we analyzed the NYBB cohort independently including microglia from AD tissue only. The dataset was downsampled to a maximum of 1,000 cells per donor per region (59,718 cells). In the NYBB cohort, time-to-dissection is calculated as the (time starting processing + 1hr) – (time of death). One hour is added to account for the time required for harvesting and freezing the tissue samples. We defined higher vs. lower time-to-dissection using a median split. Subsequently, we fit a *Milo* model adjusted for sex and age. Similarly, we analyzed tissue from donors from the ROS-MAP cohort, which defines PMI as the time interval between death and tissue preservation in hours. The dataset was downsampled to a maximum of 1,000 cells per donor per region (33,601 cells). The *Milo* model was adjusted for sex, age, and pathological AD status.

### Factor associations with histopathological measures

We conducted FGSEA to identify scHPF factor enrichment across DEGs for AD-related outcomes, including clinical AD diagnosis, pathological AD diagnosis (dichotomized NIA-Reagan score), rate of cognitive decline adjusted for education, amyloid β levels and neurofibrillary tangle density in the ROS-MAP cohort. We used summary statistics from previously conducted differential expression in bulk RNA sequencing data across 18,629 genes from 1,092 DLPFC samples were run using DESeq (R package *DESeq2*) and were adjusted for age at death, sex, and technical variables. DEGs with an FDR-adjusted *p*-value less than 0.05 were included in FGSEA. For scHPF factors, we tested the top 100 loaded genes.

We also projected several external datasets onto the scHPF model (see full description below), including a subset of the ROS-MAP cohort with available snRNA-seq data was projected onto our scHPF model to further explore association with AD proteinopathies (*N* = 424 donors). We tested the association between factor scores and Braak/CERAD scores. A logistic regression was fit for donor-aggregated factor scores in association with dichotomized Braak/CERAD score, adjusted for age, sex, study, and PMI as (i.e., dichotomized score ∼ age + sex + study + PMI + factor score). For CERAD, ‘low’ was defined by categories 0/1 and high was defined by 2/3. For the Braak stage, low was defined by stages 0-3, and high was defined by stages 4-6.

A subset of the discovery cohort with available neuropathological measures (Braak stage, *N*=116; CERAD score, *N*=109). In the discovery cohort, we tested the association between donor-aggregated factor scores and Braak/CERAD score. Using a logistic linear regression for high vs. low Braak/CERAD score, models were included donor-aggregated factor scores adjusted for age and (i.e., dichotomized score ∼ age + sex + factor score). For models including data from all source sites (NYBB, RUSH and UWA), models were also adjusted for site. For CERAD, ‘low’ was defined by categories 0/1 and high was defined by 2/3. For the Braak stage, low was defined by stages 0-3, and high was defined by stages 4-6.

### Histopathological quantification of tau in the frontal cortex, temporal cortex, and hippocampus

To explore differences in factors of microglia from different histopathological contexts, we conducted phosphorylated tau (pTau) AT8 staining in tissues from BA9/46, BA20/21, or hippocampus. First, 6 μM paraffin-embedded tissue sections were deparaffinized with CitriSolv Clearing Agent (Decon Laboratories, In., 1601) for 20 min at Room Temperature (RT), followed by an ethanol series (100%, 100%, 70%) for 30 seconds each. The slides were then put in distilled water for 1 min at RT. Antigen retrieval was performed with pH 6.0 citrate (Sigma-Aldrich, C9999) and heating with a microwave for 25 min at 400 watts. After 5 min in tap water, slides were blocked using 3% BSA and then incubated with pTau AT8 primary antibody (Invitrogen, MN1020) at 4C overnight.

The secondary antibody (ThermoFisher Scientific, A32787) for the pTau staining was diluted in PBS at 1:500 and then incubated for 30 min at RT. After washing with PBS, the lipofuscin autofluorescence was quenched using Trueblack (Biotium, 23007) for 2 min at RT and slides were mounted using the Prolong Gold with Dapi (Thermo Fisher Scientific, P36941).

Images were captured using the Nikon Eclipse Ni-E immunofluorescent microscope at a 20x magnification, covering all six cortical layers in a zigzag pattern, with approximately 35-40 pictures taken per subject. Images were subsequently analyzed using CellProfiler software, where tangles and pTau plaques were size-filtered, and the ‘area occupied’ measurement was used for the analysis.

To test the association between donor-level and cell-level factor scores and area occupied by tau plaques/tangles (high vs. low area defined by the median), we fit LMMs. For donor-level mixed-models were adjusted for age, sex, time-to-tissue processing (> 12 hours), brain region and the random-effect of donor. At the cell-level, models were adjusted for donor age, sex, time-to-tissue processing (> 12 hours), cell UMI count and cell proportion of mitochondrial gene expression, as well as the random-effect of donor. For cell-level models combining microglia from across brain regions, models were also adjusted for brain regions. For model building, the dataset was downsampled to a maximum of 1,000 cells per donor per region.

### Projecting query datasets onto the scHPF model

To project query datasets for projection into the scHPF model, we began by evaluating genes included in scHPF model construction that were missing in query datasets. Depending on the origin of the query data, which came from different institutions, types of technologies, and models, there were anywhere between zero and 437 missing genes. In looking at the missing genes, we evaluated the highest rank they held in our scHPF factors, which are ordered by contribution of genes.

To account for missing genes relative to our reference, we chose to add zeros, which aligns with practice in other recent label transfer approaches (e.g., *scArches*^55^). Query data is reconstructed around the expression matrices with the full gene set included and converted to loom objects as described above. Query data is downsampled to the same threshold of 854 for median UMIs, prepared for scHPF projection, projected into the reference model, and then scored. Data projected in this way first included 10x v3 chemistry microglia from our data that were excluded from the training set and 10x v2 chemistry microglia that we had removed as part of only computing a model from our 10x v3 data. In doing so, we computed scHPF factor scores for all query cells based on a framework derived purely from our 10x v3 data. Using a UMAP model trained on the retained scHPF factors, we also projected query datasets into this low-dimensional reduction. In addition, we projected external datasets onto the scHPF model for factor validation which are described below (**Table S5**).

### High-grade glioma (Yuan et al., 2018).^56^

We projected 5,499 myeloid cells from high-grade glioma surgical resections across eight donors (sex: 3 female, 5 male; age: 49-74 years) and three tumor types, including classical (*n*=3), mesenchymal (*n*=2), and proneural (*n*=3) types (**Figure S8**). To explore differences in factors across tumor types, restricted the analysis to compare the mesenchymal type to classical, across 22 factors expressed in at least 25% of cells. The *GRID2*^High^ factor (scHPF_10) was excluded due to expression in only 8.9% of cells. Models were adjusted for cell UMI count and the random effect of donor (**Figure S8D** and **Table S27**). We also compared the proportion of cells with high-versus low-expression profiles across cells from classical versus mesenchymal tumors (Chi-squared test) (**Figure S8E**). High-expression cells were defined as those with a factor score greater than the median expression + 2×MAD (based on all cells in the dataset). Lastly, to evaluate whether factors capture DEGs between mesenchymal versus classical tumor-derived myeloid cells, we applied a MAST model adjusted for UMI count (**Figure S8F**).

### Compound-treated glioblastoma slice culture (Zhao et al., 2021; Levitin et al., 2023).^20,57^

We projected 52,688 myeloid cells from glioblastoma slice cultures treated with various compounds onto the scHPF model (**Figure 2B,C** and **Figure S9**). Out of nine drug treatment conditions with at least 1,000 cells, we focused downstream analysis on seven treatments with at least three donors with 200 cells in each treated and untreated condition (ana-12, etoposide, givinostat, ispinesib, panobinostat, R04929097, topotecan) (**Figure S9A**). To identify differential expressed factors between treated and untreated cells, we fit an LMM adjusted for cell UMI count and proportion of mitochondrial gene expression, as well as the random effect of donor (**Figure S9B, C** and **Table S28**). We also compared the proportion of myeloid cells classified as low vs. high-expressing across selected factors with significant perturbation in topotecan-treated cells (**Figure S9D**). To validate observed effects at the gene level, we also fit a MAST model adjusted for UMI count, to identify DEGs between topotecan-treated and untreated cells (**Figure S9E**).

### Murine-xenografted hiPSCs (Hasselman et al., 2019).^5^

WT (*n*=14,907) and 5X (*n*=11,137) xMG transcriptomes were projected into the scHPF model (**Figure S10**). To test between 5X and WT transcriptomes, we fit a cell-level LMM across 22 factors which are expressed in at least 25% of cells (excluding *PLCG2*^high^), adjusted for cell UMI count and proportion of mitochondrial gene expression, as well as the random effect of donor (**Figure S10B** and **Table S29**). We also tested the differential abundance of microglial neighborhoods using *Milo* in a subset of 19,834 cells (downsampled to a maximum of 5,000 cells per donor; *k*-neighbors = 30, deltaFC = 3) (**Figure S10D-G**). At the gene level, we also fit a MAST model, adjusted for UMI count and sex, to identify DEGs between WT and 5X xMGs across factors of interest IFN-I response (scHPF_20), *APOE*^High^ (scHPF_11) and *GPNMB*^High^ (scHPF_26) (**Figure S10H**).

### CNS-substrate treated hiPSCs (Dolan et al., 2023).^26^

We projected data from 54,805 iMGs differentiated from human embryonic H1 stem cells treated with CNS-relevant substrates (synaptosomes, myelin debris, apoptotic neurons, synthetic Aβ fibrils; **Figure S11A-C)**. To explore the differential expression of factors across clustering-defined microglial subtypes (**Figure S11B**), we fit an LMM comparing H1-derived iMGs from each cluster to all other clusters, adjusted for UMI count, percentage of mitochondrial gene expression, and the random effect of the replicate number. In cases where the variance of the random effect was very small or 0 (i.e., causing a convergence failure), the model was refit without the random effect. *P*-values were adjusted for multiple tests across all factors and clusters using Bonferroni correction. The analysis was restricted to 22 factors which were expressed in at least 25% of cells (excluding *NPY1R*^High^ which was expressed in 21.7% of cells). For differential expression of factors across treatments (**Figure S11C**), we fit an LMM comparing treated and untreated cells, adjusted for UMI count, percentage of mitochondrial gene expression, and the random effect of the replicate number. In cases where the variance of the random effect was 0, the model was refit without the random effect (**Table S32**). *P*-values were adjusted for multiple tests across test using Bonferroni correction.

We also projected data from 41,555 iMGs from three iPSC lines from healthy individuals with *APOE* ε3/ε3 status and no *TREM2* mutations and subsequently treated with apoptotic neurons or PBS (**Figure S11D-H**). For iPSC-derived iMGs, we focused on 20 factors which are expressed in at least 25% of cells, excluding the *NPY1R*^High^ (scHPF_4), *GRID2*^High^ (scHPF_10), and *PLCG2*^High^ (scHPF_22) factors which were expressed in only 16-24% of cells (**Figure S11E**). To explore the differential expression of factors across PBS-treated and AN-treated iMGs, we fit an LMM adjusted for UMI count, percentage of mitochondrial gene expression, and the random effect of the replicate number (**Table S33**).

In addition to the iMG data from Dolan et al., 2023, we also projected 59,624 microglial nuclei from 51 frontal cortex biopsies (Brodmann Area 8 or 9) of individuals with suspected idiopathic normal pressure hydrocephalus (**Figure S13**).

### Compound-treated HMC3 cells (Haage et al., 2024).^17^

We projected 16,855 HMC3 cells across five treatment conditions (untreated, DMSO, Camptothecin, Narciclasine, Torin2) onto the scHPF model (**Figure S12A, B**). To evaluate differentially expressed factors across treated and untreated/DMSO-treated cells, we fit an LM adjusted for cell UMI count across 19 factors expressed in at least 25% of cells (**Figure S12C** and **Table S34**). The excluded factors were motility/adhesion (scHPF_9), HLA^High^/APC (scHPF_15), *CX3CR1*^High^ (scHPF_16), and C1Q^High^/phagocytic (scHPF_17) which were expressed in only 6.2-19.3% of cells.

### CRISPR-edited HMC3 cells (*unpublished*)

We projected 9,364 HMC3 cells with CRISPR-mediated perturbations across 90 target genes using Perturb-seq onto the scHPF model (**Figure S12F**). To evaluate the modulation of factors between KO and non-perturbed (NP) cells across target genes, we analyzed 20 genes with at least 30 cells per condition and 15 factors which are expressed in at least 25% of cells. The excluded factors were IL/IFN-γ signaling (scHPF_1), Glycolysis (scHPF_2), TLR/MAPK signaling (scHPF_7), motility/adhesion (scHPF_9), HLA^High^/APC (scHPF_15), *CX3CR1*^High^ (scHPF_16), C1Q^High^/Phagocytic (scHPF_17), PLCG2^High^ (scHPF_22) which were expressed in only 2.1-21.8% of cells. We fit an LMM for cell-level factor scores comparing KO versus NP cells adjusted for sequencing depth and the random effect of sgRNA guide (**Figure S12G** and **Table S36**). For models with a random effect of sgRNAs with a variance of 0, the model was refit without the random effect. *P*-values were adjusted for multiple tests across all factors and target genes using Bonferroni correction.

### Induced-transcription factor microglia-like cells (Dräger et al., 2022).^58^

We projected data from 19,834 singlets of a CRISPR i/a screen of induced-transcription factor microglia-like cells (iTF-Microglia) onto the scHPF model (**Figure S14**). To identify differentially expressed factors across knock-outs with at least 100 cells (33 genes) compared to non-targeted controls, we fit an LM adjusted for cell UMI count across 19 factors expressed in at least 25% of cells, not including glycolysis (2), HLA^High^/APC (scHPF_15), S100/TLR signaling (scHPF_23), and *CX3CR1*^High^ (scHPF_16) factors (**Figure S14B** and **Table S35**).

### Religious Orders Study (ROS)/Memory and Aging Project (MAP)

To evaluate the generalizability of our scHPF model, we projected snRNA-seq data from three cohorts of the ROS-MAP study, a combined, longitudinal, epidemiological study of aging and cognitive decline with well-characterized clinical data assessing the incidence of AD-dementia and related neuropathologies. The three snRNA-seq datasets included (1) 424 donors (hereafter referred to as *CUIMC1*)^13,59^ (**Figure 3**); (2) a 10x Multiome dataset derived from the ROS-MAP cohort (*n*=232; hereafter referred to as *CUIMC2*; *unpublished*); (3) a 425 donors (hereafter referred to as *MIT*)^12^ (**Figure S19A**). These datasets had overlapping donors; therefore, we excluded overlapping donors in CUIMC2 (*n*=212 donors; 25,150 cells) and MIT (*n*=127; 25,252 cells) cohorts.

Using CUIMC1, we first explored the overlap between factors and traditional cluster-based definition of microglial subtypes (**Figure S16A**). To identify marker genes for previously defined microglial subtypes,^59,60^ we conducted differential expression analysis using the MAST model comparing each subtype to all others. For DEGs (Bonferroni p < 0.05), we then conducted fast gene set enrichment analysis (FGSEA) using R package *fgsea*^61^ to identify scHPF factor enrichment (top 100 markers) to identity enrichment with either upregulated or downregulated genes per subtype.

To evaluate the stability of factor scores, we used an overlapping set of 235 donors between CUIMC1 and MIT, two independently-generated datasets. For donor-level, mean-aggregated factor scores, we calculated the Spearman correlation coefficient (**Figure S19C**).

Subsequently, we explored the association of donor-level, aggregated factor scores with AD-related neuropathological outcomes, including NIA-Regan diagnosis, (2) mean percent area of the cortex occupied by amyloid β across eight brain regions (*amyloid, amyloid_sqrt*), mean density of neurofibrillary tangles (*tangles, tangles_sqrt*), and cognitive decline (*cogng_demog_slope*), as well as, region-specific measures for amyloid, tangles, diffuse plaque burden using silver staining across five brain regions (*plaq_d_*\*), neuritic plaque burden using silver staining across five brain regions (*plaq_n_*\*), and neurofibrillary tangle burden using silver staining across five brain regions (*nft_\**) (**Figure 3** and **Figure S16**). We fit LMs for the association between targeted donor-level factor scores (mean-aggregated across cells) and neuropathological traits of interest across brain regions, adjusted for study, age at death, sex, and PMI (**Figure 3B, Figure S16B)**. Focusing on the primary measures of interests (pathological AD diagnosis, amyloid levels, neurofibrillary tangles levels, and cognitive decline), we fit similar LMs in CUIMC2 and MIT and subsequently conducted a mixed-effects meta-analysis (R package *metafor*), first, across the two independent cohorts, and lastly across all three cohorts, weighted by cohort size (**Figure S19D, E, Table S7**, and **Table S8**).

Lastly, to explore the effects of PMI, as observed in the discovery cohort, we fit an LM PMI and factor scores, adjusted for age, sex, and study within each cohort with a subsequent meta-analysis (**Figure S19F, Table S41, Table S42**).

We also conducted a differential abundance analysis using *Milo* to compare the abundance of microglial subpopulations (*k*-neighbors = 30) between AD and non-AD tissue, as well as tissue with high versus low a amyloid β levels or neurofibrillary tangles (**Figure S18)**. Differential abundance models were adjusted for sex, age at death, postmortem interval, and study (spatial FDR-corrected p-value < 0.1). Given that meta-neighborhood groupings are informed by differential abundance, the meta-neighborhoods were defined independently across the three models. Therefore, we also compared the neighborhood composition overlap between meta-neighborhood definitions, calculated as the proportion of shared cells, for example, comparing meta-neighborhood 1 in model 1 and meta-neighborhood 1 in model 2.

Lastly, to validate the association of factor scores between AD and non-AD tissue, we also conducted differential expression analysis at the gene level, comparing donors with high/intermediate NIA-Reagan scores (1/2) compared to those with low scores (3/4) using DESeq2 models using donor-level pseudobulk expression adjusts for sex, study, and age at death (**Figure S17F**).

### ARACNe model construction

ARACNe^25^ is a widely used method for reconstruction of regulatory networks in transcriptomic data that leverages mutual information to infer strong relationships between upstream regulators and differences in gene expression. Here, we chose to use ARACNe to investigate regulators of specific scHPF factors. To run ARACNe, scRNA-seq data must be aggregated into meta-cells to improve the depth of gene detection in individual samples used for network construction. For ARACNe meta-cell construction, we only used data used to train the 10x version 3 chemistry model described above. To begin, we constructed a *k*-nearest neighbor graph with *k*=50 using the scHPF factors for neighbor identification. We then randomly sample cells to use as the centroid for each meta-cell (we chose to construct 2,146 meta-cells aggregating 50 neighboring cells each). We normalized for total counts per cell, dividing the count for each gene by total counts without adding a pseudocount, then took the mean across each gene for all neighbors included in each meta-cell to give us the final value per gene per meta-cell.

As postprocessing of the regulatory network, we defined the directionality of regulation, either up-or down-regulation, using the derived TF mode-of-regulation (TFmode) coefficient. Finally, we calculated the Spearman rank-correlation coefficient between the TF and target log-normalized transcript counts. For the full network, see **Table S9**.

### Discovery ARACNe network annotation

We annotated regulons for enriched DEGs associated with AD-related traits and GO biological processes. Regulons by enrichment for DEGs associated with AD-related traits in bulk RNAseq data using a similar FGSEA approach as described for factor annotation. DEGs across AD-related traits with an FDR-adjusted p-value less than 0.05 were tested. Biological pathway enrichment across predicted TF targets was performed using enrichGO from clusterProfiler for up– and down-regulated targets independently. Biological processes were reduced into overarching categories using rrvgo with using –log_10_ transformed q-values and the default similarity threshold of 0.7 (medium).

### Validation in snRNA-seq data

To validate the association of prioritized TFs (*IRF7*, *ARID5B*, *CEBPA*, *MITF*, and *PPARG*) with their targeted factors, we re-projected the three snRNA-seq datasets (CUIMC1, CUIMC2, MIT) onto the scHPF model while excluding the five TFs of interest (*IRF7*, *ARID5B*, *CEBPA*, *MITF*, and *PPARG*) to ensure independent calculation of IFN-I response (20) and *GPNMB*^High^ (26) factors. Using donor-level mean-aggregated factor scores and normalized TF expression, we fit a linear regression to test the association of TF expression adjusted for average UMI count, age, sex, study and PMI, across non-overlapping donors. The effect of TF expression was subsequently meta-analyzed across the three cohorts using a random-effect model, weighted by cohort size in using the R package *metafor*. To understand the contribution of individual TFs to variance explained in the *GPNMB*^High^ (26) factor, we conducted variance partitioning using the *R* package *vegan*^62^. At the donor level, we computed (1) the total unique contributions of the combined TFs (*ARID5B*, *CEBPA*, *MITF*, *PPARG*) adjusted for covariates (sex, age, PMI, study, sequencing depth), and (3) individual unique TF contributions adjusted for covariates, as well as the effect of the other three TFs.

We also aimed to evaluate the robustness of the discovery ARACNe regulons within the nuclear compartment; therefore, we built an independent ARACNe model in the CUIMC1 dataset. Using a similar approach for meta-cell construction as for the discovery ARACNe network, we used the 23-factor scHPF model to construct 1,604 meta-cells, mean-aggregating raw transcript counts across 50 neighboring cells. Prior to ARACNe modeling, the counts were log_2_-normalized after adding a pseudocount. The regulatory network was then reconstructed using ARACNe-AP using the same list of 1,637 human transcription factors for the discovery network. From the gene expression matrix, we first calculated a mutual information threshold (0.0214) for retaining edges (i.e., TF-target interactions) at a default *p*-value < 110^-8^. The final consolidated consensus network was computed by retaining statically significant (Bonferroni-corrected *p* < 0.05) TF-target interactions across 200 bootstraps of the input matrix. Directionality of TF-target regulation was inferred using TFmode and Spearman rank-correlation coefficient as described for the discovery network. For the nucleus-based ARACNe network, we conducted similar analysis as in the discovery network including enrichment across factors and enrichment of regulon targets with AD-related pathological traits.

### In situ transcriptomic profiling using MERSCOPE

#### Gene panel

We designed a custom probe library with 412 protein-coding genes including known marker genes to detect all cortical cell subtypes for Vizgen’s MERFISH platform, MERSCOPE. We enriched the panel to include markers for microglia, the 23 scHPF factors, and key TFs of interest (**Table S13**).

#### Tissue preparation

Postmortem fresh frozen prefrontal cortex tissue (BA9) was from four was embedded in Optimum Cutting Temperature medium (VWR, 25608-930) and sectioned on a Leica Cryostat at −20°C at 10 μm onto MERSCOPE coverslips (Vizgen, 2040003). From each donor, we obtained three non-sequential 10 μm slices, equaling 12 tissue slices in total. The tissue was then processed and imaged according to manufacturer’s instructions (Vizgen). Once adhered to the coverslip, the tissue was fixed, followed by three washes with 1× PBS. After aspiration, the tissue slide was then incubated in 70% EtOH for at least 24 h to permeabilize the tissue. After washing with Formamide Wash Buffer, the sample was incubated with the custom probe library and left to hybridize for 36-48 h. The sample was then washed and incubated at 47°C with Formamide Wash Buffer twice. The tissue was then embedded in a polyacrylamide gel matrix followed by incubation with a tissue clearing solution overnight at 37°C to remove organic material (e.g., proteins, lipids) which may contribute to autofluorescence.

#### Imaging

After achieving transparency, the tissue was washed with the Sample Prep Wash Buffer (Vizgen, 20300001) and incubated with DAPI/PolyT Staining Reagent (Vizgen, 20300021) for 15 min with agitation. The coverslip was then assembled into the imaging chamber and inserted into the microscope for imaging. Each section was imaged using the MERSCOPE 500 Gene Imaging Kit (Vizgen, 0400006) on a MERSCOPE (Vizgen). Briefly, the sample was loaded into a flow chamber connected to the MERSCOPE Instrument. First, a low-resolution mosaic was acquired at a 10x objection, then regions of interest were selected for high-resolution imaging using a 60x lens across six 1.5 μm-thick Z-planes. For high-resolution imaging, the focus was locked to the fiducial fluorescent beads on the coverslip.

#### Cell segmentation

To obtain nuclear segmentation masks, we applied the 2D nuclei segmentation model from *cellpose* v.1.0.2^63^ using Vizgen’s Post-processing Tool (VPT)^64^. DAPI images were normalized using Contrast Limited Adaptive Histogram Equalization (CLAHE) for local contrast enhancement (clip_limit=0.01, filter_size=[100, 100]). For polygons, we applied boundary simplification (simplification_tol=2px), smoothing (smoothing_radius=10px), and filtering for spurious entities (minimum_final_area=100px). Overlaps between polygon entities were harmonized, a strategy in which entities with an overlap greater than 50% are merged into a single entity, while those with less than 50% are subtracted from the larger entity. For subtracted polygons, entities with an area of at least 100px (min_final_area) were retained with a minimum distance of 1px between entities (min_final_area). Transcripts were assigned to entities only if the transcript location was within the boundary of the entity.

#### Quality control, data integration and cell type annotation

Per tissue, cell entities were retained if they met the following criteria: (1) a raw UMI count greater than 25, less than 2500, or within the middle 95% of the distribution of all counts; (2) a gene count greater than 5 or within the middle 95% of the distribution of all counts; and (3) a nuclear volume greater than 50µm, less than 2500 µm, or within the middle 95% of the distribution. Given the challenges associated with image-based cell segmentation, we aimed to also exclude cell entities which were potentially poor segmented. As such, for each cell entity, we calculated a form factor score as per the definition provided by CellProfiler of 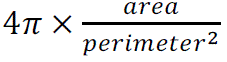 where perfectly circular objects have a form factor of 1. We excluded cell entities with a form factor below the bottom the 2% percentile.

Subsequently, data across tissues was merged and raw UMI counts were normalized using Seurat’s SCTransform adjusted for nuclear volume. Principal components were derived using RunPCA (npcs=40) after which cells across tissues integrated using RunHarmony with default parameters (lambda=1, theta=2, sigma=0.1, tau=0). To evaluate the integration of cells across tissues, we calculated Local Inverse Simpson’s Index (LISI) scores across cells.

Prior to Louvain clustering, we built a k-nearest neighbor graph using FindNeighbors (k.param=30). A UMAP was obtained using the SCT data and RunUMAP with 30 nearest neighbors. For retained clusters with more than 50 cells, we conducted cell type annotation based on differentially expressed genes (DEGs). For differential expression analysis, we used downsampled count matrices and applied a MAST model adjusted for the tissue of origin. Count matrices were downsampled using downsampleMatrix from DropletUtils to a proportion to match the tissue with the lowest median UMI count. DEGs were retained if they were expressed in at least 10% of cells and DEGs were retained if they had a log2-fold change greater than 0.25. Clusters were annotated as major cell types based on the presence of canonical marker genes among the top differentially expressed genes. Markers were selected only if they were present across all four platforms. Markers were included to identify glutamatergic neurons (*SLC17A7*), GABAergic neurons (*GAD1*), astrocytes (*ALDH1L1*), oligodendrocytes (*OLIG1*, *MOG*), OPCs (*PDGFRA*), microglia (*C1QA*), endothelial cells (*CLDN5*), pericytes (*RGS5*), CNS-associated macrophages (*LYZ*), and adaptive immunity cells (*IL7R*). We identified two clusters which were defined by a mixture of canonical markers for oligodendrocyte and glutamatergic neurons (potentially due to issues with cell segmentation). These clusters were labeled as ‘ambiguous’ and were excluded from subsequent analyses (19,704 cells).

#### Validation analyses

To demarcate gray matter and any neuronal layers, we computed spatial niches using BuildNicheAssay from Seurat v.5.1.0. To ensure that all the tissue had the same niche definitions, we arbitrarily combined the tissues into a single physical plane, separated by 2,000 µm to ensure that cells from other tissues were not included in the local neighborhood graph. Next, we built a *k*-nearest neighbor graph (*k* = 50) and used k-means clustering to define two compartments. The white matter compartment was defined through significant upregulation of oligodendrocyte markers, including *MOG*. We then attempted to define neuronal sublayers within the gray matter compartment. However, due to the limited gene panel, we did not have sufficient power to discriminate neuronal subtypes which consequently limited the ability to define neuronal layers. Nonetheless, we were able to discriminate between neuronal layer 1 as compared to deep gray matter layers 2-6. Neuronal layer 1 showed a unique enrichment of astrocytes and upregulation of *AHNAK*, *FOS*, and *ACTB*, whereas deep neuronal layers (layers 1-6) showed enrichment for *SLC17A7*, *GAS7*, and *HSP90AB1*. As such, we were able to assign microglia as being derived from one of three possible layer niches.

Using our scHPF projection pipeline, microglia were scored for each factor. Microglia were defined as ‘high-expressing’ where those with a factor score greater than the median score plus two median absolute deviations as compared to ‘low-expressing’ microglia. For each factor, we explored whether TF expression is associated with the expression of up to the 20 top factor genes. At the cell level, we fit a linear mixed-effects model for each gene expression and TF expression, adjusted for scaled UMI count and the random effect of tissue of origin. Per factor, we then meta-analyzed the effect of all top genes using a mixed-effects meta-analysis (R package *metafor*^65^). Models which failed to converge were excluded from the TF-specific meta-analysis. We chose this approach since TFs may not only regulate a factor but may also be included within the top genes of the factor. As such, scoring a factor without the TF may reduce the overall signal of the factor. A meta-analysis across multiple genes that are collectively either upregulated or downregulated by a TF may improve analytic power.

Next, we aimed to explore the distribution of IFN-I Response (20) and *GPNMB*^High^ (26) factor scores *in situ*. First, we evaluated whether factor expression is spatially autocorrelated. For both factors, we calculated the global Moran’s I^66^ with Monte-Carlo permutation testing (*nsim* =10,000) via the *R* package *Voyager*^67^ as a measure of overall spatial autocorrelation (i.e., non-random patterning) of factor expression within each tissue layer across each tissue (**Figure S27** and **Figure S31**). We restricted the analysis to tissue layers with at least 100 microglia. Next, we aimed to understand, beyond spatial autocorrelation, whether there are ‘hotspots’ of factor expression, or statistically significant local aggregations of high factor scores using the local Getis-Ord Gi*^28,29^ statistic, a Z-score of spatially-lagged values of expression at each location. Using the five nearest-neighboring microglia, we calculated the Getis-Ord Gi* for each microglia (**Figure S28**, **Figure S29A**, **Figure S32**, and **Figure S33A**), similarly restricting the analysis to tissue layers with at least 100 microglia.

Next, we aimed to understand whether microglia which highly express IFN-I Response (20) and *GPNMB*^High^ (26) factors tend to occur more proximally in physical space. First, to define layer-specific local cellular neighborhoods (including all cell types), we constructed a nearest-neighbor graph to identify the 30 nearest neighbors for each cell using Seurat’s FindNeighbors and Euclidean distance for spatial coordinates. For each microglia, we counted the number of high-expression microglia among its 30 nearest neighbors (**Figure S29B-C** and **Figure S33B-C**).

We wanted to see if non-microglia cells occurring in close proximity to IR^High^ or *GPNMB*^High^ microglia differentially express genes. To define which cells are in proximity, we used the kNN graph: cells occurring in at least one IR^High^ or *GPNMB*^High^ microglial neighborhoods were defined as occurring in a ‘high-expression’ neighborhood, while non-microglia cells were denoted as being in ‘low-expression’ neighborhoods. For each cell type, we compared gene expression between ‘high’ and ‘low’ neighborhood cells using a MAST model adjusted for UMI count (**Figure S30** and **Table S45**). The DEG analysis was restricted to cell types with at least 50 cells that were classified as being in proximity IR^High^ or *GPNMB*^High^ microglial, which excluded adaptive immunity cells.

## Data Availability

Raw scRNA-seq data (FASTQ files) generated from CD45+ cells isolated from autopsy samples and reported by Tuddenham et al., 2024 are available on GEO (https://www.ncbi.nlm.nih.gov/geo/query/acc.cgi?acc=GSE204702) and Synapse (https://www.synapse.org/Synapse:syn61001870). Newly generated raw scRNA-seq data (FASTQ files) generated from CD45+ cells isolated from autopsy samples will be made available on Synapse. ScHPF factor scores for projected datasets, raw and processed MERFISH data will also be made available on Synapse.

## Acknowledgements

We thank the individuals and their families who donated the brain samples used in this project, as well as the study investigators and staff who steward the data.

The work was supported by the Chan-Zuckerberg Initiative’s Neurodegeneration Challenge Network grant CS-02018-191971. Research reported in this publication was supported by fellowship funding from the Canadian Institutes of Health Research under award number MFE-171325 and the National Cancer Institute of the National Institutes of Health (NIH) under award number F30CA261090. Some of the work emerged from support from NIH and the National Institute of Aging (NIA) grants R01 AG070438, U01 AG061356, RF1 AG057473, R01AG048015, as well as through support from National Institute of Neurological Disorders and Stroke (NINDS) grant number R01NS103473. In part, study data was provided by the Columbia University Alzheimer’s Disease Research Center which was supported by the NIA/NIH under award number P30AG066462. In part, study data was provided by the Columbia University Parkinsonism Brain Bank which was supported by the Parkinson’s Foundation. In part, study data was provided by the National Multiple Sclerosis Brain Bank at Columbia University which was funded by the National Multiple Sclerosis Society under grant number PTAABP5286. In part, study data was provided by the Religious Orders Study and Rush Memory and Aging Project (ROS-MAP) cohort at Rush University Medical Center. ROS-MAP is supported by P30AG10161, P30AG72975, R01AG15819, R01AG17917, U01AG46152, U01AG61356. ROS-MAP resources can be requested at https://www.radc.rush.edu. In part, study data was provided by the Adult Changes in Thought (ACT) study which was funded by the NIA/NIH under grants U19AG066567 and U01AG006781. To learn more about ACT, visit https://actagingstudy.org/. We also would like to thank Dr. Phyllis L. Faust at the Department of Pathology and Cell Biology at Columbia University and Dr. Elan D. Louis at the Department of Neurology at the at UT Southwestern Medical Center for proving tissue samples of essential tremor.

All statements in this report, including its findings and conclusions, are solely the responsibility of the authors and do not necessarily represent the views of the National Institute on Aging or the National Institutes of Health.

All figures were created using Inkscape (https://inkscape.org/). Figure 1A Figure 2A, and Figure 5A, Figure S20A, C, and F were created using BioRender (https://www.biorender.com/).

## Author Contributions

Conceptualization: V.S.M., J.F.T., P.L.D.

Methodology: V.S.M., J.F.T., P.L.D. Software: H.K., P.A.S.

Validation: V.S.M., K.C., R.C., V.C.H., Y.M., M.T., Y.Z.

Formal Analysis: V.S.M., J.F.T., P.A.S.

Sample Resources: N.A.S., R.N.A., A.F.T., P.C., C.S.R., D.K., J.S., D.A.B.

Data Generation: R.C., M.T., M.O., Y.Z.

Data Curation: V.S.M., J.F.T., M.F.

Original Draft: V.S.M., P.L.D.

Review & Editing: All authors Visualization: V.S.M.

Supervision: P.L.D.

Project Administration: P.L.D. Funding Acquisition: P.L.D.

## Correspondence

Correspondence and requests for materials should be addressed to Philip De Jager.

