## Supplementary Figures for "A factor-based analysis of individual human microglia uncovers regulators of an Alzheimer-related transcriptional signature"

(14) Chan Zuckerberg Biohub, New York, New York, USA.

Corresponding author (†)

Philip L. De Jager, MD PhD
Center for Translational & Computational Neuroimmunology
Department of Neurology
622 West 168th street
New York, NY 10032


### ***Supplementary Figures***

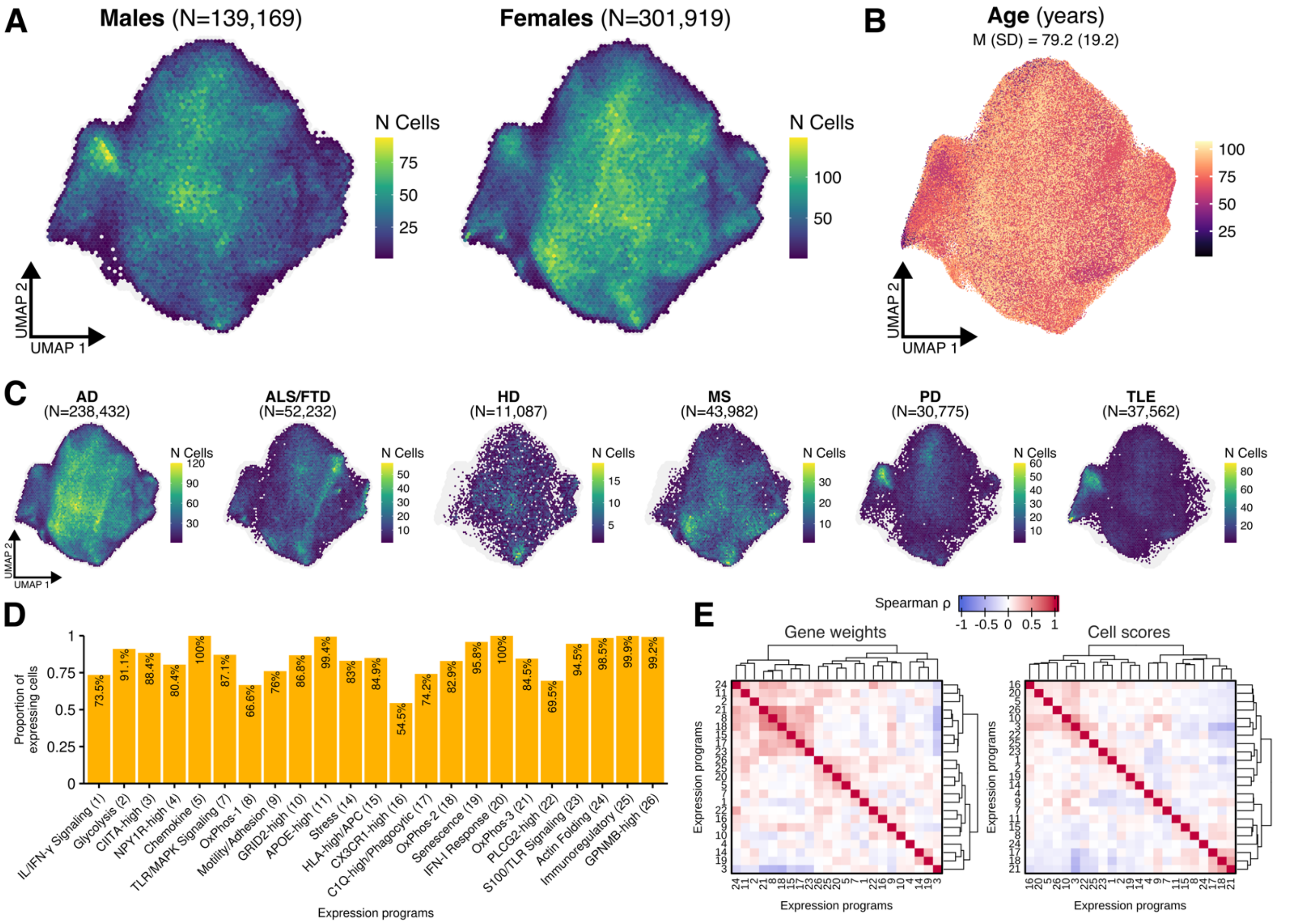

**Figure S1. Sample characterization within the scHPF model and distribution of factors.** **(A-C)** UMAP embedding showing donor metadata, including **(A)** sex, **(B)** age at death, and **(C)** diagnostic category across 441,088 microglial transcriptomes from 161 donors. *Note.* For the diagnostic category, only the most frequent diagnoses are shown (> 10,000 cells). **(D)** Proportion of microglia expressing a factor (score > 0.01). **(E)** Spearman correlation coefficient matrices of factors based on gene loadings (left panel) and cell scores (right panel).

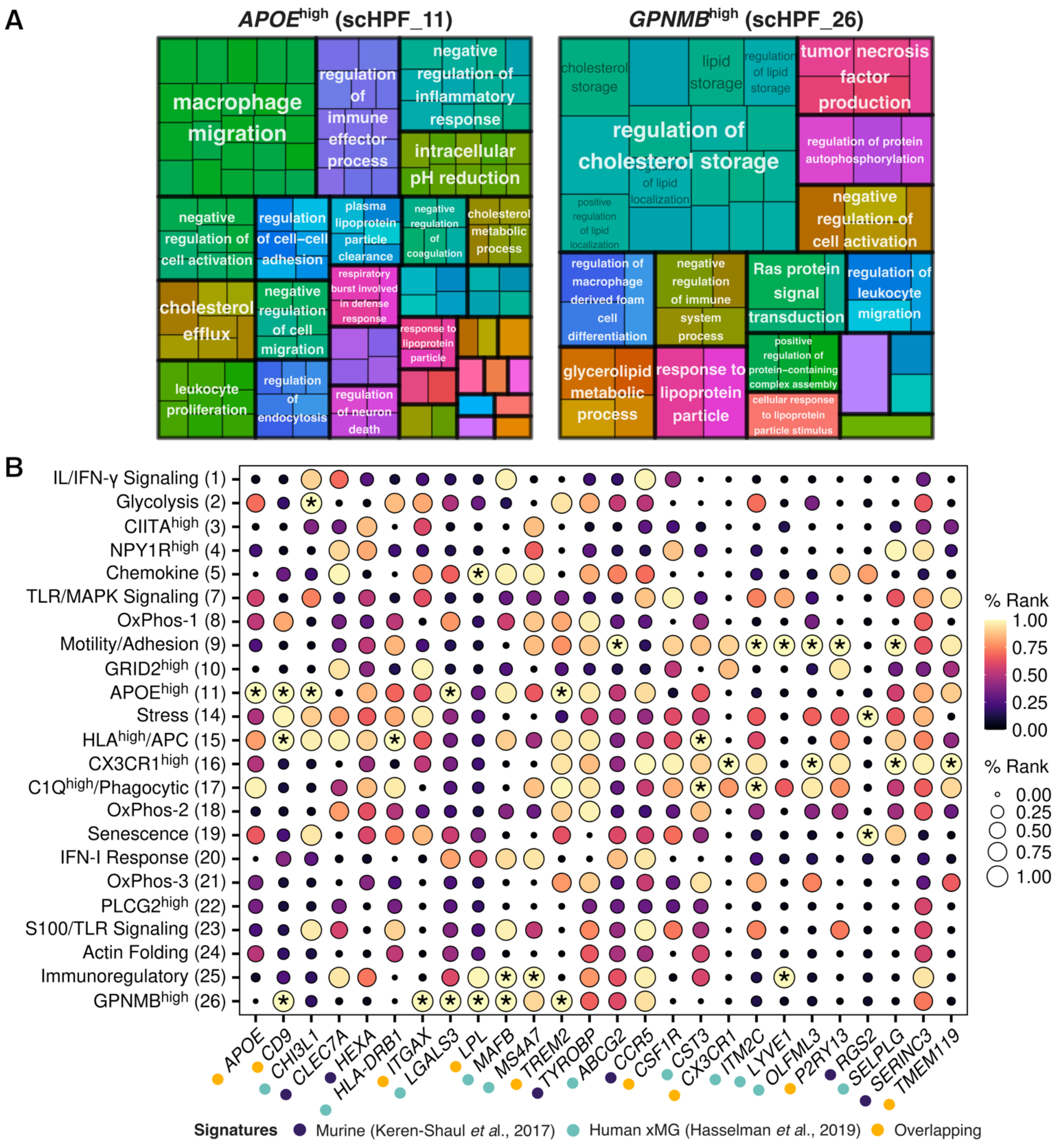

**Figure S2. Disease-associated microglia signatures across scHPF factors.**
**(A)** GO treemap diagram showing hierarchical organization of significantly enriched GO pathways across the factors with enrichment for murine-defined disease-associated microglia (DAM) genes (FDR $p$-value < 0.05 and $q$-value < 0.05). Note, the CX3CR1$^{High}$ (16) factor has no significantly enriched GO pathways at the FDR threshold. **(B)** Overlap between factor genes (y-axis) and disease-associated gene signatures (x-axis) reported in human murine-xenograft microglia (Hasselman et al., 2019) and 5xFAD murine microglia (Keren-Shaul et al., 2017). Color and size represent percentage rank of gene loading per factor, where higher percentage rank and greater size represent a greater gene loading (i.e., importance). Asterisk denotes genes that are among the top 100 genes per factor.

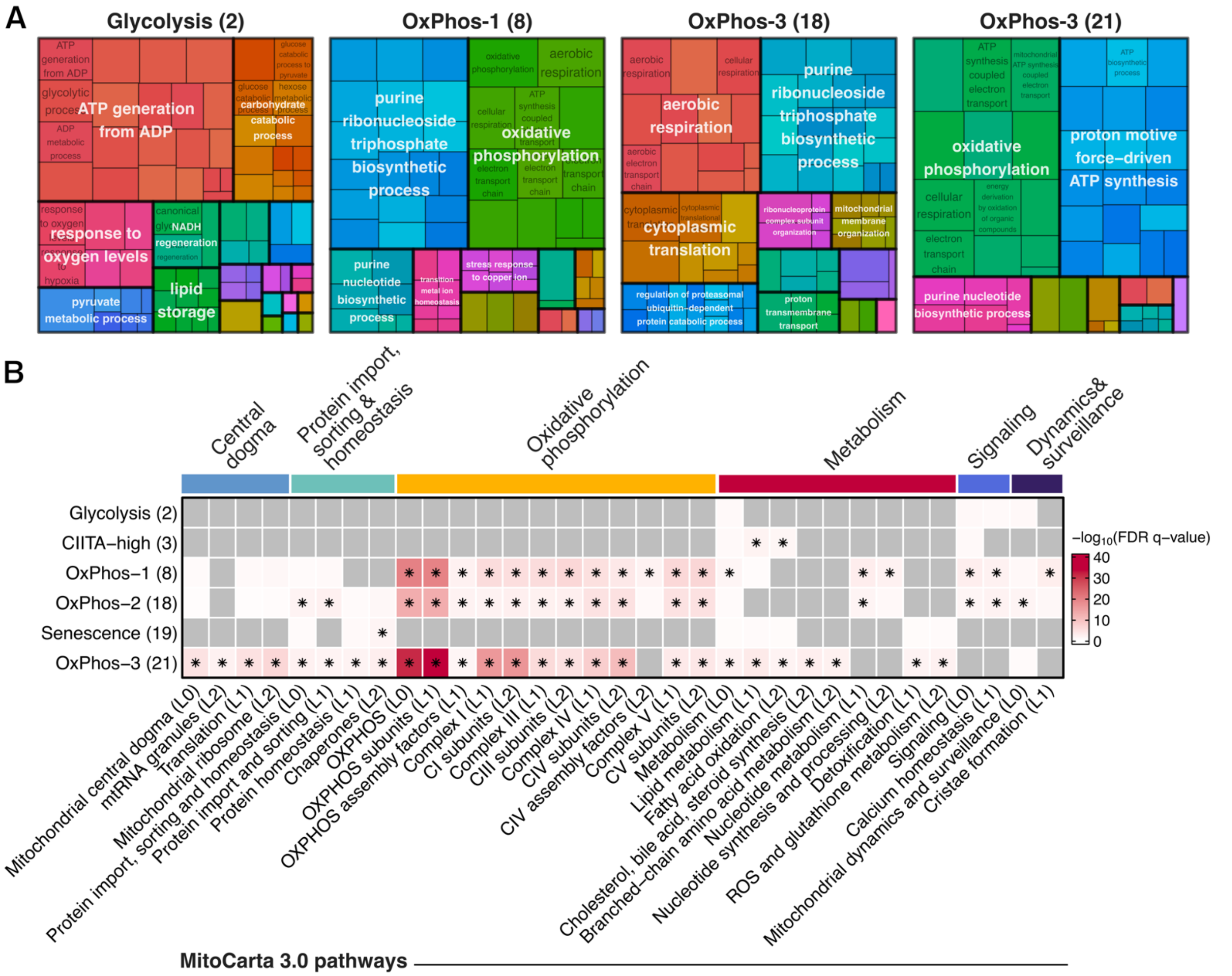

**Figure S3. Mitochondrial functional pathway enrichment across factors.**

**(A)** GO annotation for factors capturing metabolic pathways. **(B)** Heatmap showing $-\log_{10}$ FDR q-values for hypergeometric enrichment of the top-100 genes for each factor (y-axis) across MitoCarta 3.0 pathways (x-axis) (see **Table S17**). For MitoCarta pathways, hierarchy level is shown in brackets and colored bars (top) indicate the top-level pathway. Showing all significantly enriched pathways across factors (FDR p-value $< 0.05$, q-value $< 0.05$). Grey indicates failed tests due to too few overlapping genes (minGSSize = 10). Hypergeometric tests were conducted using enricher from clusterProfiler across 77 MitoCarta pathways from a universe of 18,640 possible genes (all possible scRNA-seq genes and those annotated to MitoCarta pathways).

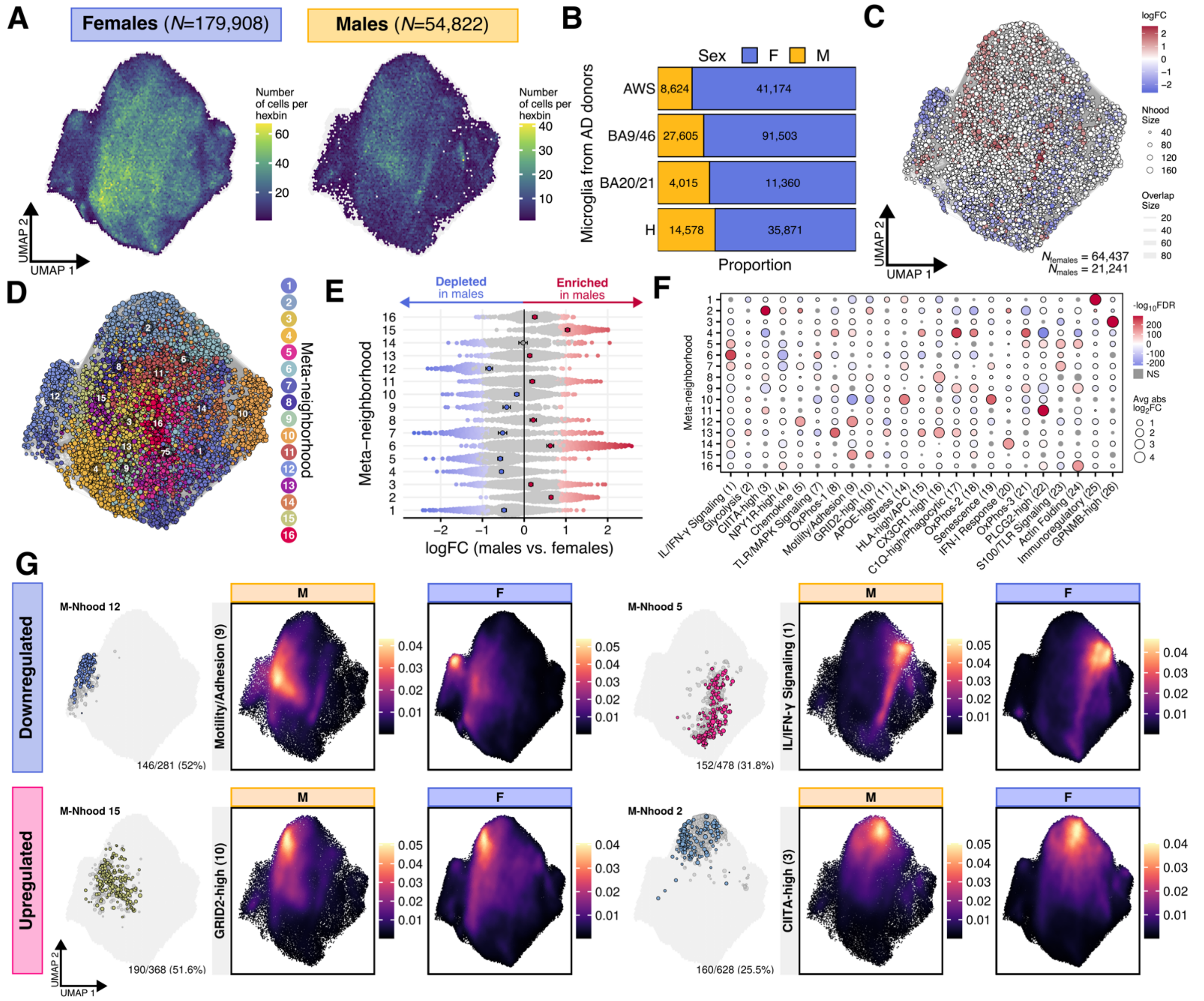

**Figure S4. Males diagnosed with AD show a depletion of motility and enrichment *GRID2*$^{\text{High}}$ signatures.**
**(A)** UMAP embedding showing cells from the AWS (anterior watershed), BA20/21 (temporal cortex), BA9/46 (frontal cortex), and H (hippocampus) derived from male and female donors diagnosed with AD. Cells represented as binned, hexagonal 'meta-cells', colored by the number of cells within a hexagonal bin (*nbins*=100). **(B)** Distribution of cell counts across males and females by brain tissue. **(C)** Abstracted KNN graph representations, colored by neighborhood differential abundance log-fold change comparing males to females. Nodes represent neighborhoods and edges indicate the number of cells commonly shared between neighborhoods. Node size represents the number of cells in the neighborhood. Colored nodes indicate significant differences in abundance associated with sex, after correction for age, brain region of origin, and site. Red indicates significant enrichment in tissue from male donors, blue indicates significant depletion (spatial FDR-corrected *p*-value < 0.05; see **Table S18**). **(D)** Abstracted KNN graph representation colored by assigned meta-neighborhood (*k*=30 nearest neighbors, max.lfc.delta = 1). **(E)** Beeswarm plot showing the distribution of log-fold change across derived meta-neighborhoods in panel D. Also showing averaged log fold-change per meta-neighborhood and 95% CI. Average values compared per meta-neighborhood vs. all others (Student's *t*-test, FDR-corrected *p*-value < 0.05). **(F)** Dotplot showing log-fold change in neighborhood factor scores, comparing all neighborhoods assigned to a single meta-neighborhood versus all others (Wilcoxon test, FDR-corrected *p*-value < 0.001). **(G)** UMAP space showing kernel density estimation across key signatures differentially employed by microglia from males and females.

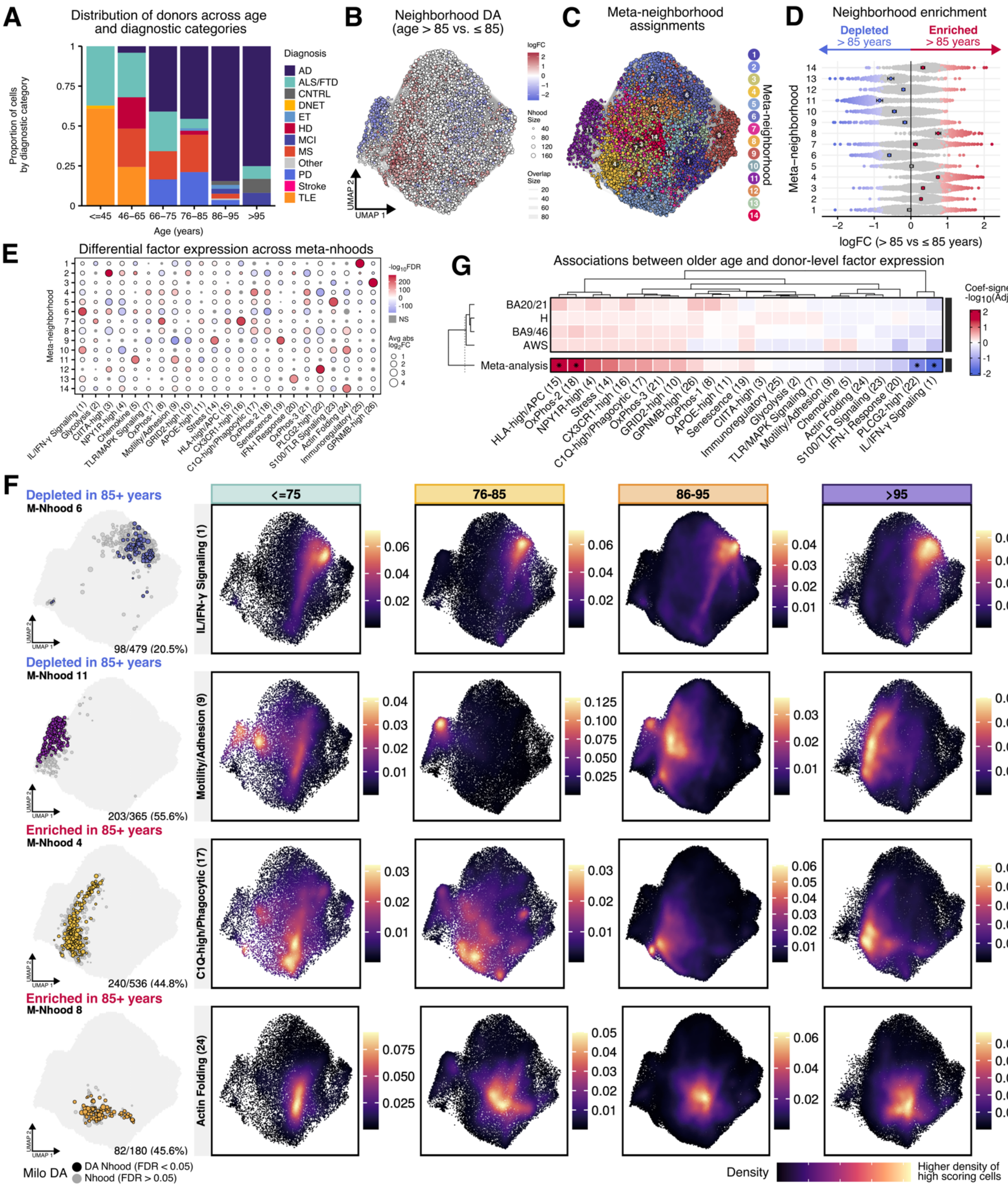

**Figure S5. Differential abundance of microglia in older age.**
**(A)** Distributions of cells across diagnostic categories. **(B)** Abstracted KNN graph representations of a subset of 87,678 microglia from AD tissue only, colored by neighborhood differential abundance comparing microglia derived from tissue of older donors versus younger donors (spatial FDR-corrected $p$-value < 0.05; see **Table S19**). **(C)** Abstracted KNN graph representations colored by meta-neighborhood assignment (max.lfc.delta = 1). **(D)** Beeswarm plot showing the distribution of log-fold change across derived meta-neighborhoods. **(E)** Dotplot showing differential factor expression across meta-neighborhoods (Wilcoxon test, FDR-corrected $p$-value < 0.001). **(F)** UMAP space showing kernel density estimation across factors defining differentially abundant meta-neighborhoods. **(G)** Heatmap showing linear regression results across brain region for the association between older age and donor-level factor scores, adjusted for sex (FDR $p$-value < 0.05; see **Table S20**).

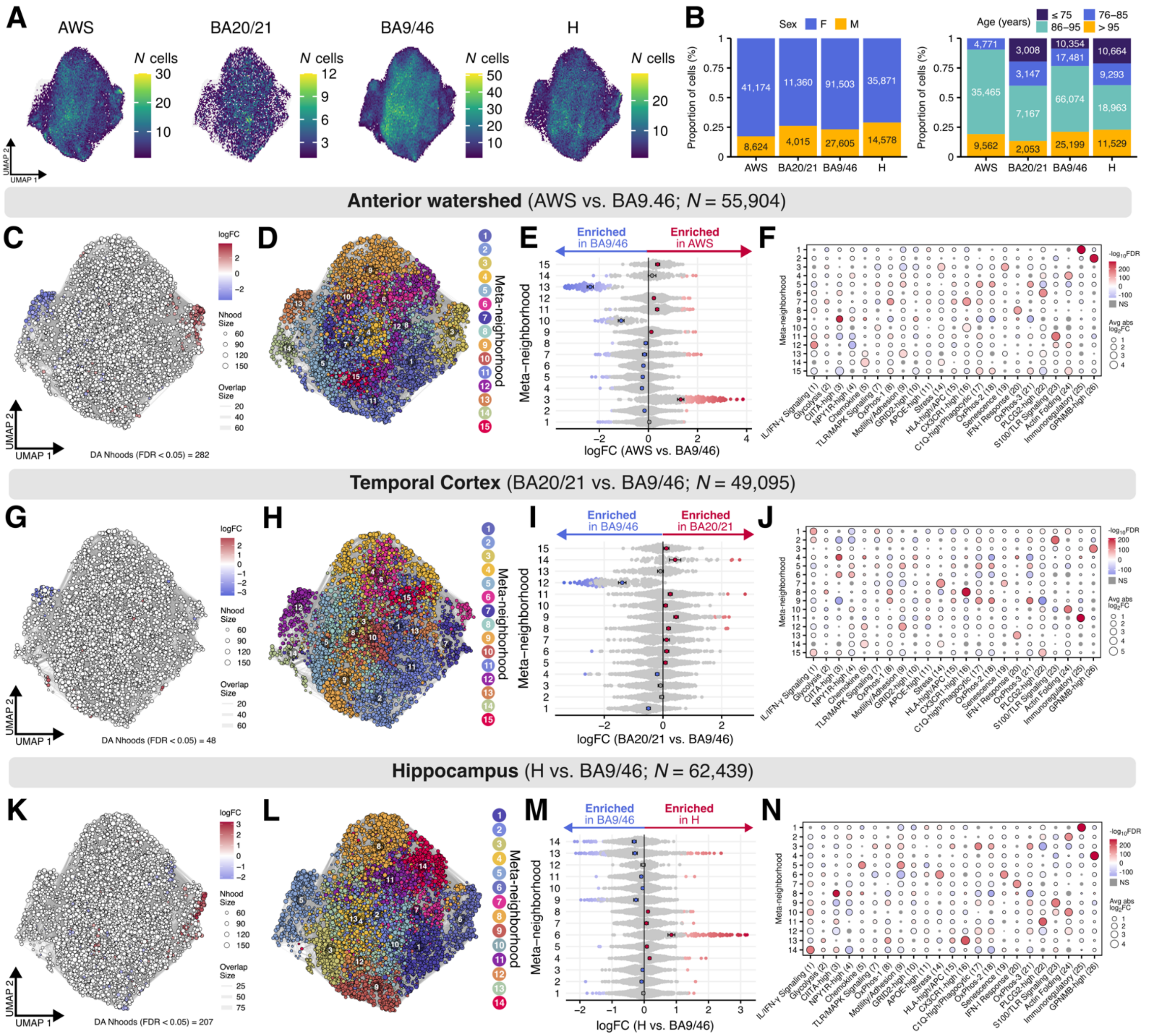

**Figure S6. In AD tissue, white matter shows enrichment microglia subpopulations highly expressing stress and senescence signatures compared to gray matter.**
**(A)** UMAP embedding showing cells derived from the anterior watershed (AWS), temporal cortex (BA20/21), frontal cortex (BA9/46), and hippocampus (H). **(B)** Distribution of cells by sex and age category for tissue of donors diagnosed with AD across the AWS, BA20/21, BA9/46, and hippocampus. **(C, G, K)** Abstracted *KNN* graph representations of a subset of microglia from tissue of donors diagnosed with pathological AD, colored by neighborhood differential abundance log-fold change compared to the BA9/46 (spatial FDR *p*-value < 0.05). Red indicates significant enrichment of neighborhood tissue from the specified regions, blue indicates significant depletion; spatial FDR p-value < 0.05; see **Table S22**, **Table S23**, **Table S24**). **(D, H, L)** Abstracted *KNN* graph representations of microglia from panel B colored by meta-neighborhood assignment (max.lfc.delta = 1). **(E, I, M)** Beeswarm plot showing the distribution of log-fold change across derived meta-neighborhoods (spatial FDR *p*-value < 0.05). **(F, J, N)** Dotplot showing log-fold change in neighborhood factor scores, comparing all neighborhoods assigned to a single meta-neighborhood versus all others (Wilcoxon test, FDR-corrected p-value < 0.001).

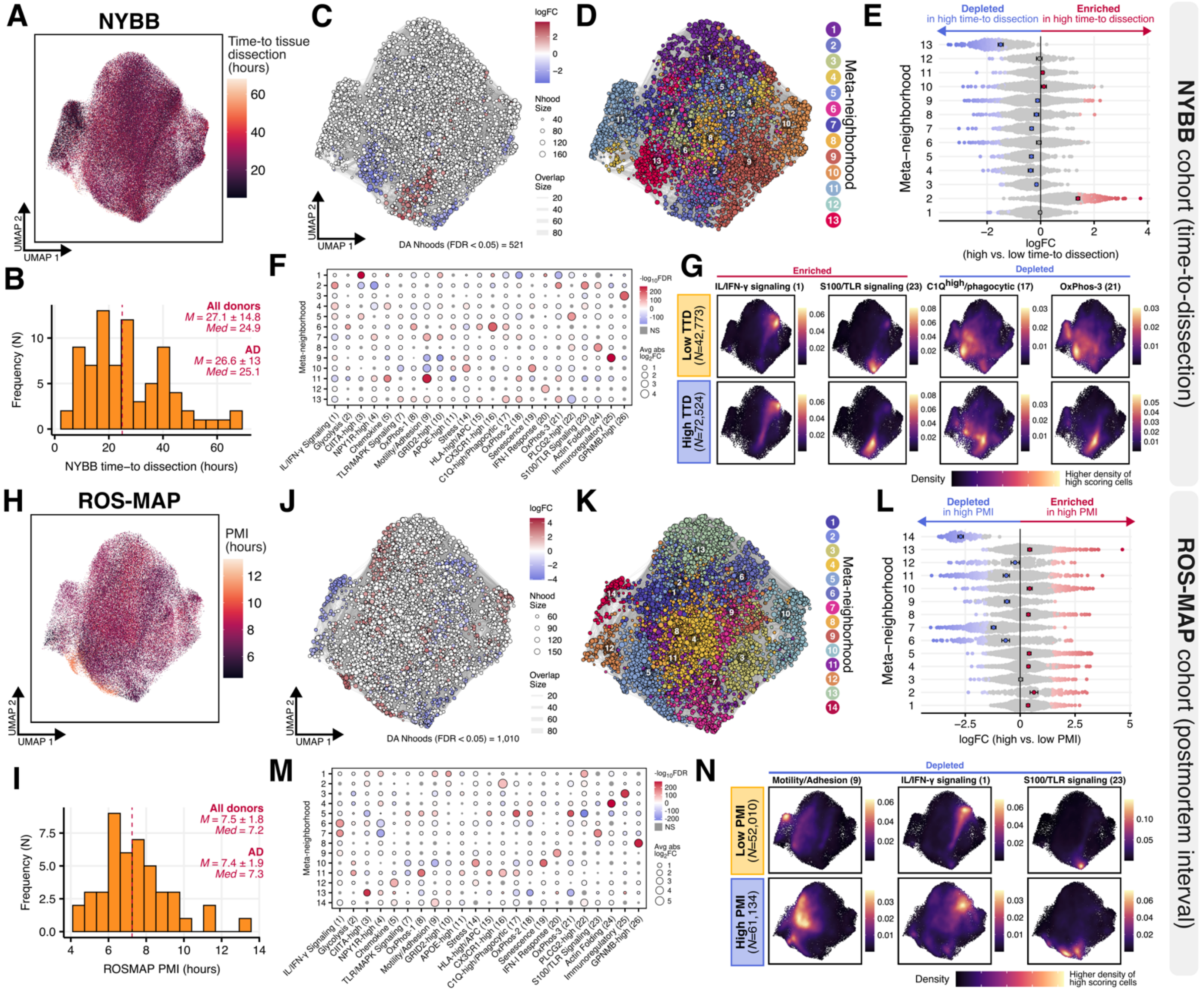

**Figure S7. Association of longer postmortem time to tissue processing with factor expression.**
**(A)** UMAP embedding showing distribution of time-to dissection in the NYBB cohort. **(B)** Histogram showing distribution of time-to dissection values in the NYBB cohort. **(C)** Abstracted *KNN* graph representation colored by neighborhood differential abundance between tissue (AD only) with higher versus lower time-to-dissection (spatial FDR p-value < 0.05; see **Table S25**). **(D)** Abstracted *KNN* graph representations colored by assigned meta-neighborhood. **(E)** Beeswarm plot showing the distribution of log-fold change across derived meta-neighborhoods (panel D) (spatial FDR-corrected *p*-value < 0.05; max.lfc.delta = 2). **(F)** Dotplot showing log-fold change in neighborhood factor scores, comparing all neighborhoods assigned to a single meta-neighborhood versus all others (Wilcoxon test, FDR *p*-value < 0.001). **(G)** UMAP space showing the density key signatures across microglia from tissue with higher vs. lower time-to-dissection in the NYBB cohort. **(H)** UMAP embedding showing distribution of PMI in the ROS-MAP subset of the discovery cohort. **(I)** Histogram showing distribution of PMI in the ROS-MAP cohort. **(J)** Abstracted *KNN* graph representations, colored by neighborhood differential abundance log-fold change comparing microglia derived from tissue higher (above the median) versus lower (below the median) postmortem interval, in AD tissue only (spatial FDR-corrected *p*-value < 0.05; max.lfc.delta = 2, see **Table S26**). **(K)** Abstracted *KNN* graph colored by assigned meta-neighborhood. **(L)** Beeswarm plot showing the distribution of log-fold change across derived meta-neighborhoods (panel K) (spatial FDR *p*-value < 0.05). **(M)** Dotplot showing log-fold change in neighborhood factor scores, comparing all neighborhoods assigned to a single meta-neighborhood versus all others (Wilcoxon test, FDR *p*-value < 0.001). **(N)** UMAP space showing the density of key signatures across microglia from tissues with higher vs. lower PMI in the ROSMAP cohort.

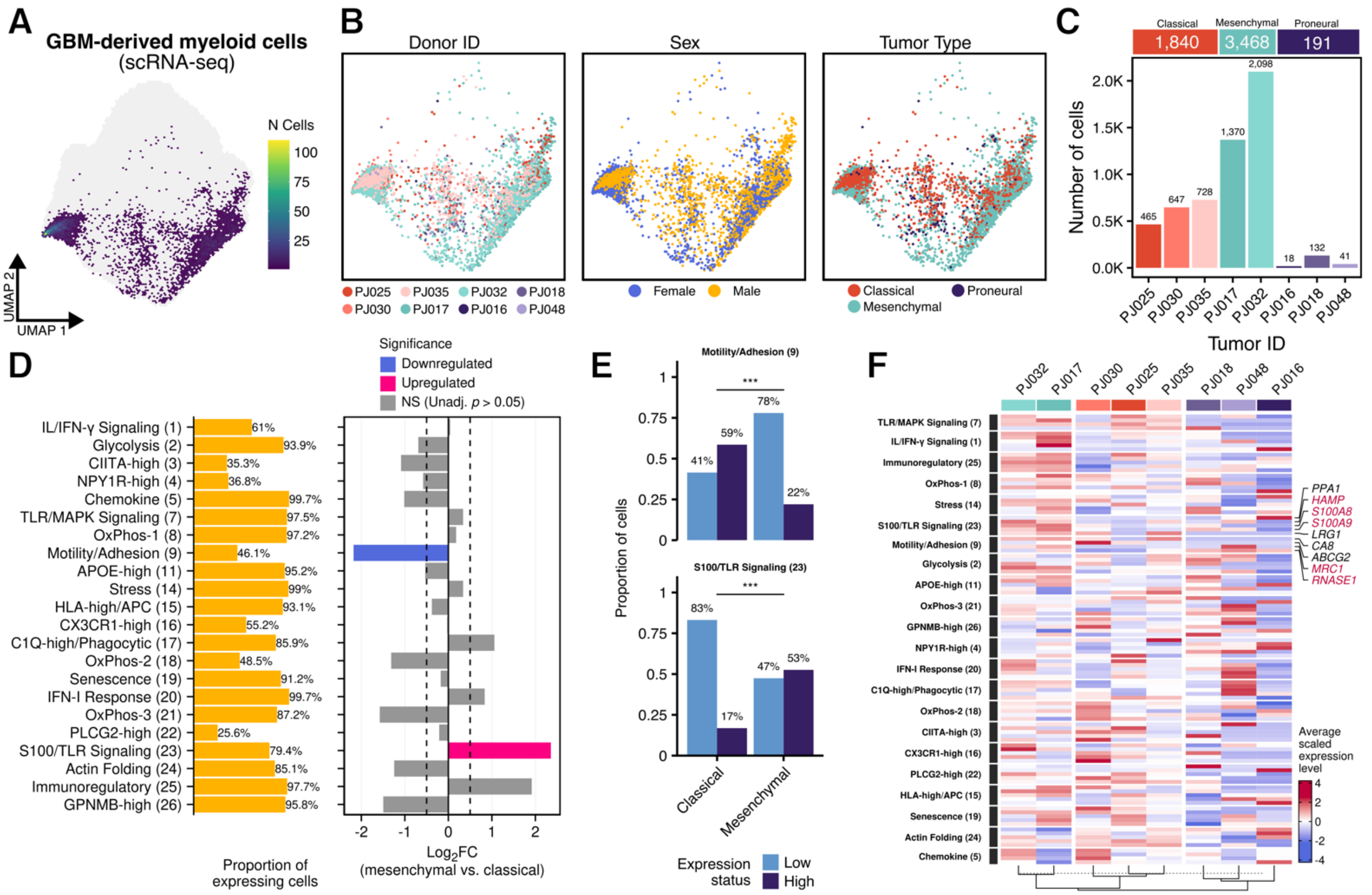

**Figure S8. Factors capture shifts in myeloid signatures from high-grade glioma.**
**(A)** Single-cell myeloid transcriptomes ($N$=5,499) from eight surgical resections of high-grade glioma within the reference UMAP space. Cells represented as binned, hexagonal 'meta-cells', colored by the number of cells (*nbins*=100). **(B)** UMAP space showing distribution of demographic and clinical variables, including glioma subtype. **(C)** Number of myeloid cells across individual tumors, colored by glioma subtype. **(D)** Differential expression of factors in myeloid cells derived from mesenchymal glioma tumors compared to classical tumors. The right panel shows the number of cells classified as 'expressing' a factor (threshold > 0.1) across classical and mesenchymal tissue. The left panel shows the differential expression of factors between cells from mesenchymal versus classical glioma (LMM model; see **Table S27**). None of the factors show a statistically significant difference (FDR $p$-value > 0.05). Color indicates factors that are differentially expressed at an uncorrected threshold $p < 0.05$. Error bars show the 95% profile likelihood confidence interval of the model coefficient. **(E)** Barplot showing proportion of myeloid cells classified as low vs. high-expressing (above median expression + 2×MAD). *Significance*: *** $p < 0.001$ (chi-squared test). **(F)** Heatmap showing average scaled expression levels of the top 10 marker genes for factors of interest across donors. Red asterisks indicate DEGs among the top markers of the S100/TLR signaling factor (scHPF_23) between mesenchymal and classical samples (MAST model, Bonferroni $p < 0.05$).

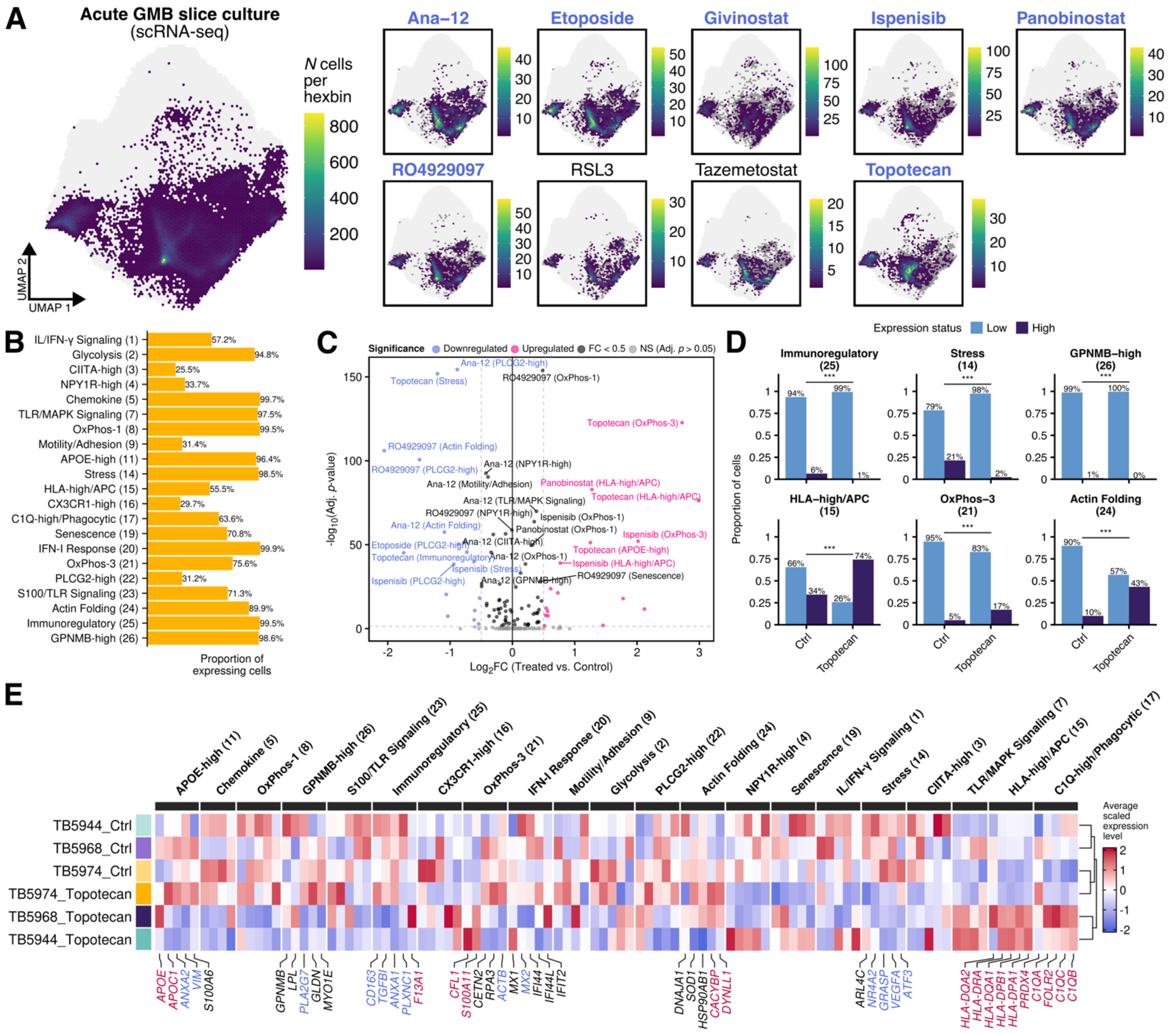

**Figure S9. GBM myeloid cells treated with topotecan show upregulated phagocytic signatures.** **(A)** Myeloid cells from glioblastoma slice cultures (N=52,688) within the reference UMAP space (light gray background). In UMAPs split by treatment condition, dark gray meta-cells show control cells from the same donors. Blue text indicates conditions included in subsequent analyses. **(B)** Barplot showing the proportion of cells 'expressing' a factor (threshold > 0.1) across all cells. **(C)** Differential expression of factors between treated and untreated cells (see also **Table S28**). **(D)** Barplot showing proportion of low vs. high-expressing cells across selected factors with significant perturbation with topotecan treatment. **(E)** Heatmap showing average scaled expression levels across factors. Color indicates significant DEGs (MAST model, Bonferroni $p < 0.05$).

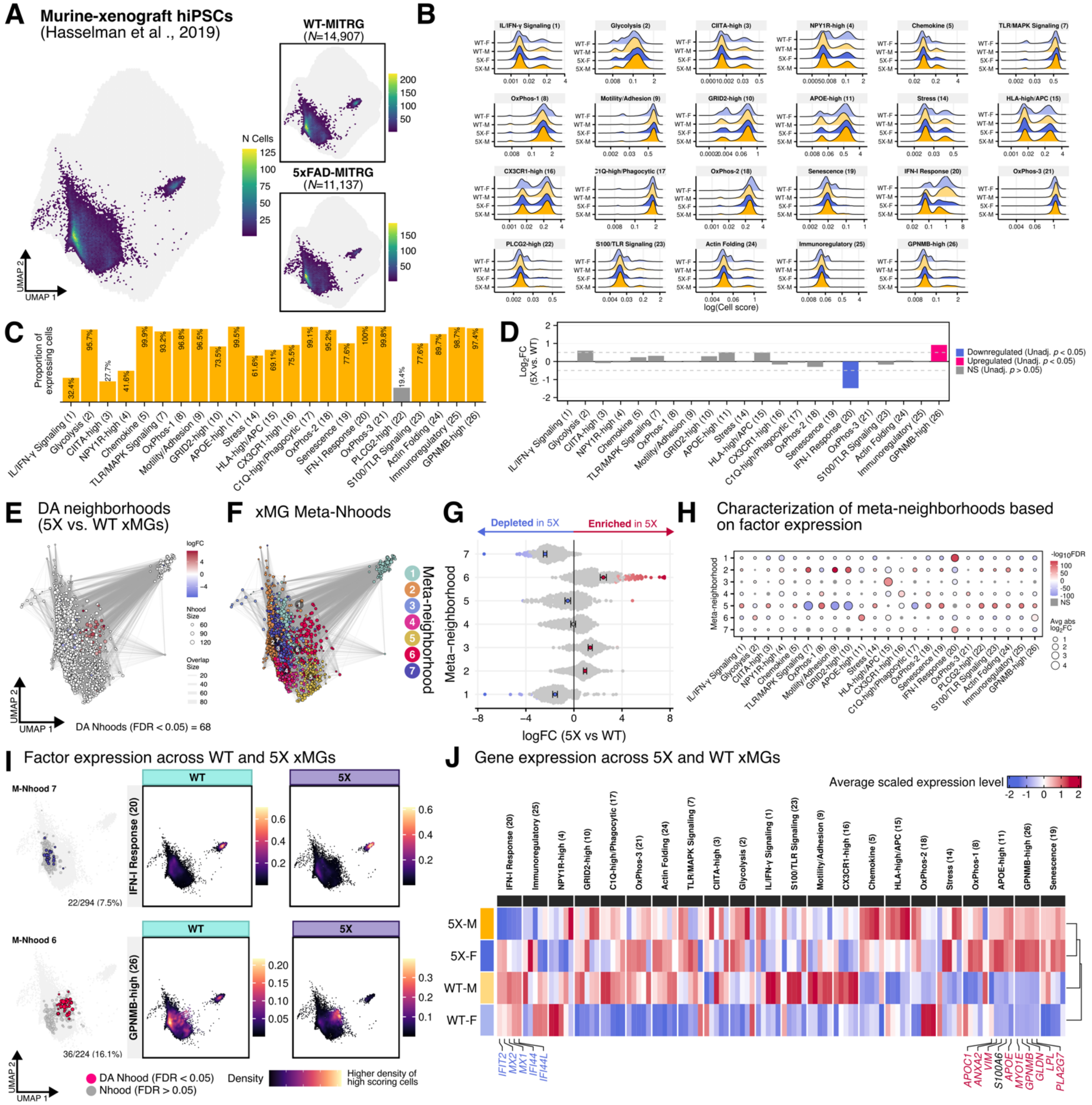

**Figure S10. Factors capture upregulation of the *GPNMB*^High signature in 5xFAD-xenografted human microglia.** **(A)** 14,907 WT and 11,137 5X xMG transcriptomes within the reference UMAP space. **(B)** Distribution of factor cell scores WT and 5X mice (dotted line shows cell score > 0.01). **(C)** Proportion of all xMG transcriptomes 'expressing' a factor (cell score > 0.01). *PLCG2*^High (scHPF_22) excluded from further analysis due to low expression. **(D)** Differential factor expression between 5X and WT xMGs. Color indicates differentially expressed factors (unadjusted $p < 0.05$; see Table S29). **(E)** Abstracted *KNN* graph representation of differentially abundant neighborhoods between 5X versus WT xMGs (spatial FDR $p$-value < 0.05, see **Table S30**). **(F)** Meta-neighborhood identity based on differential abundance. **(G)** Beeswarm plot showing the distribution of log-fold change across meta-neighborhoods (spatial FDR $p$-value < 0.05). **(H)** Dotplot showing differential expression of factors across meta-neighborhoods (Wilcoxon signed-rank test, FDR $p$-value < 0.001). **(I)** Enriched/depleted neighborhoods and corresponding factor density plots. **(J)** Heatmap showing average scaled expression of marker genes across 5X and WT mice. Colored font indicates DEGs (MAST, Bonferroni $p < 0.05$).

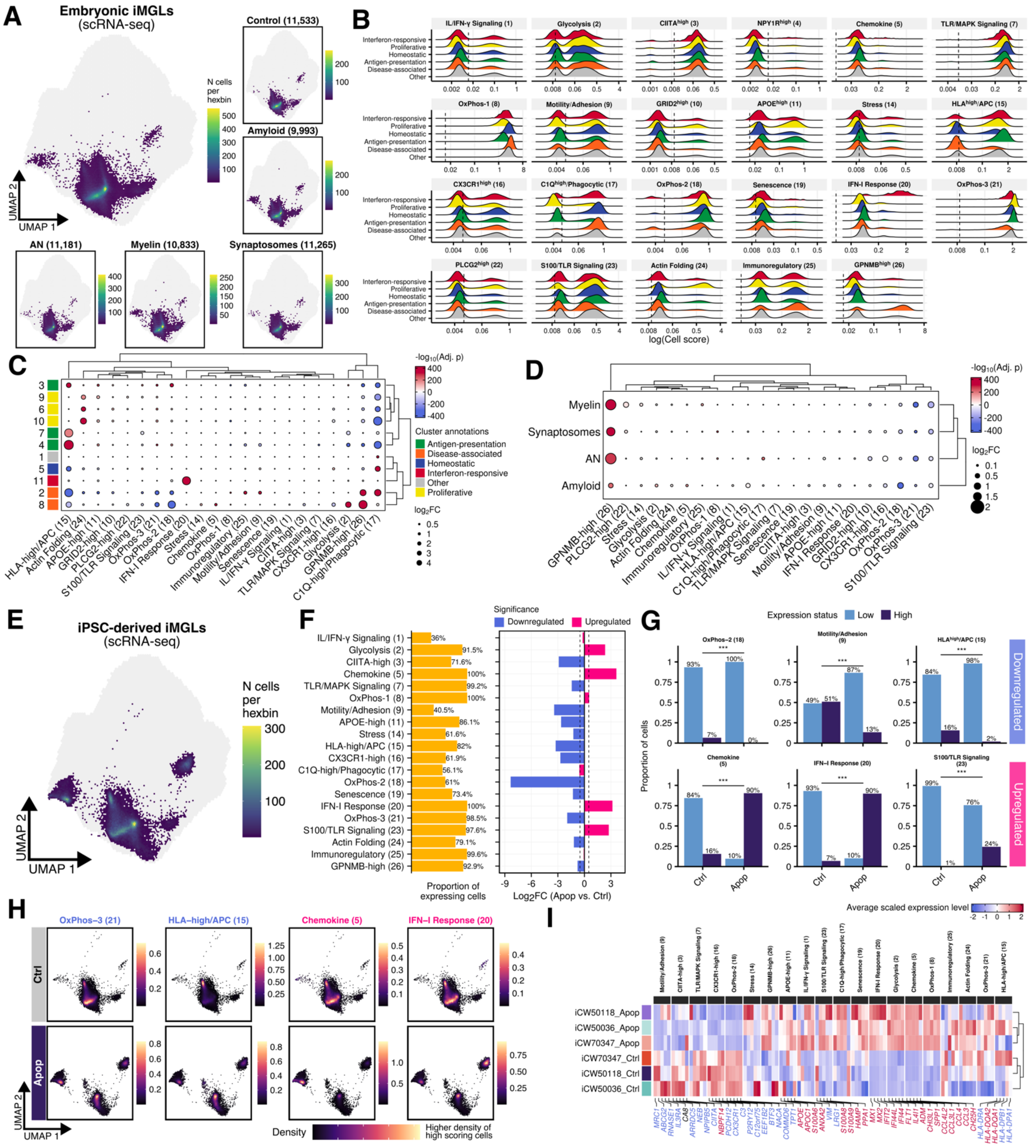

**Figure S11. Factors recapitulate signatures identified in CNS-substrate-exposed iMGs.**
**(A)** Projection of 54,805 CNS-substrate-exposed H1-derived iMGs (Dolan et al., 2023). **(B)** Distribution of factor cell scores microglia cluster annotations (dotted line shows cell score > 0.01). **(C)** Heatmap showing differential factor expression iMG cluster annotations (see **Table S31**). **(D)** Heatmap showing differential factor expression across treatment conditions (see **Table S32**). **(E)** Projection of 41,655 iPSC-derived iMGs transcriptomes (Dolan et al., 2023). **(F)** Left panel shows the proportion of cells 'expressing' a factor (cell score > 0.01) across iMGs. Right panel shows differential factor expression between AN-treated and untreated (see **Table S33**). **(G)** Barplot comparing proportion of cells classified as low vs. high-expressing across selected factors between AN-treated and untreated iPSC-derived iMGs. *Significance levels*: *** $p < 0.001$. **(H)** UMAP space showing kernel density estimation across key factors. **(I)** Heatmap showing expression levels of the top marker genes across factors. Colored font indicates DEGs (MAST, Bonferroni $p < 0.05$).

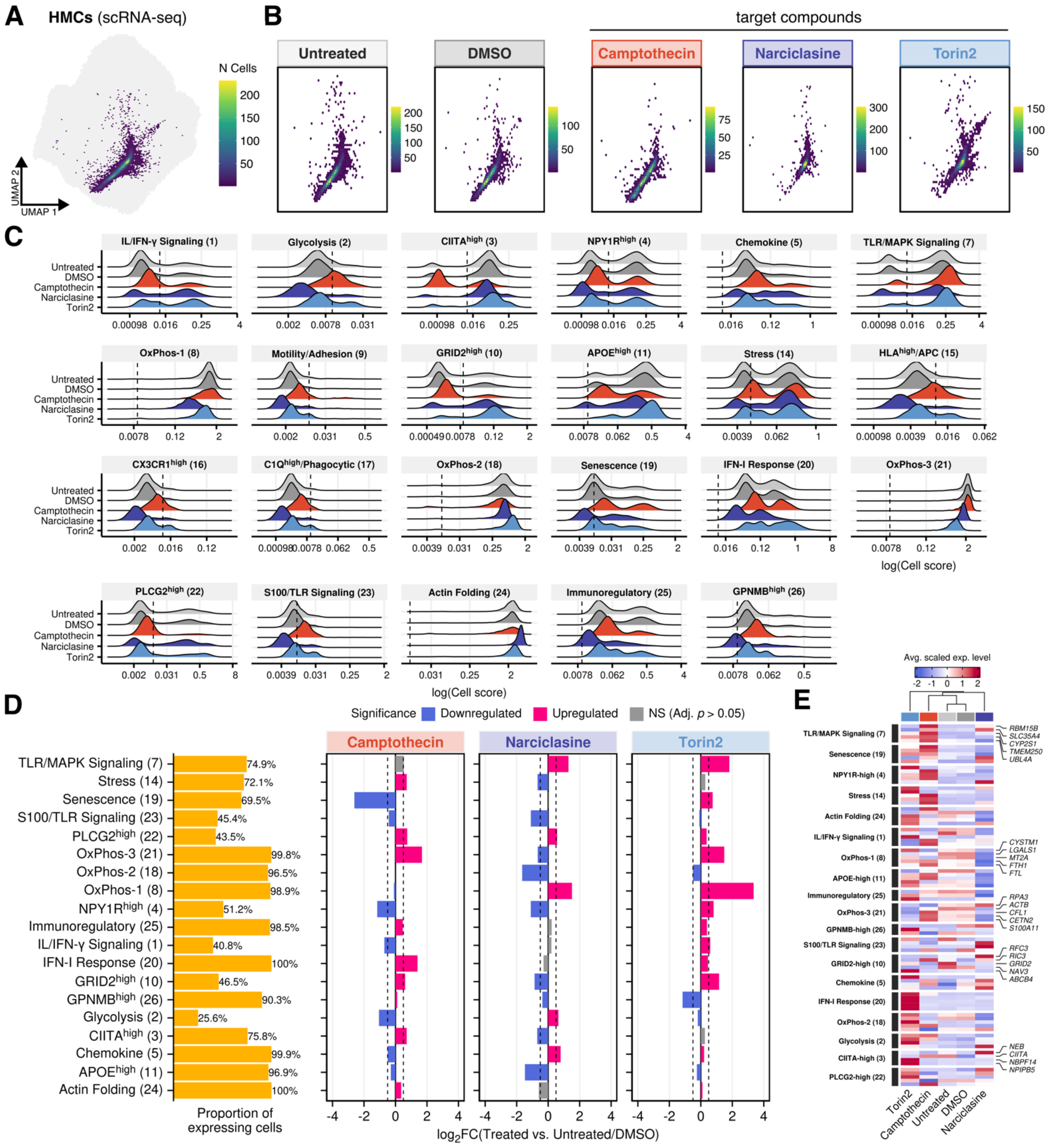

**Figure S12. Factors capture shifts in HMC3 cell signatures in response to stimulation with compounds.** **(A, B)** Projection of single-cell RNA-sequencing data from 16,855 HMC3 cells into the reference scHPF model (gray background). **(C)** Ridgeline plot showing the distribution of cell scores across factors ($log_2$ scale). Dotted line represents threshold for 'expression' (raw score > 0.1). **(D)** Barplot showing the proportion of cells 'expressing' a factor (raw score > 0.1) across all HMC3s. Left panel show differential factor expression between treated and untreated/DMSO-treated HMC3 cells (LMM; see **Table S34**). **(E)** Heatmap showing average scaled gene expression levels of marker genes across factors by treatment condition. Genes are clustered by expression level. Annotation shows the top five marker genes across top differentially expressed factors (panel D).

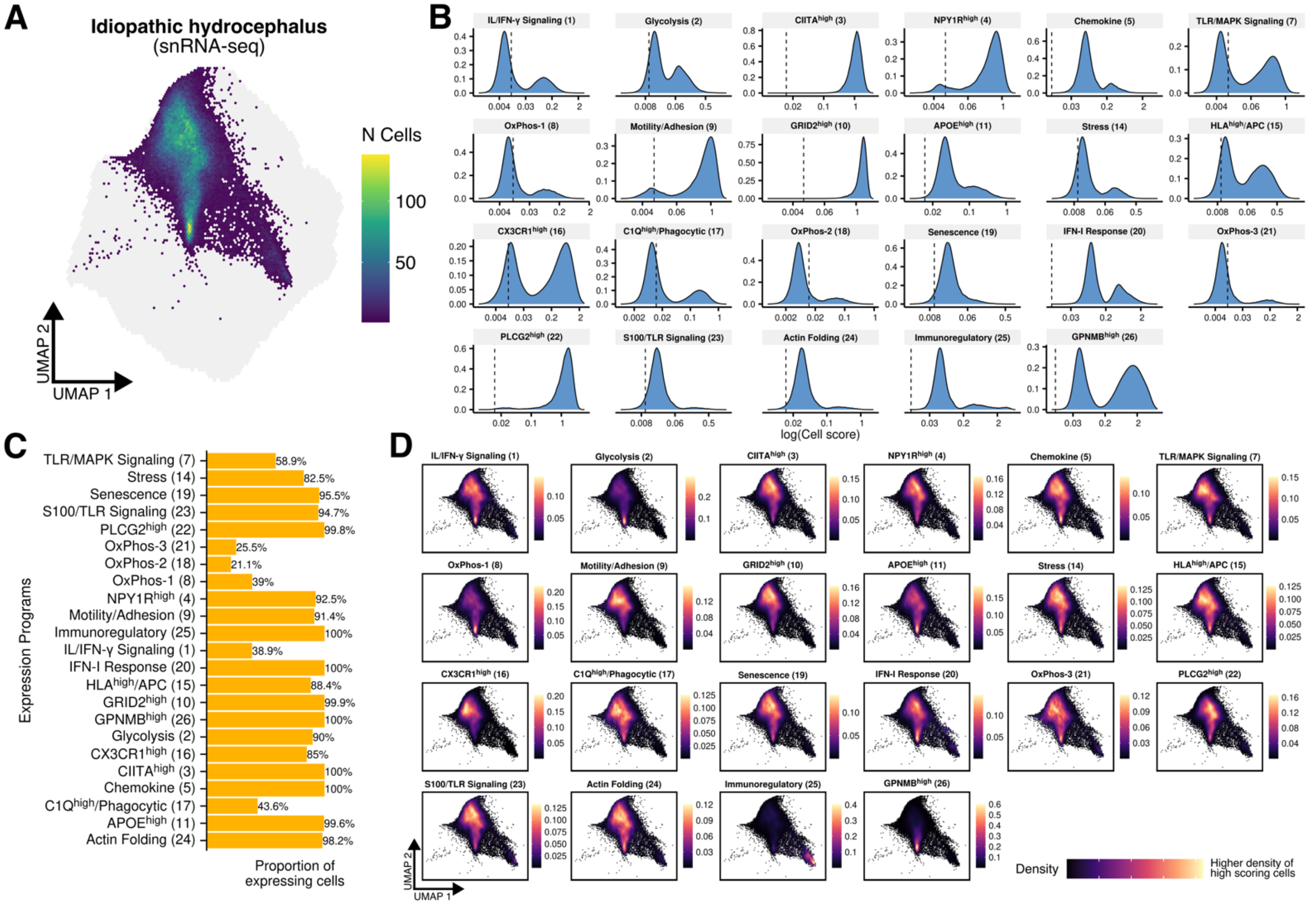

**Figure S13. Distribution of microglia factor expression in idiopathic hydrocephalus.**
**(A)** UMAP space showing the projected transcriptomes of 59,624 myeloid cells from single-nucleus RNA-sequencing of 51 frontal cortex biopsies (BA8/9) from adults with suspected idiopathic normal pressure hydrocephalus. **(B)** Barplot showing the proportion of transcriptomes 'expressing' a factor (cell score > 0.01) across 59,624 microglia nuclei. **(C)** UMAP space showing kernel density estimation across factors expressed in > 25% of cells.

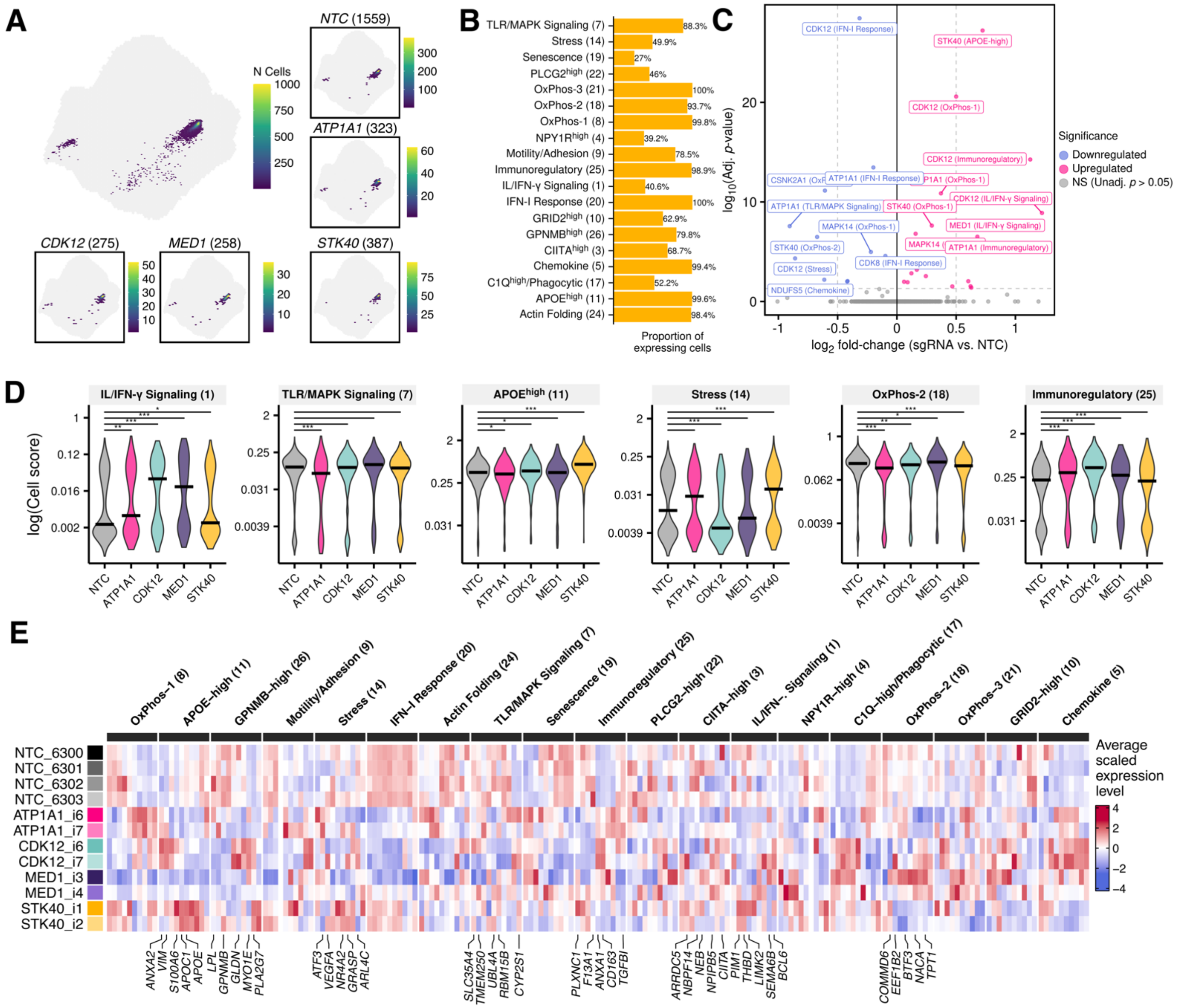

**Figure S14. Perturbation of factors in a pooled CRISPR i/a screen TF-Microglia.**
**(A)** Projection of 19,834 iPSC-derived induced-transcription factor microglia-like cells (iTF-Microglia) into the reference scHPF model (light gray background). Note, showing only iTF-Microglia classified as sgRNA singlets. **(B)** Barplot showing the proportion of iPSCs 'expressing' a factor (threshold > 0.01). **(C)** Differential factor expression between iTF-Microglia expressing targeting sgRNAs and non-targeting controls (NTCs; see also **Table S35**). **(D)** Comparison of factor scores significantly perturbed genes (Wilcoxon signed-rank test). *Significance levels*: *** $p < 0.001$, ** $p < 0.01$, * $p < 0.05$. **(E)** Heatmap showing average scaled expression levels of the top marker genes for differentially expressed factors across *ATP1A1*, *CDK12*, *MED1*, *and STK40* sgRNAs, as well as NTCs.

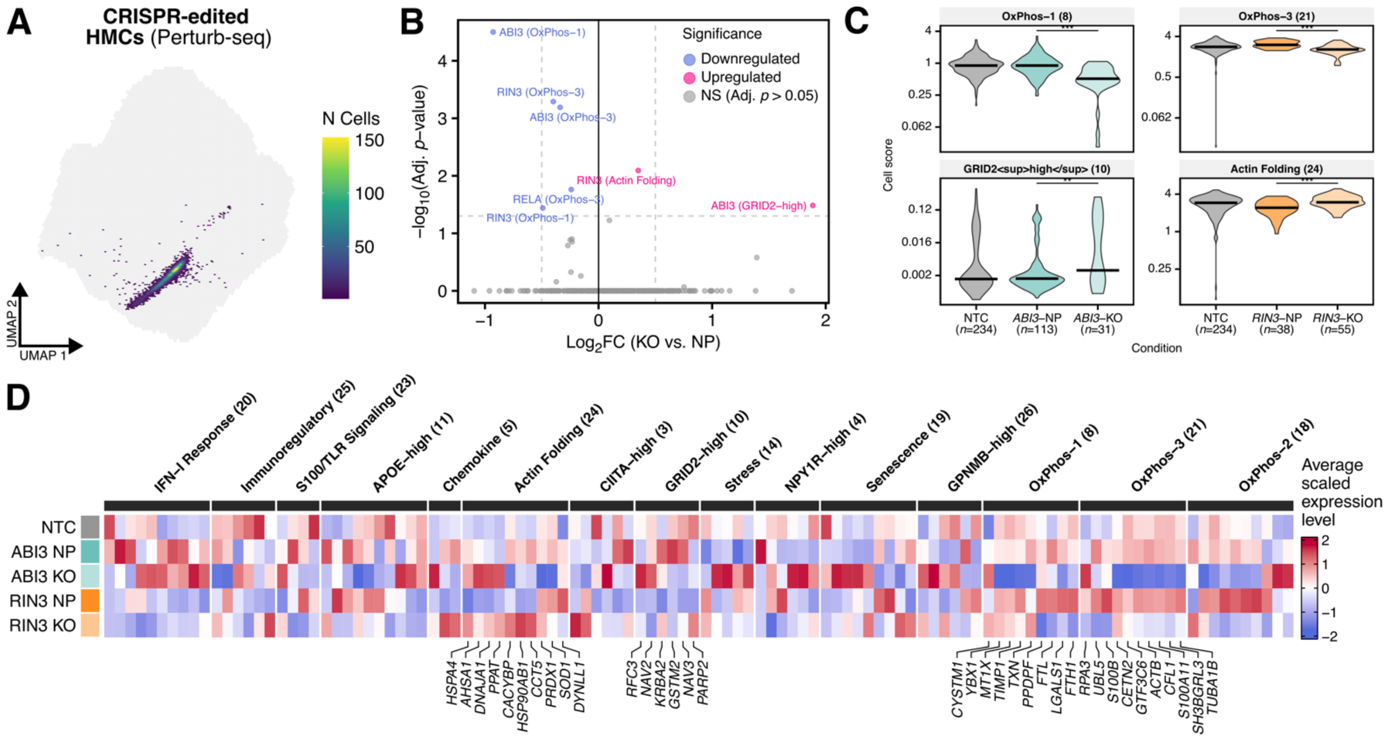

**Figure S15. Factors capture shifts in HMC3 cell signatures in response to CRISPR-mediated perturbations.** **(A)** Projection of a Perturb-seq dataset of 9,364 HMC3 cells with CRISPR-mediated perturbations into the reference scHPF model. **(B)** Modulation of factors between knock-out and non-perturbed cells (LMM model; **Table S36**). **(C)** Distribution of factors scores between non-perturbed cells and gene knock-outs (Wilcoxon signed-rank test). Significance levels: *** $p < 0.001$, ** $p < 0.01$. **(D)** Heatmap showing average scaled gene expression levels of ten marker genes across factors by knock-out conditions. Some factors have less genes due to zero expression across all conditions. Genes are clustered by expression level. Annotation shows the top ten marker genes across top differentially expressed factors (panel C).

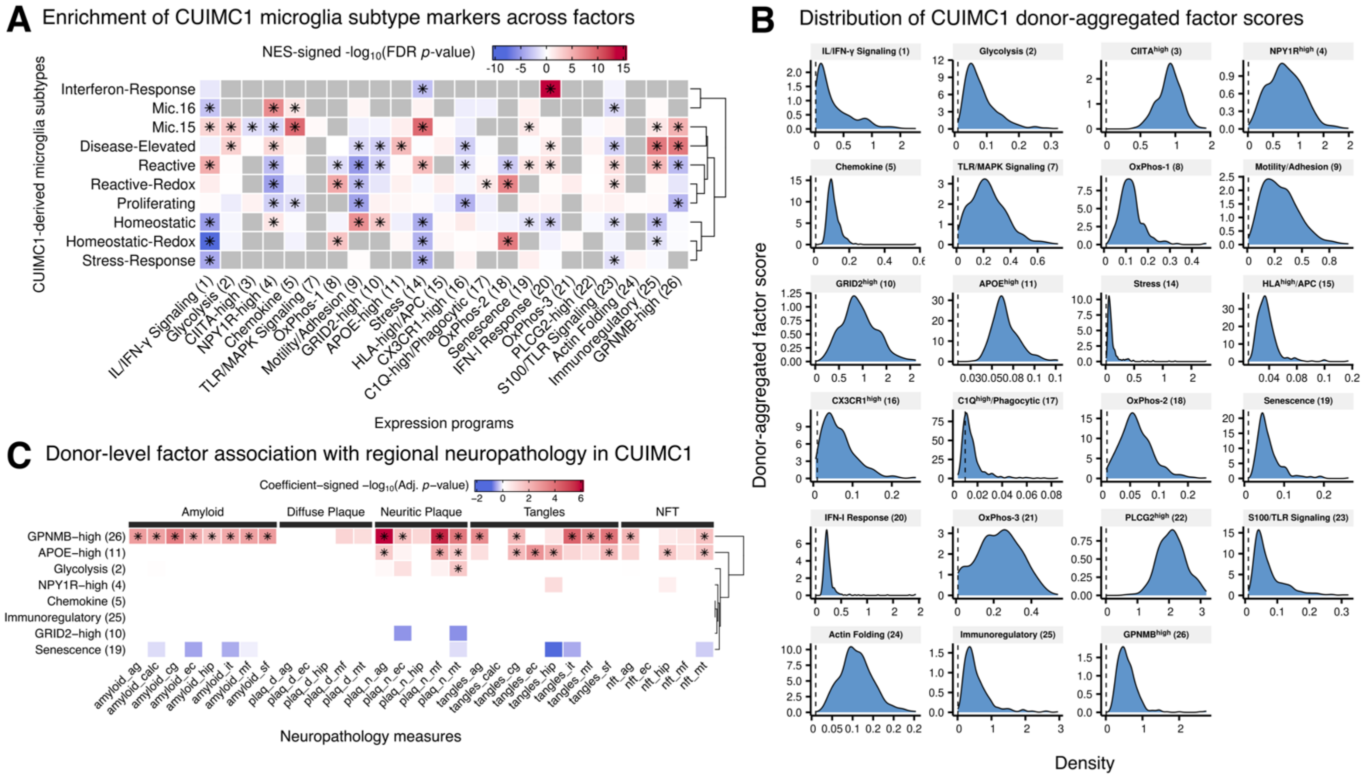

**Figure S16. Continued exploration of factor associations with AD neuropathology in the ROS-MAP CUIMC1 cohort.** **(A)** FGSEA enrichment of DEGs between clustering-derived microglia subtypes and scHPF factors (see **Table S37**). Grey indicates a failed test due to too few overlapping DEGs and factor marker genes. FDR correction for the number of factors. Significance levels: *FDR *p*-value < 0.05. **(B)** Distribution of donor-aggregated factor scores. (**C**) Linear regression results for the association between targeted donor-level factor scores and neuropathological traits of interest across brain regions (see **Table S38**). Abbreviations. ag, angular gyrus; calc, calcarine cortex; cg, anterior cingulate cortex; ec, entorhinal cortex; hip, hippocampus; it, inferior temporal cortex; mt, mid-temporal cortex; mf, mid-frontal cortex; sf, superior frontal cortex.

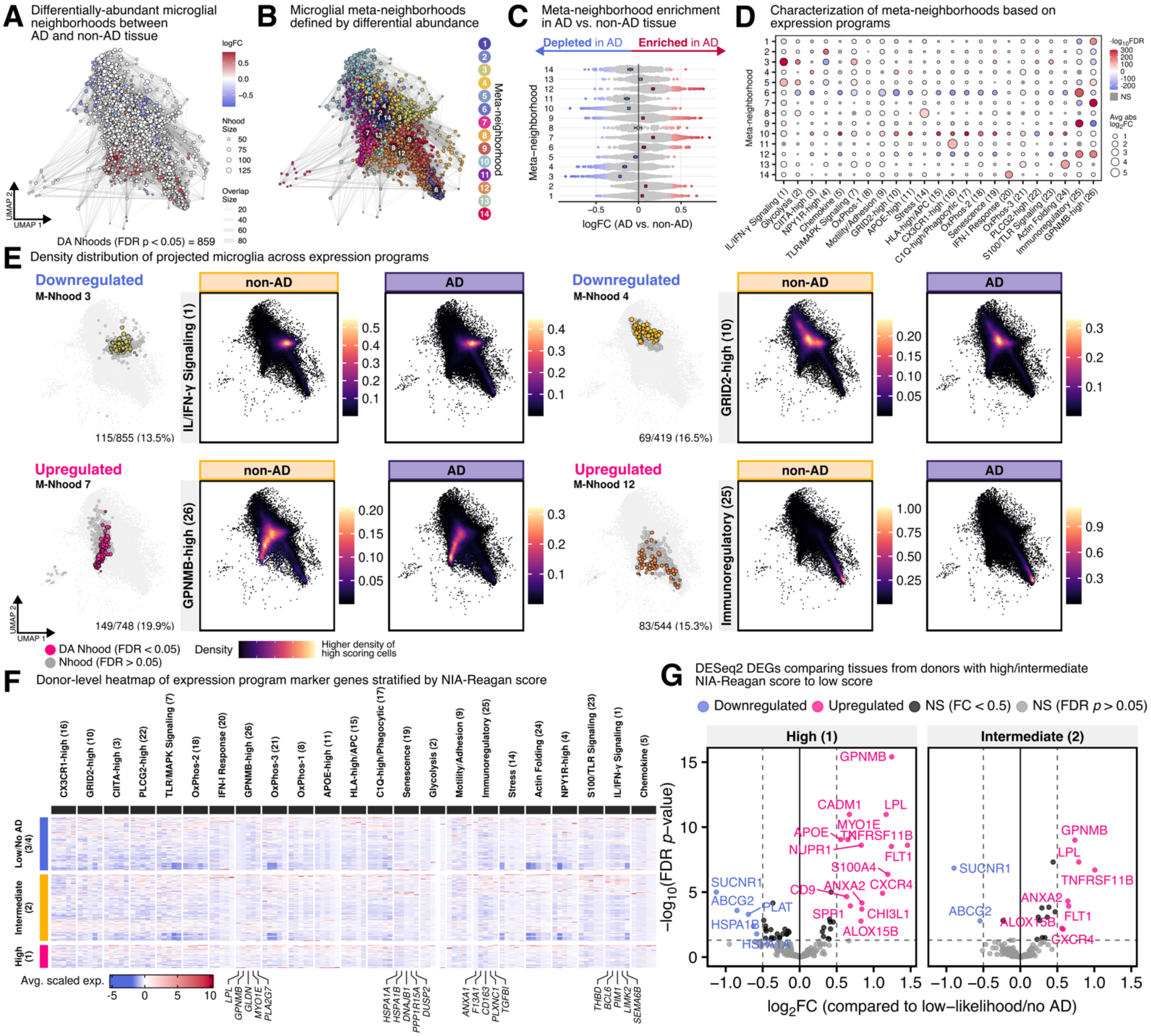

**Figure S17. Differential abundance of microglia highly-expressing GPNMB$^{High}$ (scHPF-26) across AD diagnosis.** **(A)** Abstracted KNN graph ($k$=30 nearest neighbors) representation of nuclei, colored by neighborhood differential abundance between AD to non-AD tissue (spatial FDR-adjusted p-value < 0.05; see **Table S6**). **(B)** Abstracted KNN graph representations colored by meta-neighborhood. **(C)** Beeswarm plot showing the distribution of log fold-change across derived meta-neighborhoods (spatial FDR-adjusted $p$-value < 0.1). Showing averaged log fold-change per meta-neighborhood and 95% CI. Average values compared per meta-neighborhood vs. all others (Student's t-test, FDR-corrected $p$-value < 0.05). **(D)** Dotplot showing log-fold change in neighborhood factor scores, comparing all neighborhoods assigned to a single meta-neighborhood versus all others (Wilcoxon test, FDR-corrected $p$-value < 0.001). **(E)** UMAP space showing microglial nuclei with highest factor scores using density kernel estimation. **(F)** Heatmap showing the top ten gene markers across factors of interest comparing AD pathological categories based on NIA-Reagan scores. **(G)** Differentially-expressed genes between donors with high/intermediate NIA-Reagan scores compared to those with low scores (3/4).

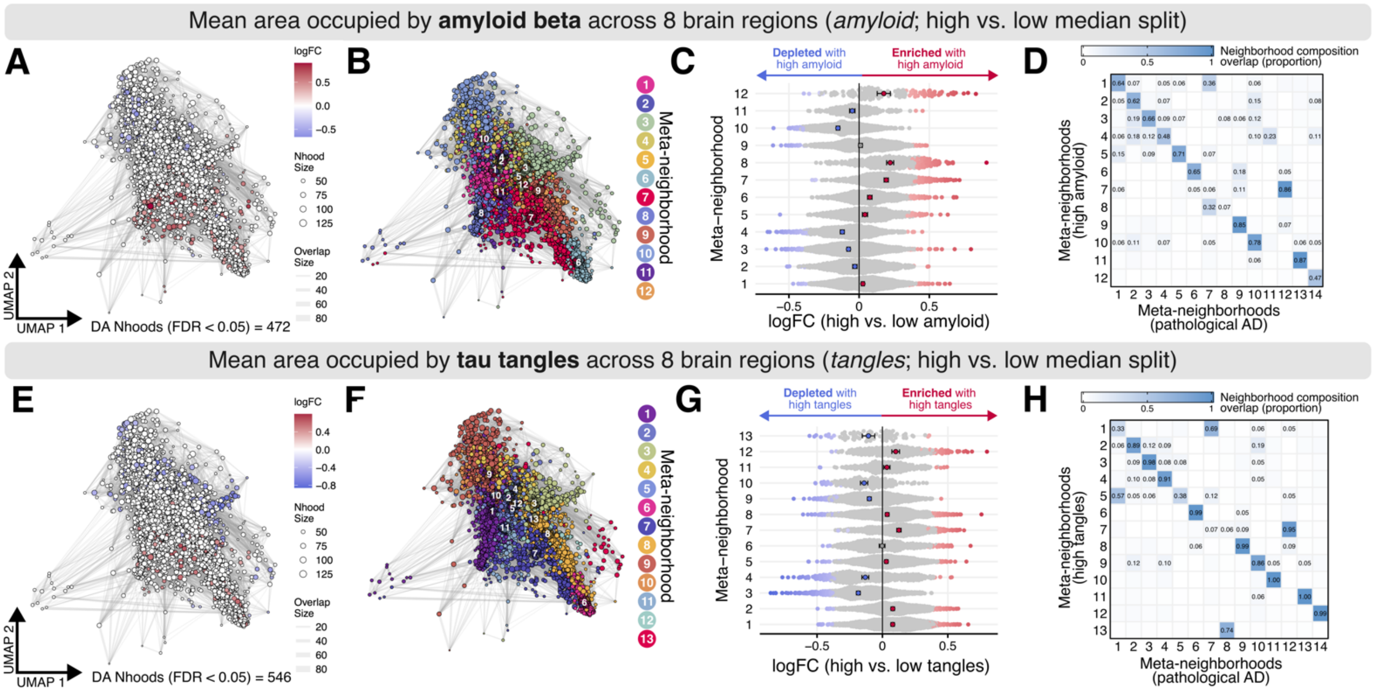

**Figure S18. Continued exploration of microglial differential abundance in the ROS-MAP CUIMC1 cohort.** **(A, E)** Abstracted KNN graph representations of the CUIMC1 cohort, colored by neighborhood differential abundance (spatial FDR $p$-value < 0.05; see **Table S39** and **Table S40**). **(B, F)** Abstracted KNN graph representations of microglia colored by meta-neighborhood assignment. **(C, G)** Beeswarm plot showing the distribution of log-fold change across derived meta-neighborhoods. Red indicates significant enrichment in those with greater specified pathology, blue indicates significant depletion; spatial FDR $p$-value < 0.05). Showing averaged log fold-change per meta-neighborhood and 95% CI (Student's t-test, FDR $p$-value < 0.05). **(D, H)** Meta-neighborhood composition similarity matrices, comparing meta-neighborhood cellular composition across analyses, using the neighborhoods defined by AD diagnosis (see Error! Reference source not found.) as a reference. Color indicates the proportion of overlapping neighborhoods (within a meta-neighborhood) calculated as the number of shared neighborhoods divided by the number of unique neighborhoods. Labels indicate proportion overlap >5%.

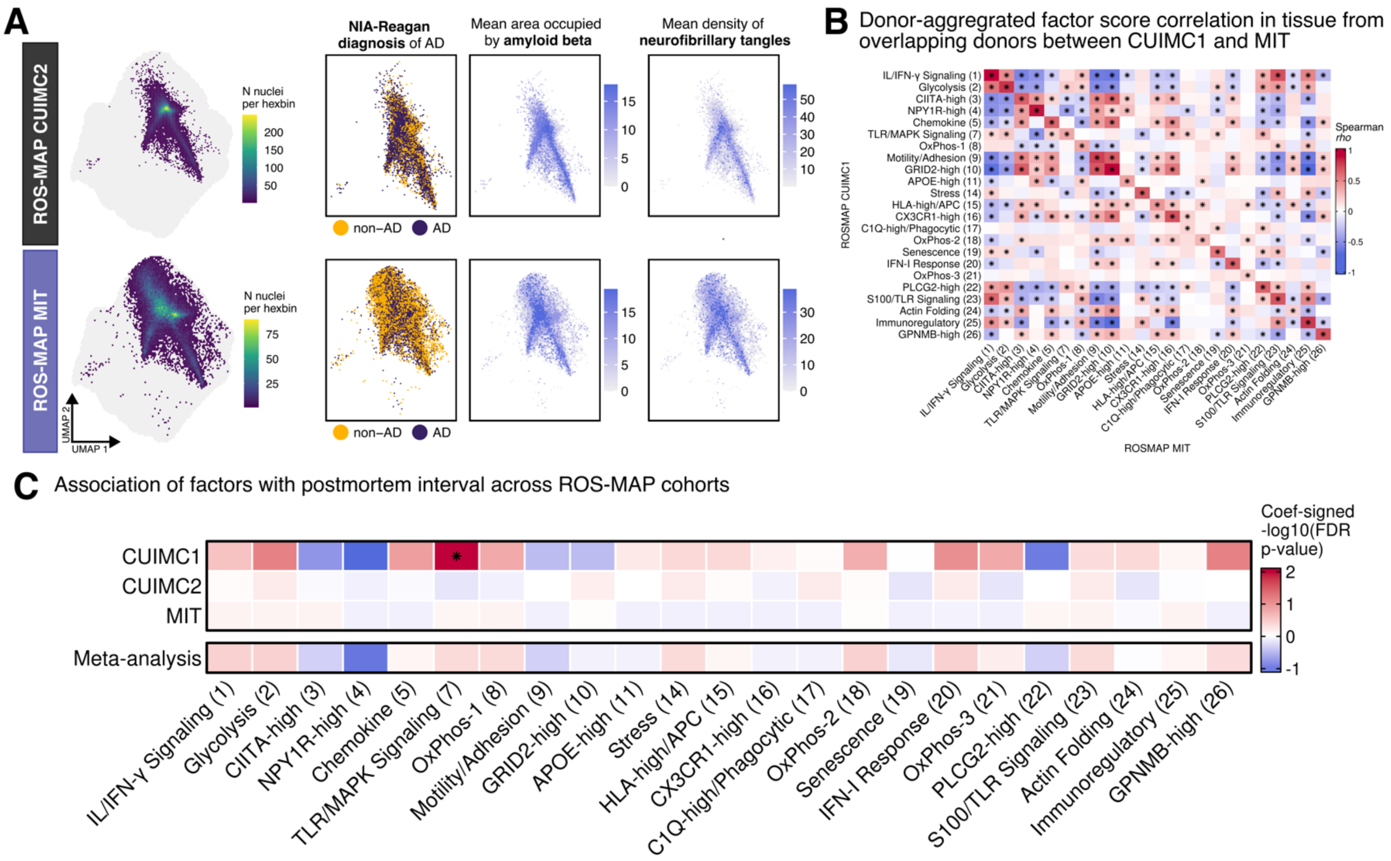

**Figure S19. Validation of factors in two independently-generated DLPFC snRNA-seq datasets.** **(A)** UMAP embedding showing projected microglial transcriptomes from an independent, ROS-MAP CUIMC2 and MIT cohorts. Adjacent plots show the distribution of pathological variables within the UMAP space, including dichotomized NIA-Reagan AD diagnosis (ad_reagan), mean area occupied by amyloid β across eight brain regions (amyloid), and mean density of neuronal neurofibrillary tangles across eight brain regions (tangles). **(B)** Correlation matrix between donor-aggregated factor scores from 235 overlapping donors from the ROS-MAP CUIMC1 and MIT cohorts. *Significance levels*: *$p < 0.001$. **(C)** Linear regression results for donor-level factor scores (mean-aggregated across cells) in association with postmortem interval, adjusted for study, age at death, and sex (see **Table S41** and **Table S42**). *Significance levels:* *FDR $p < 0.05$.

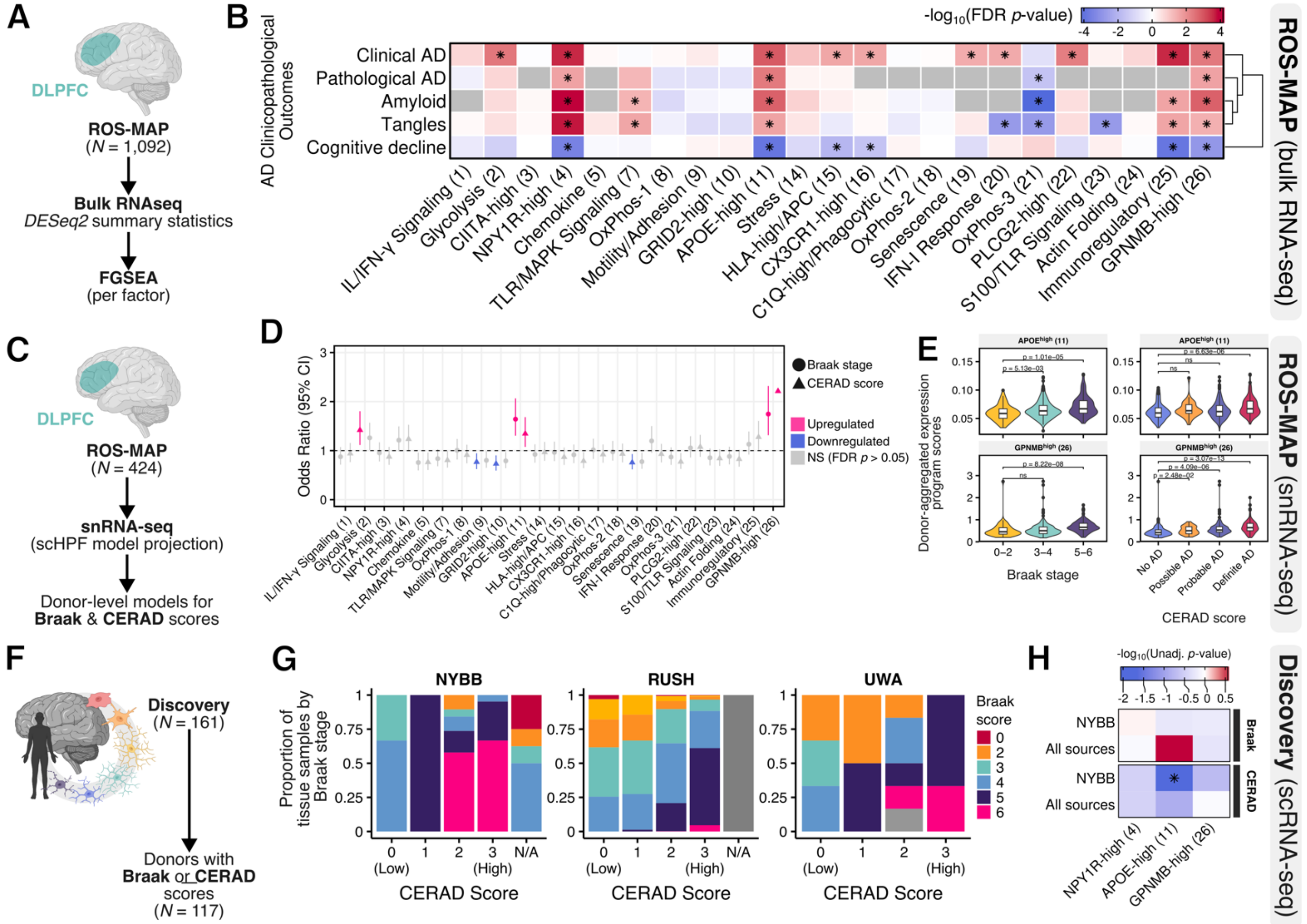

**Figure S20. DAM-like factors show positive association with AD-related amyloid and tau histopathology measures.** **(A, B)** FGSEA enrichment between factor marker genes DEGs across AD-related outcomes from bulk RNA sequencing data (18,629 genes; 1,092 DLPFC samples; see **Table S43**). FDR correction for the number of factors. Note, for cognitive decline, lower scores are worse. **(C, D)** Association results between factor scores and Braak/CERAD score in the projected CUIMC1 cohort. **(E)** Distribution of donor-aggregated factor scores for *APOE*$^{High}$ (scHPF_11) and *GPNMB*$^{High}$ (scHPF_26) across Braak stage and CERAD score categories. Significance shows Dunn's test FDR-adjusted p-value. **(F)** A subset of the discovery cohort with available neuropathological measures (Braak stage, *N*=116; CERAD score, *N*=109). **(G)** Overlap between Braak stages and CERAD scores across sites of tissue origin. **(H)** Association results between donor-aggregated factor scores and Braak/CERAD score in the discovery cohort. None of the models survived FDR correction.

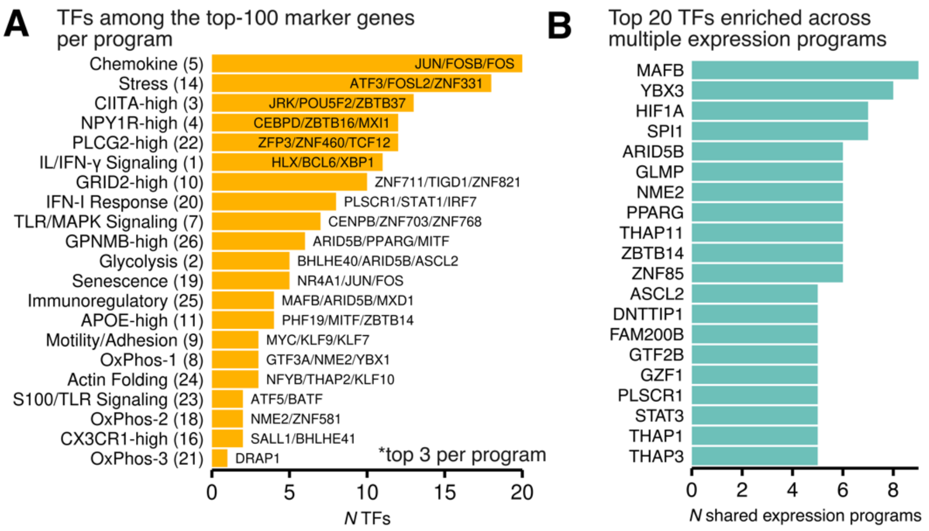

**Figure S21. Distribution of ARACNE-predicted TF regulons across scHPF factors.**
**(A)** Barplot showing the number of TFs among the top loading 100 genes across factors. **(B)** Barplot showing TFs that share enrichment of targets across the number of factors.

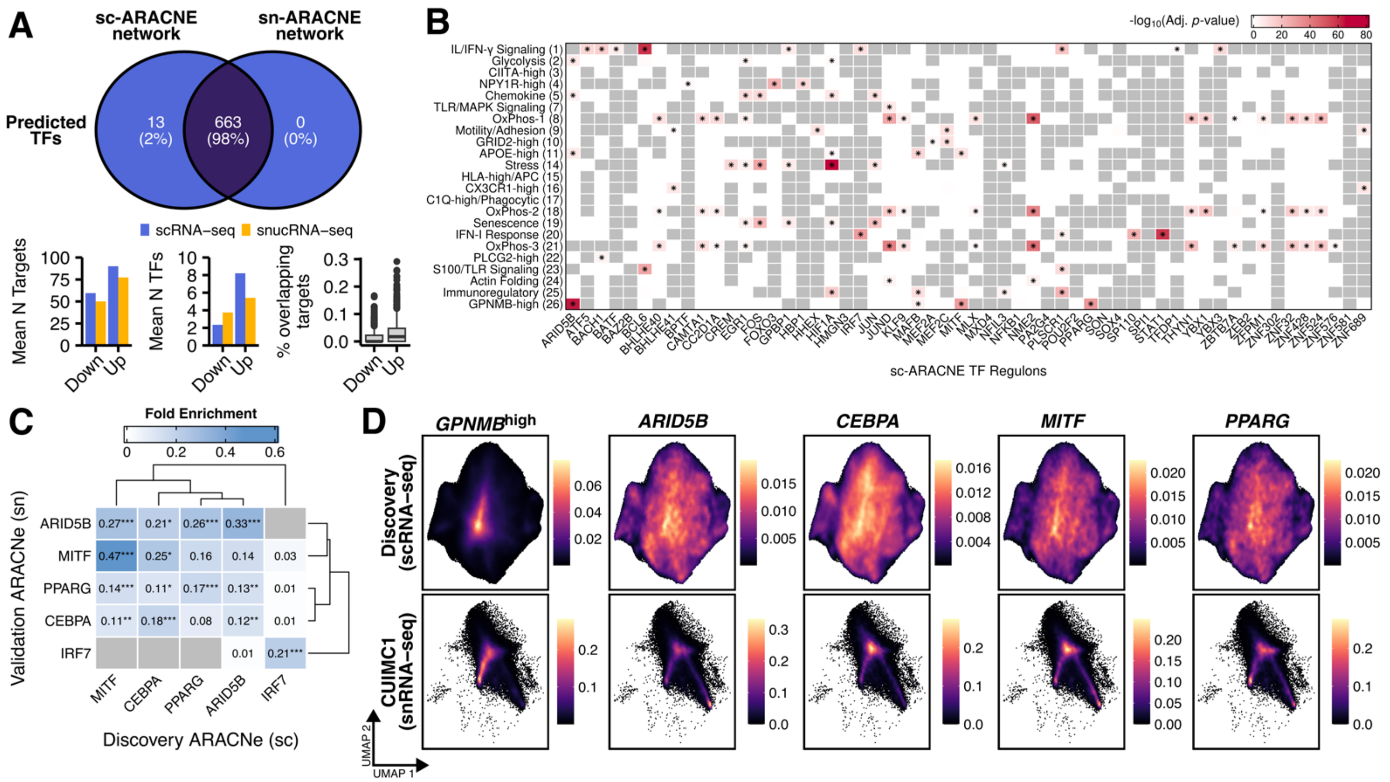

**Figure S22. ARACNe regulatory network in ROS-MAP CUIMC1 cohort.**
**(A)** Comparison of ARACNe networks independently-derived from single-cell vs. single-nucleus data, including predicted up- and down-regulated targets (see also **Table S11**). **(B)** Heatmap showing -$\log_{10}$ FDR $q$-values for hypergeometric enrichment of the top-100 loaded genes for each factor across single-cell prioritized ARACNe regulons. Grey indicates failed tests due to too few overlapping genes (see also **Table S44**). (**C**) Proportion of shared targets between single-cell regulons and single-nucleus regulons. Showing regulons prioritized for GPNMB$^{High}$ (scHPF_26) factor (i.e., ARID5B, CEBPA, MITF, PPARG) and IFN-I response (scHPF_20) (i.e., IRF7). Color indicates the proportion of shared targets calculated as the number of shared targets divided by the number of unique genes between the single-cell and single-nucleus regulon. Significance showing FDR-corrected $p$-value < 0.05 on a hypergeometric test. **(D)** Reference UMAP space showing the estimated density of microglial signatures, including the GPNMB$^{High}$ (scHPF_26) factor, individuals TFs (ARID5B, PPARG, MITF, CEBPA) and the GPNMB$^{High}$ (scHPF_26) factor.

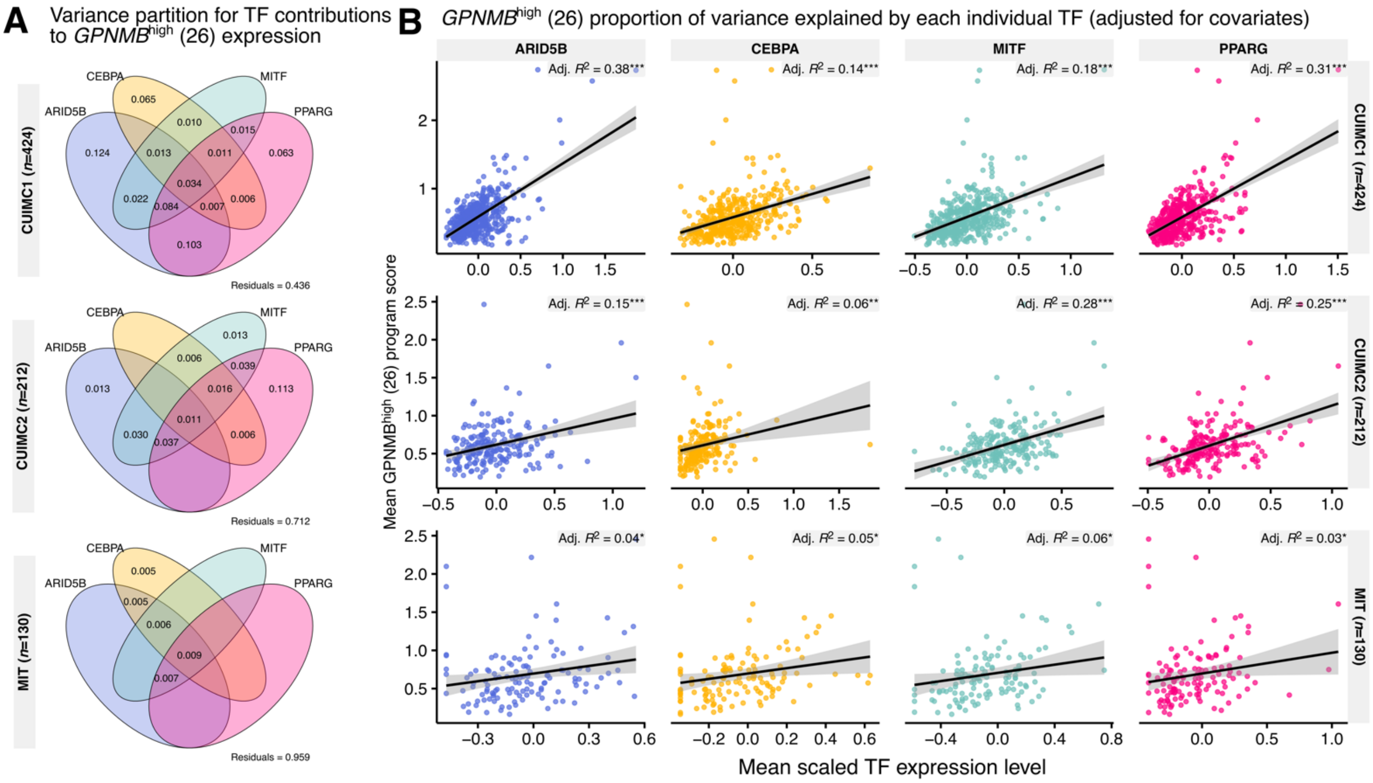

**Figure S23. Partitioning for individual TF contributions to variation in GPNMB$^{High}$ (26) scores.**
**(A)** Variance explained in donor-level *GPNMB*$^{High}$ (scHPF_26) factor score by TFs, unadjusted for technical and demographic variables. Values < 0.05% are not shown. Residuals denote the proportion of variance that remains unexplained by the TFs. **(B)** Variance explained in GPNMB$^{High}$ (scHPF_26) factor score by individual TFs. Showing donor-level mean-aggregated factor scores and TF levels. Adjusted $R^2$ showing the effect of the regulator, adjusted for technical and demographic variables (age at death, sex, PMI, study), but not the other TFs. Significance testing using permutation test for constrained correspondence analysis (10,000 permutations).

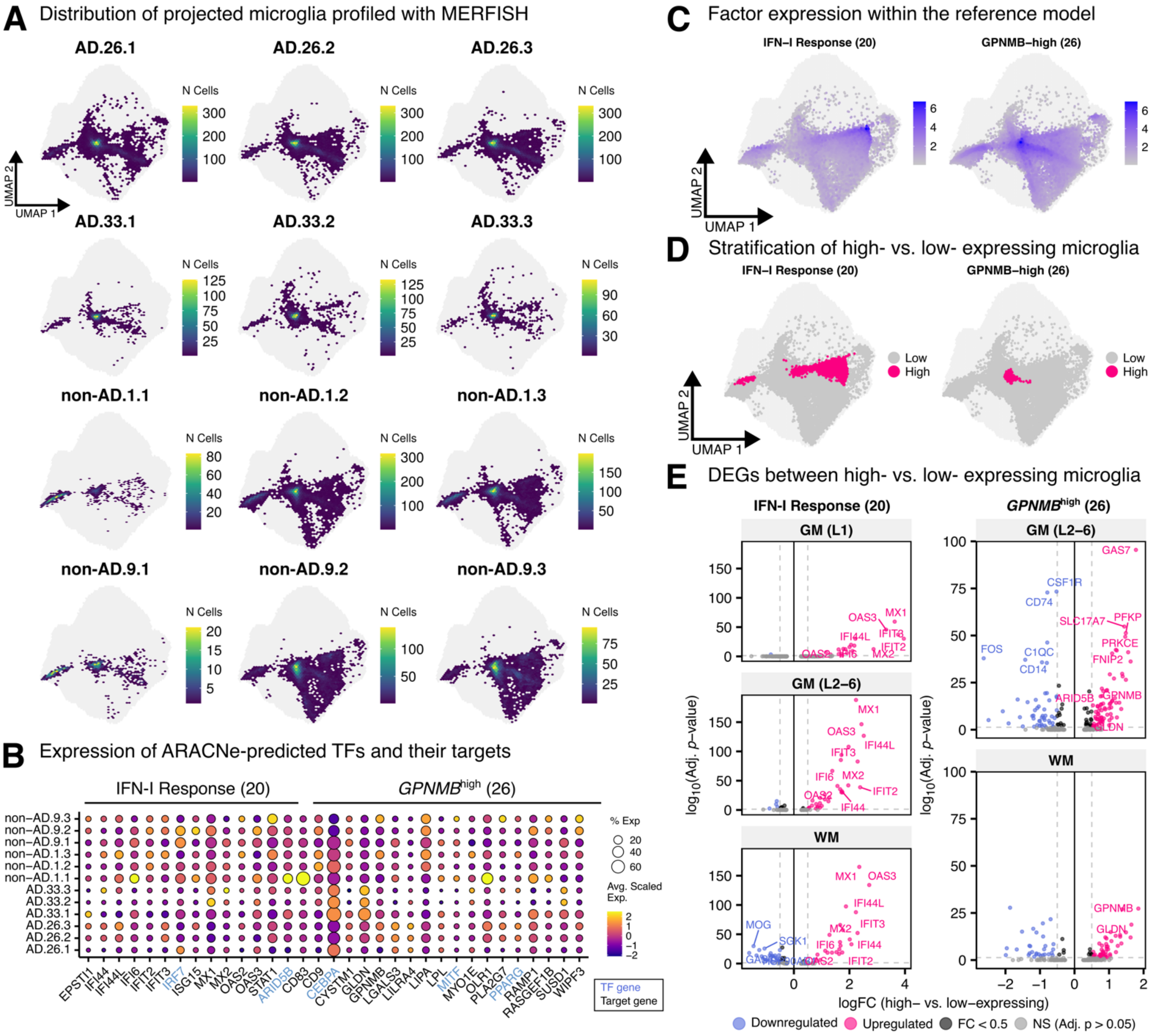

**Figure S24. IFN-I Response (scHPF_20) and *GPNMB*$^{High}$ (scHPF_26) factors can be identified *in situ*.** **(A)** UMAP embedding showing projected microglial transcriptomes from 12 tissues profiled using MERFISH (*n*=50,391 nuclei. Cells represented as binned, hexagonal 'meta-cells', colored by the number of transcriptomes within a hexagonal bin. **(B)** UMAP space showing the distribution of factor scores. **(C)** Dot plot showing the expression of marker genes for IFN-I response (scHPF_20) and *GPNMB*$^{High}$ (scHPF_26) factors, as well as ARACNe-predicted TFs, *IRF7*, *ARID5B*, *CEBPA*, *MITF*, and *PPARG*. **(D)** UMAP space showing the distribution of cells defined as high- vs. low expressing. *GPNMB*$^{High}$ high-expressing microglia represent 2.42% of the microglial population, while IR$^{High}$ microglia represent 4.77% of the microglial population. **(E)** DEGs between high- vs. low-expressing cells per factor based on a MAST model adjusted for UMI count and the random effect of donor tissue (see **Table S45**).

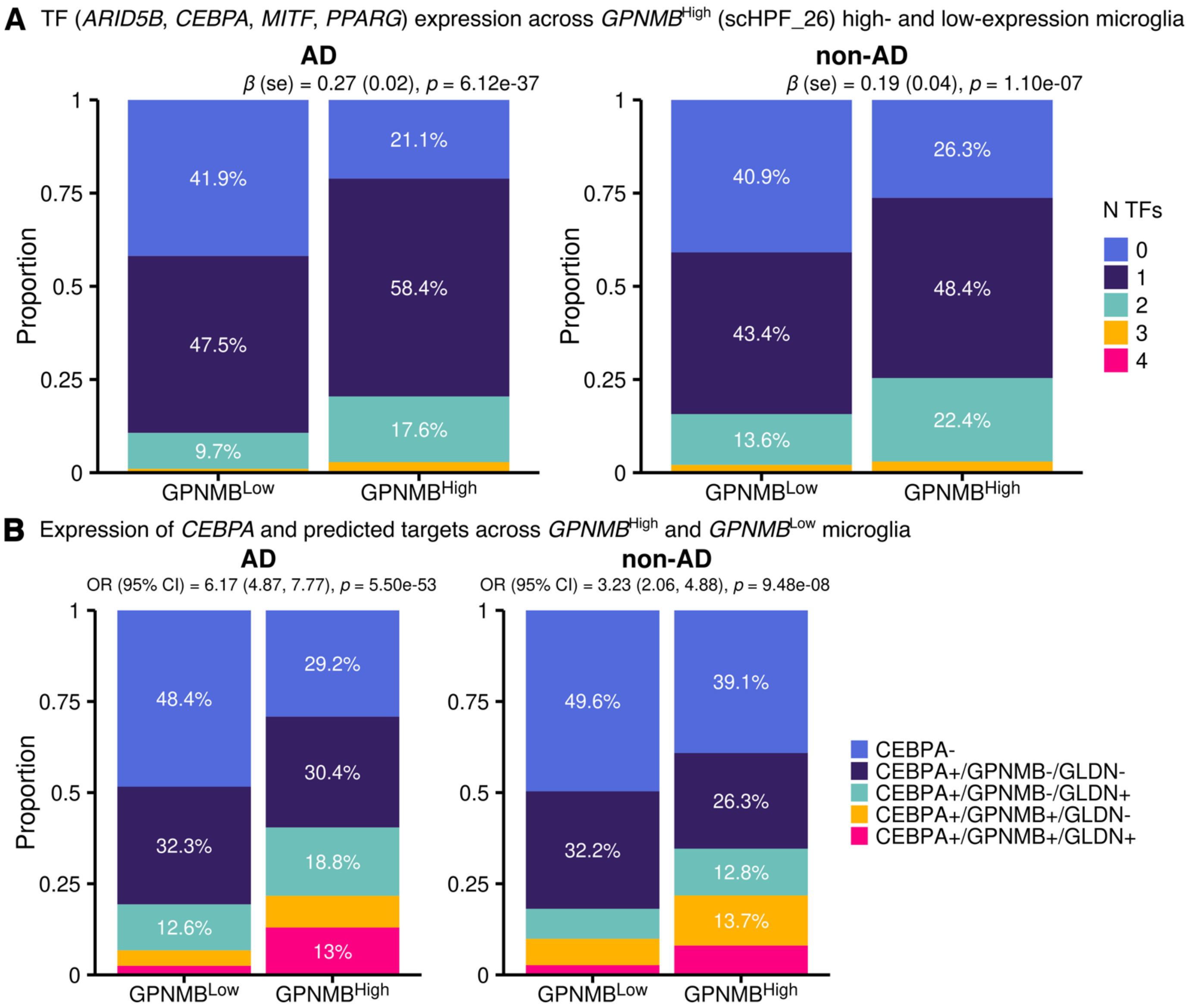

**Figure S25. TF and ARACNE-predicted target expression across *GPNMB*$^{High}$ and *GPNMB*$^{Low}$ microglia.**
**(A)** Barplot showing the proportion of *GPNMB*$^{High}$ vs. *GPNMB*$^{Low}$ microglia expressing total numbers of predicted TFs. Number of TFs is the count of TFs that have an expression level greater than 0 (LMM, adjusted for UMI count and the random effect of donor tissue). **(B)** Barplot showing the proportion of *GPNMB*$^{High}$ vs. *GPNMB*$^{Low}$ microglia that are triple positive for *CEBPA*, *GPNMB* and *GLDN* compared to all others (LMM, adjusted for UMI count and the random effect of donor tissue).

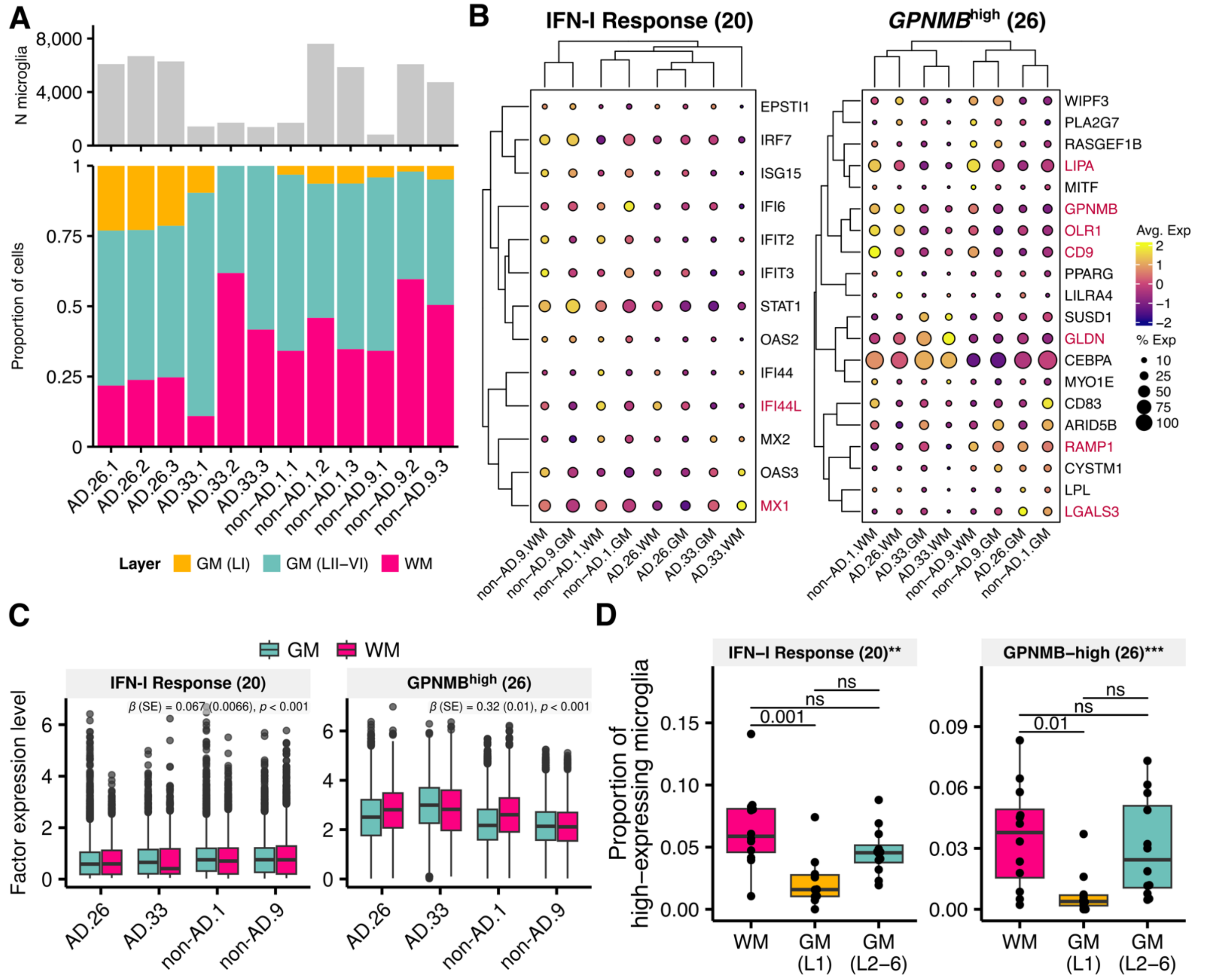

**Figure S26. Characterization of tissue layer niches and microglia factors.**

**(A)** Distribution of microglia across WM and GM niches per tissue sample. **(B)** Expression profiles across WM and GM niches based on marker genes for IFN-I response (scHPF_20) and *GPNMB*$^{\text{High}}$ (scHPF_26) factors. Red font face indicates genes that show differential expression across WM and GM niches (MAST adjusted for UMI count and diagnosis, as well as the random effect of donor). **(C)** Boxplots comparing factor scores between WM and GM across all tissues using a linear-mixed effects model adjusted for UMI count, diagnosis and the random effect of donor (FDR adjusted p-value). **(D)** Boxplot comparing the proportion of microglia highly expressing a factor of interest across tissue WM and GM niches.

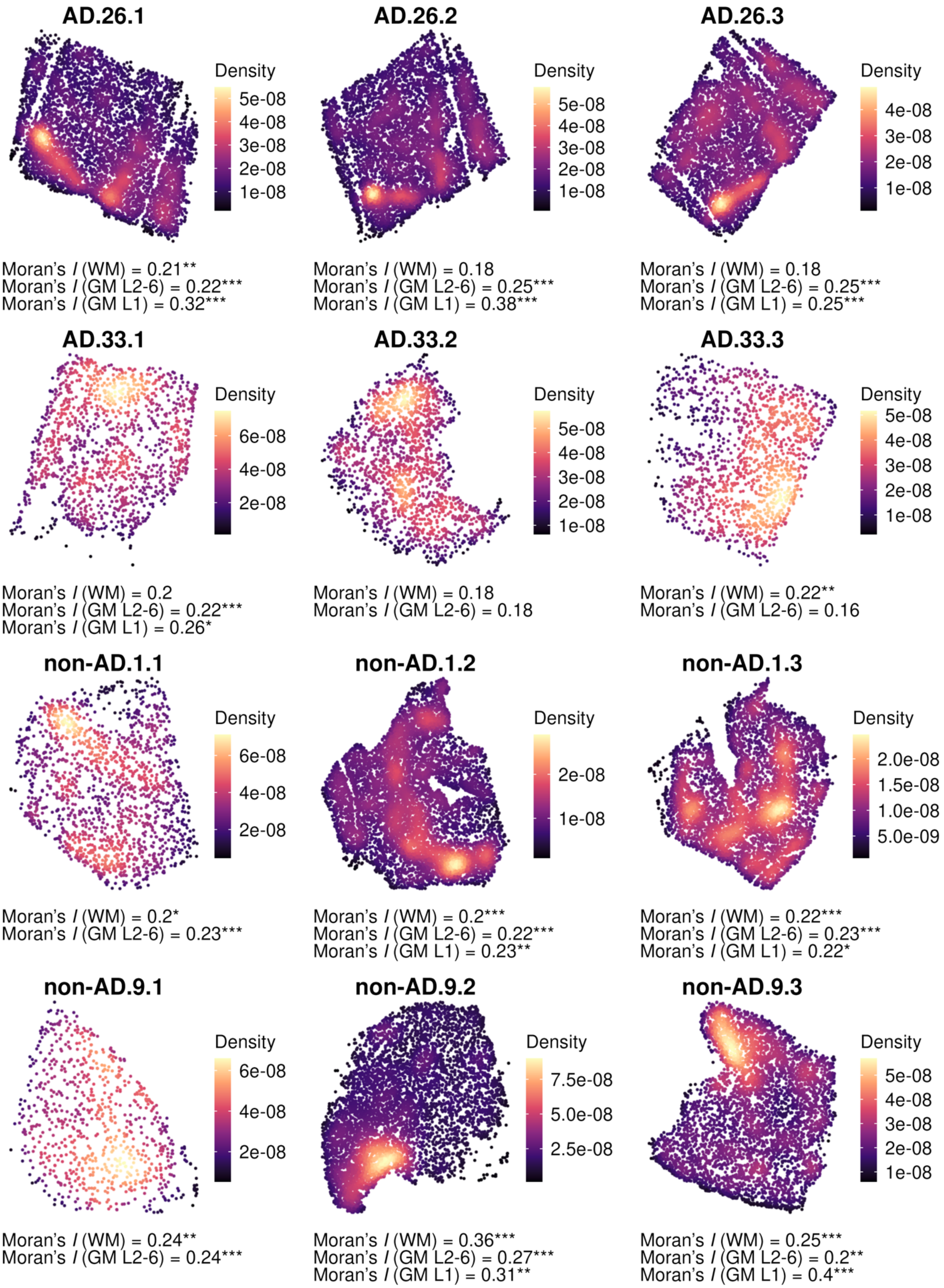

**Figure S27. Global (tissue-wide) spatial autocorrelation of IFN-I Response (scHPF_20) expression.** Color shows weighted kernel density estimation using *R* package Nebulosa[68] where lighter color indicates greater density of cells with higher factor scores compared to darker color across the twelve tissues profiled using MERFISH. Per tissue, layer-specific global Moran's I was calculated using R package Voyager. A higher global Moran I's indicates greater spatial autocorrelation. Note, in tissue with less than 100 microglia per layer, Moran's I was not calculated. Overall, the Moran's I was highest in GM L1 [M (SD) = 0.296 (0.068)], GM L2-6 [M (SD) = 0.22 (0.032)], and WM [M (SD) = 0.22 (0.049)]. *Significance levels*: *** $p < 0.001$, ** $p < 0.01$, * $p < 0.05$.

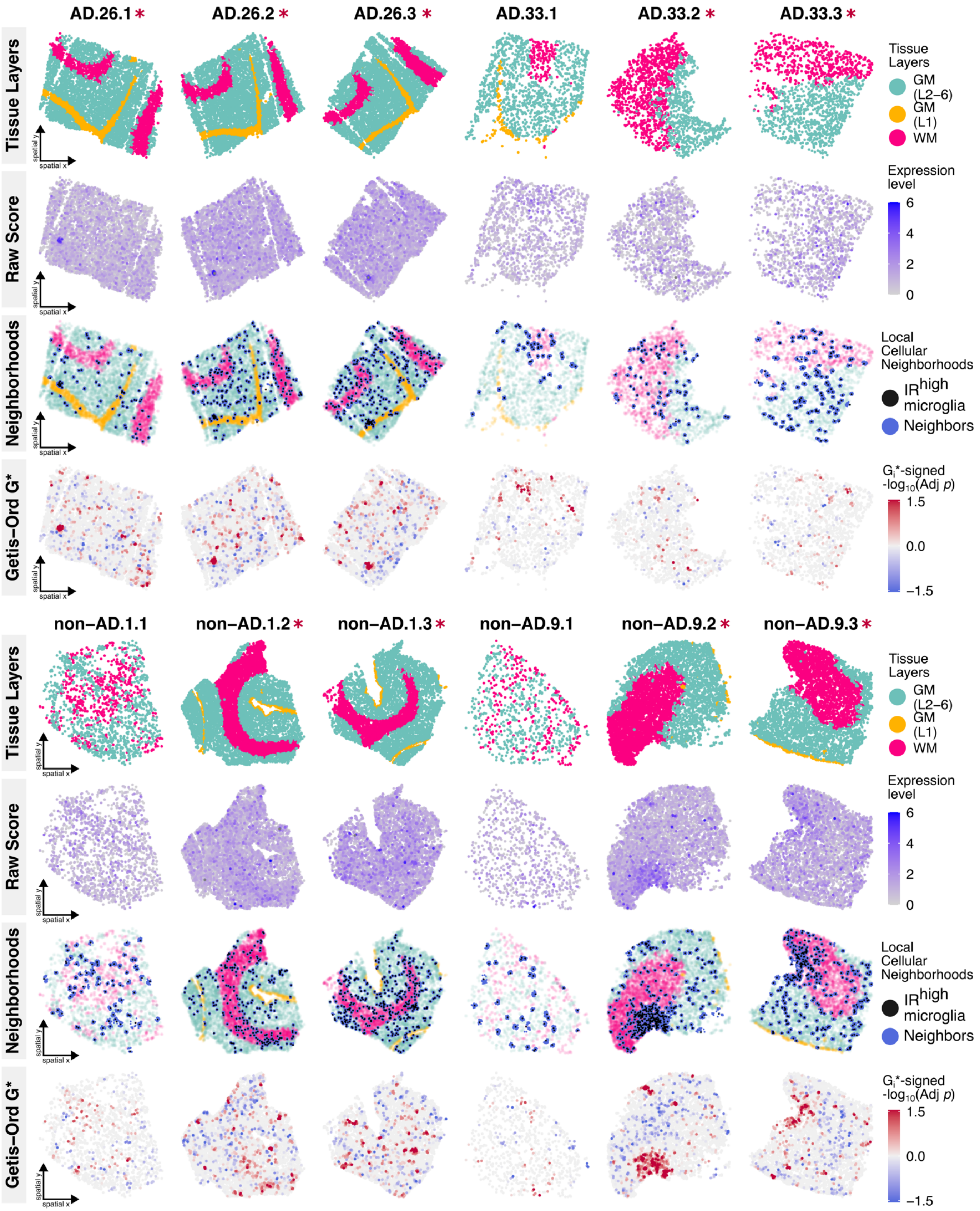

**Figure S28.** ***In situ*** **distribution of IR$^{High}$ local cellular neighborhoods.**
In situ plots showing the distribution of IFN-I Response (scHPF_20) expression levels and spatial scores across microglia for each of the twelve tissues. Row 1: Annotation of spatial layers for reference. Row 2: Raw scHPF_20 score per microglia. Row 3: Distribution of IR$^{High}$ microglia and any k-nearest neighbors. Row 4: Distribution of Getis-Ord $G_i$* score per microglia. Red asterisk denotes adjacent sections per donor (sister section or within 1-2 sections distance due to sectioning failure).

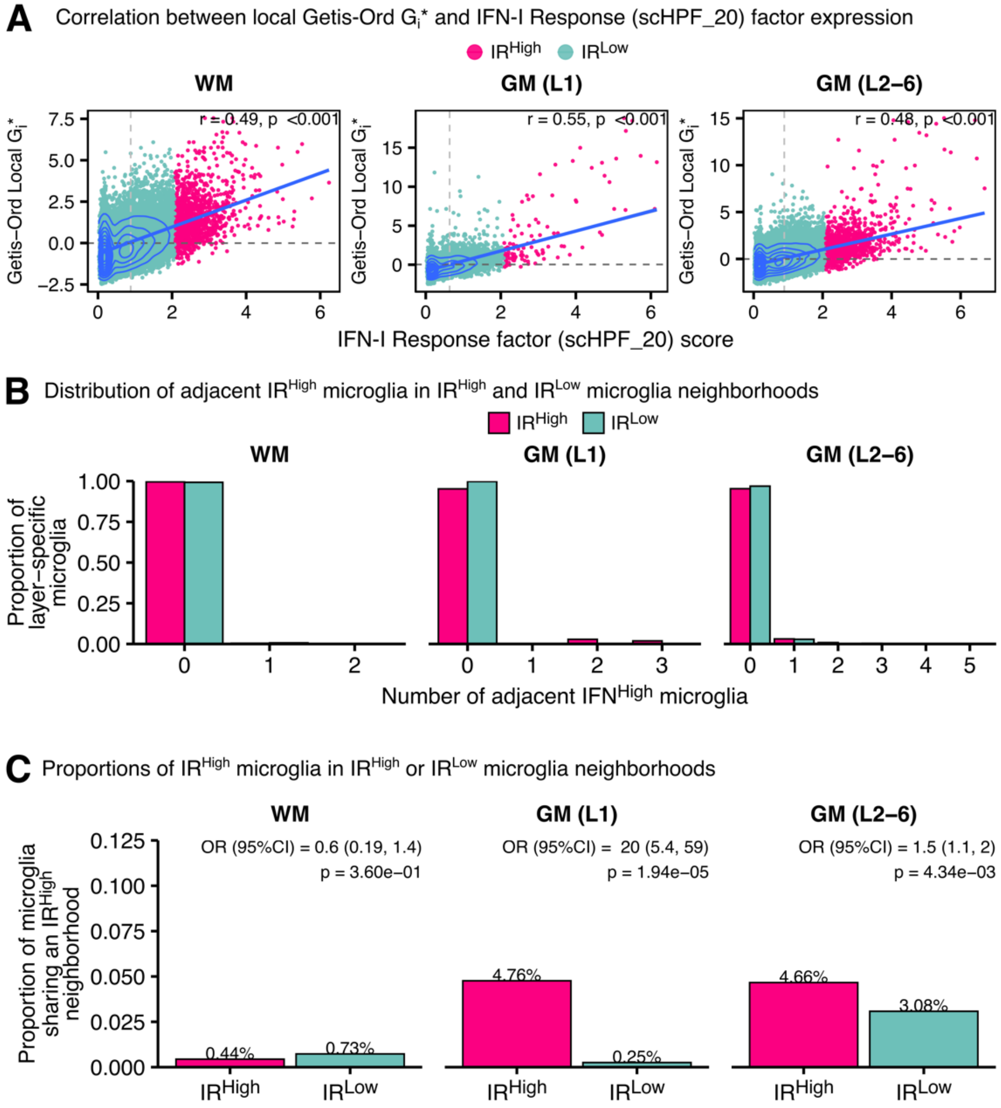

**Figure S29. Measures of local clustering for IR$^{High}$ factor (scHPF_20).**
**(A)** Scatterplot showing the positive correlation between IR$^{High}$ factor (scHPF_20) score and Getis-Ord Gi* score (local clustering; Pearson's correlation). **(B)** Distribution of IR$^{High}$ and IR$^{Low}$ microglia sharing neighborhoods with IR$^{High}$ microglia. **(C)** Proportion of IR$^{High}$ and IR$^{Low}$ microglia sharing neighborhoods with at least one IR$^{High}$ microglia (Fisher's exact test).

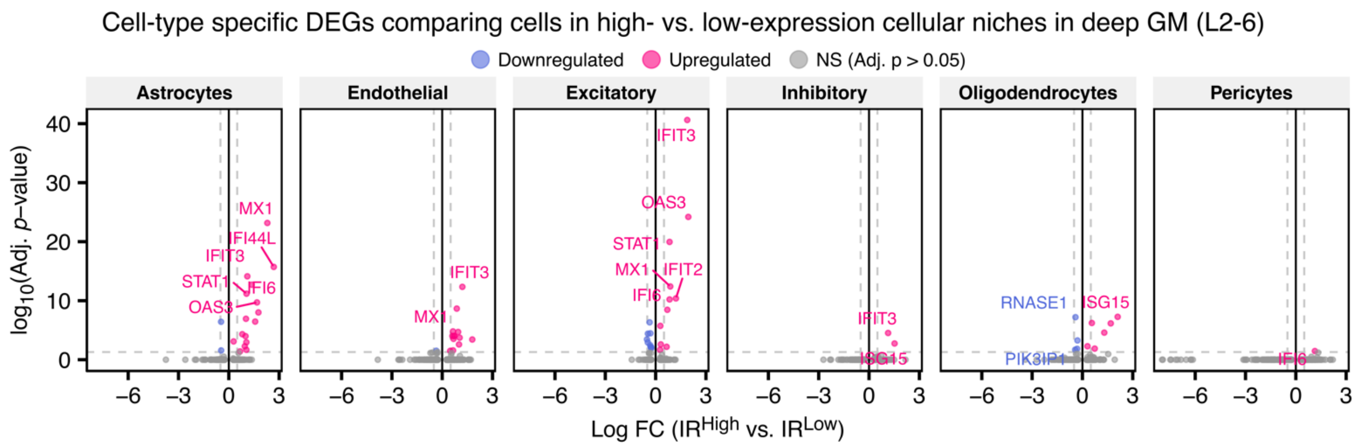

**Figure S30. DEGs across cell types in IR$^{High}$ and IR$^{Low}$ neighborhoods.**
Differentially expressed genes (DEGs) between cells in IR$^{High}$ and IR$^{Low}$ neighborhoods based on a MAST model adjusted for UMI count (see **Table S45**). Testing was only done for cell types with a minimum number of 50 cells across genes with a minimum logFC = 0.1.

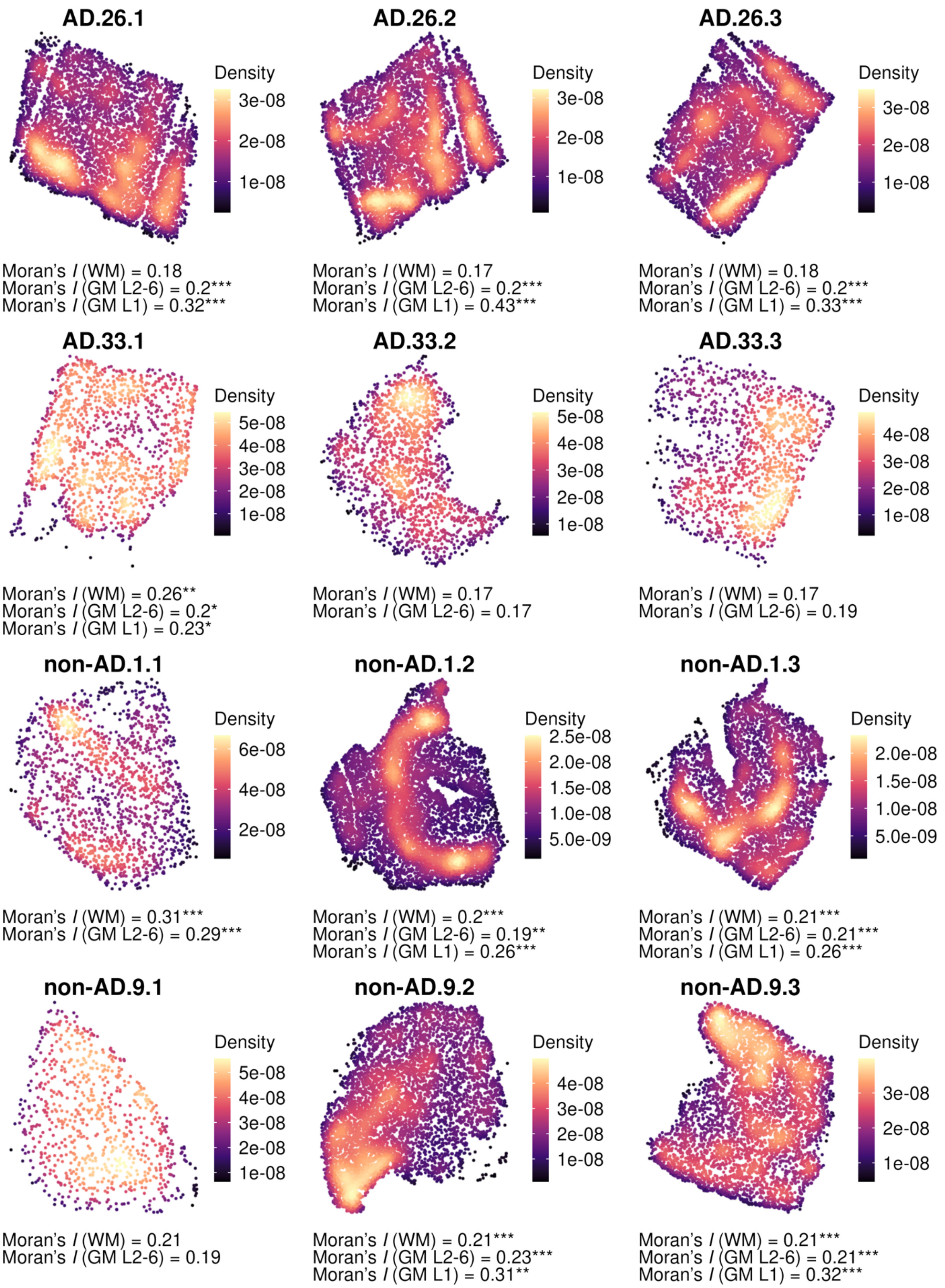

**Figure S31. Global (tissue-wide) spatial autocorrelation of *GPNMB*$^{High}$ (scHPF_26) expression.** Color shows weighted kernel density estimation using *R* package Nebulosa[68] where lighter color indicates greater density of cells with higher factor scores compared to darker color across the twelve tissues profiled using MERFISH. Per tissue, layer-specific global Moran's I was calculated using R package Voyager. A higher global Moran I's indicates greater spatial autocorrelation. Note, in tissue with less than 100 microglia per layer, Moran's I was not calculated. Overall, the Moran's I was highest in GM L1 [M (SD) = 0.31 (0.06)], GM L2-6 [M (SD) = 0.21 (0.031)], and WM [M (SD) = 0.21 (0.042)]. *Significance levels*: *** $p < 0.001$, ** $p < 0.01$, * $p < 0.05$.

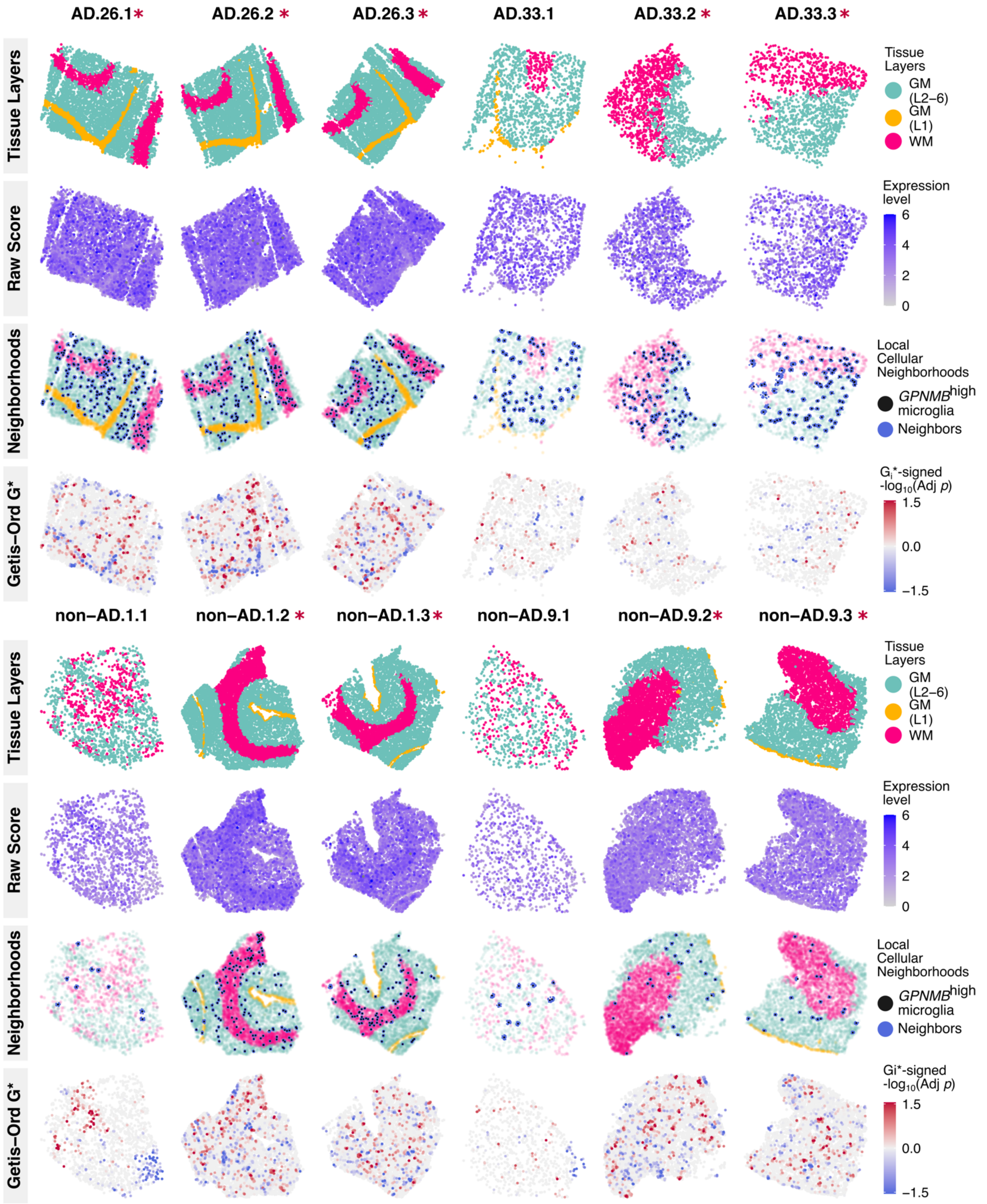

**Figure S32. In situ distribution of *GPNMB*$^{High}$ local cellular neighborhoods.**

In situ plots showing the distribution of *GPNMB*$^{High}$ (scHPF_26) expression levels and spatial scores across microglia for each of the twelve tissues. Row 1: Annotation of spatial layers for reference. Row 2: Raw scHPF_26 score per microglia. Row 3: Distribution of IR$^{High}$ microglia and any k-nearest neighbors. Row 4: Distribution of Getis-Ord $G_i$* score per microglia. Red asterisk denotes adjacent sections per donor (sister section or within 1-2 sections distance due to sectioning failure).

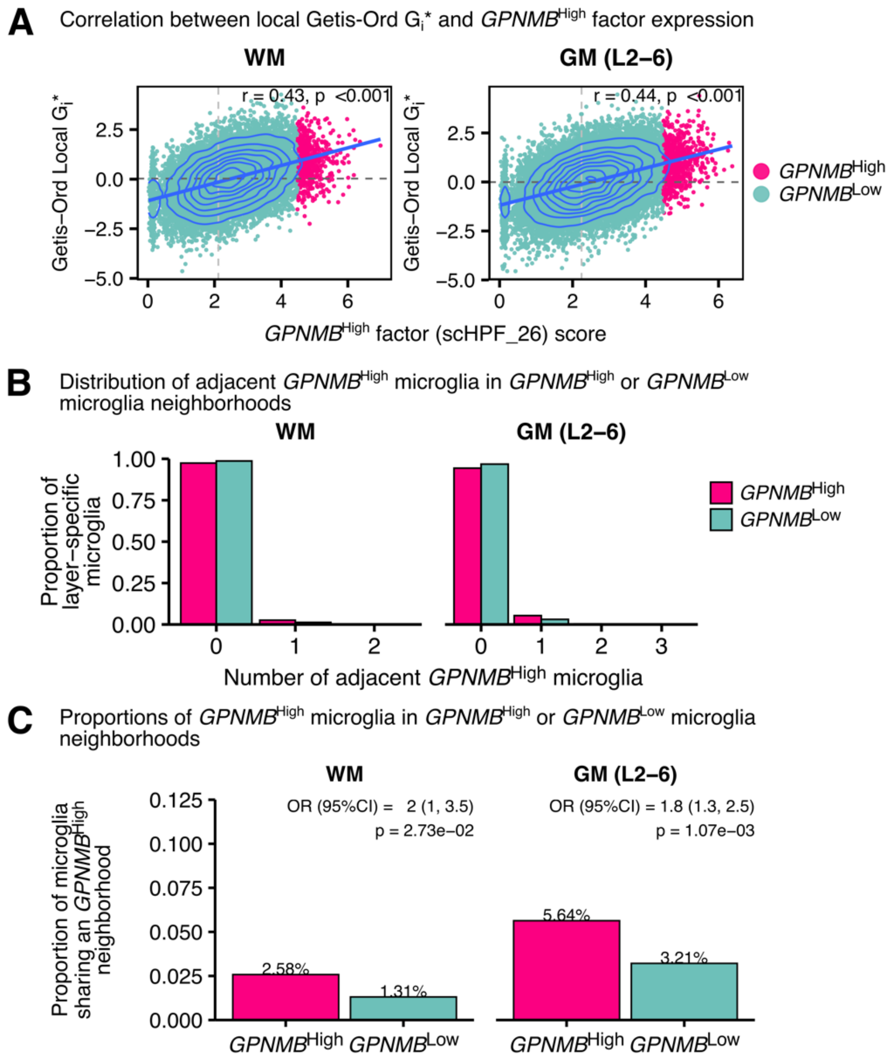

**Figure S33. Measures of local clustering for *GPNMB*$^{High}$ factor (scHPF_26).**
**(A)** Scatterplot showing the positive correlation between *GPNMB*$^{High}$ factor (scHPF_26) score and Getis-Ord Gi* score (local clustering; Pearson's correlation). *Note*: GM (L1) excluded from analysis due to low number of *GPNMB*$^{High}$ microglia (< 1%). **(B)** Distribution of *GPNMB*$^{High}$ and *GPNMB*$^{Low}$ microglia sharing neighborhoods with *GPNMB*$^{High}$ microglia. **(C)** Proportion of *GPNMB*$^{High}$ and *GPNMB*$^{Low}$ microglia sharing neighborhoods with at least one *GPNMB*$^{High}$ microglia (Fisher's exact test).

#### ***Supplementary Tables***

**Table S1.** Donor demographics and distribution of cells after quality control.
**Table S2.** Donor and tissue metadata.
**Table S3.** Individual gene loadings across the 23-factor scHPF model (8,478 genes).
**Table S4.** *Gene Ontology* enrichment across scHPF factors.
**Table S5.** Summary of external datasets projected onto the scHPF model.
**Table S6.** Milo differential abundance analysis for pathological AD in the ROS-MAP CUIMC1 cohort.
**Table S7.** Factor associations with AD-related neuropathology across ROS-MAP cohorts.
**Table S8.** Meta-analysis of factor associations with AD-related neuropathology in the ROS-MAP cohorts.
**Table S9.** Discovery ARACNe network.
**Table S10.** ARACNe regulon enrichment across scHPF factors (discovery network).
**Table S11.** Validation ARACNe network (CUIMC1 snRNA-seq)
**Table S12.** Enrichment of AD-related neuropathology across sc- and sn-ARACNE TF regulons.
**Table S13.** Vizgen MERSCOPE 412-gene panel.
**Table S14.** TF expression association with ARACNE-predicted target genes.
**Table S15.** Meta-analysis of TF expression association with ARACNE-predicted target genes.
**Table S16.** Cell-type specific DEGs comparing $IFN^{High}$ vs. $IFN^{Low}$ cellular niches *in situ*.
**Table S17.** MitoCarta pathway enrichment across scHPF factors.
**Table S18.** *Milo* differential abundance analysis across sexes in AD.
**Table S19.** *Milo* differential abundance analysis across age in AD.
**Table S20.** Association between age > 85 years and donor-level factor expression across brain regions.
**Table S21.** Meta-analysis of age association across brain regions at the donor-level.
**Table S22.** *Milo* differential abundance analysis in AWS compared to BA9/46 in AD.
**Table S23.** *Milo* differential abundance analysis in BA20/21 compared to BA9/46 in AD.
**Table S24.** *Milo* differential abundance analysis in the hippocampus compared to BA9/46 in AD.
**Table S25.** *Milo* differential abundance analysis with higher time-to tissue processing in the NYBB cohort.
**Table S26.** *Milo* differential abundance analysis with higher PMI in the discovery ROS-MAP cohort.
**Table S27.** Differential factor expression in mesenchymal versus classical glioma resection.
**Table S28.** Differential factor expression in drug-treated glioblastoma slice culture.
**Table S29.** Differential factor expression between 5xFAD-MITRG and WT-MITRG mice.
**Table S30.** *Milo* differential abundance analysis between 5X and WT xMGs.
**Table S31.** Factor enrichment across iMG clusters defined by Dolan et al., 2023.
**Table S32.** Differential factor expression across H1-derived iMGs treated with CNS substrates.
**Table S33.** Differential factor expression across AN-treated and untreated iPSC-derived iMGs.
**Table S34.** Differential factor expression in compound-treated HMC3 cells.
**Table S35.** Differential factor expression in a pooled CRISPR i/a screen of iTF-Microglia.
**Table S36.** Differential factor expression between KO and non-perturbed HMC3 cells.
**Table S37.** FGSEA enrichment with ROS-MAP CUIMC1 microglia subtypes.
**Table S38.** Factor associations with detailed neuropathology in the CUIMC1 cohort.
**Table S39.** Milo differential abundance analysis for high amyloid levels in the ROS-MAP CUIMC1 cohort.
**Table S40.** Milo differential abundance analysis for high tangle levels in the ROS-MAP CUIMC1 cohort.
**Table S41.** Factor associations with PMI across ROS-MAP cohorts.
**Table S42.** Meta-analysis of factor associations with PMI in the ROS-MAP cohorts.
**Table S43.** FGSEA enrichment of scHPF factor top genes across the sn-ARACNE regulons.
**Table S44.** ARACNe regulon enrichment across scHPF factors (validation network).
**Table S45.** Differential expression of genes between high- vs. low-expression microglia.
